# Sequence basis of transcription initiation in human genome

**DOI:** 10.1101/2023.06.27.546584

**Authors:** Kseniia Dudnyk, Chenlai Shi, Jian Zhou

## Abstract

Transcription initiation is an essential process for ensuring proper function of any gene, however, a unified understanding of sequence patterns and rules that determine transcription initiation sites in human genome remains elusive. By explaining transcription initiation at basepair resolution from sequence with a deep learning-inspired explainable modeling approach, here we show that simple rules can explain the vast majority of human promoters. We identified key sequence patterns that contribute to human promoter function, each activating transcription with a distinct position-specific effect curve that likely reflects its mechanism of promoting transcription initiation. Most of these position-specific effects have not been previously characterized, and we verified them using experimental perturbations of transcription factors and sequences. We revealed the sequence basis of bidirectional transcription at promoters and links between promoter selectivity and gene expression variation across cell types. Additionally, by analyzing 241 mammalian genomes and mouse transcription initiation site data, we showed that the sequence determinants are conserved across mammalian species. Taken together, we provide a unified model of the sequence basis of transcription initiation at the basepair level that is broadly applicable across mammalian species, and shed new light on basic questions related to promoter sequence and function.

## Introduction

Promoter is the key sequence element responsible for transcription initiation and the central hub in integrating transcriptional regulatory information. Over several decades, a handful of core promoter elements (or motifs) including the TATA-box, the Initiator (Inr) motif, and several downstream motifs (MTE, DPE, DPR) have been identified in various species(*1–3*). However, human promoters often possess none of these motifs(*3*). Moreover, while many transcription factor motifs appear near promoters(*3–5*), their roles in promoter function have not been clearly defined. For most human promoters we do not have the knowledge of which basepairs contribute to its activity.

Our understanding of how sequence patterns determine transcription start sites for the majority of human promoters thus remains incomplete(*1–3*). Moreover, the transcription initiation process involves many factors, and even a single base pair may be multi-functional(*1–3*, *6*), making this problem especially challenging. Hence, a systematic approach that simultaneously dissects multiple types of sequence dependencies is critical for solving the problem.

Therefore, basic questions, such as the following, remained open: What are the bases that contribute to any given promoter and determine transcriptional initiation patterns *at basepair resolution*? How do sequence patterns work together to determine transcription initiation sites? What is the impact of promoter sequence composition on its function? What are the key factors that determine the strand-specificity of promoters? And finally, how conserved are sequence determinants of transcription initiation across species? Developing a strategy that can be used to answer these questions would be transformative to our ability to analyze, predict, engineer and control transcription initiation.

To address the stated questions and overcome the limitations of current methodologies, we developed Puffin, a deep learning-inspired explainable model of transcription initiation at basepair resolution in the human genome. We showed that a few simple rules are sufficient to explain the sequence contribution underlying most promoter sequences. The model provided insights into mechanisms of identified sequence patterns by characterizing their position-specific effect curves for the first time, which were validated by experimental results. Puffin both recapitulated past findings and discovered new roles of known and unknown motifs, creating a new unified view of transcription initiation at the sequence level.

Beyond distilling the sequence rules of transcription initiation, we systematically characterized sequence contributions to transcriptional initiation sites at the basepair level and motif level. With this information, we further analyzed the relationship between motif contribution and gene expression regulation, explaining the connection between promoter sequence and cell type-specificity of the gene. The model also explained the sequence-basis of bidirectional transcription. Moreover, we demonstrated that the sequence dependencies captured by the model are highly conserved between mouse and human, as well as across 241 mammalian species from the Zoonomia project(*7*).

To facilitate interactive analysis of any promoter sequence, we have designed and built a user-friendly web server (https://tss.zhoulab.io) powered by Puffin which provides rich output to facilitate the understanding and manipulation of any promoter sequence of interest.

## Results

### Decoding sequence basis of transcription initiation at basepair-resolution

We hypothesized that basepair resolution transcription initiation signal patterns contain signatures of underlying sequence-based transcription initiation mechanisms. Therefore, capturing how transcription initiation patterns depend on sequence patterns may allow deconvolution of such mechanisms. Thus, we first assembled the highest coverage basepair resolution transcription initiation maps.

We integrated transcription initiation datasets from five techniques that precisely capture the 5’ end of transcripts: two variants of CAGE(*8*) and RAMPAGE(*9*) from the FANTOM and ENCODE projects that measure mostly mature transcripts and GRO/PRO-cap that measures only nascent transcripts. We aggregated all samples for each technique to obtain the most robust estimate at single-base resolution for the transcription initiation signal (Data S1). FANTOM CAGE (6.7 billion reads) and PRO-cap (2.3 billion reads) have the highest coverage within each group of techniques and were prioritized for presentation here (all analyses are consistent across techniques unless otherwise indicated).

To elucidate the sequence basis of transcription initiation, we developed the first basepair-resolution sequence models of transcription initiation signals (**Fig. 1A-C**), including a pair of complementary models: the performance-focused Puffin-D model, which is a deep learning model with a new architecture that predicts basepair-resolution signals from a long sequence context of 100kb (Fig. S1, S2), and the interpretation-focused model **Puffin**, the focus of this manuscript, which extracted a simple and compact set of sequence rules for transcription initiation at basepair resolution. The Puffin model is designed based on analyses of sequence dependencies captured by the deep learning model Puffin-D (Supplementary Text 1).

**Fig. 1.**
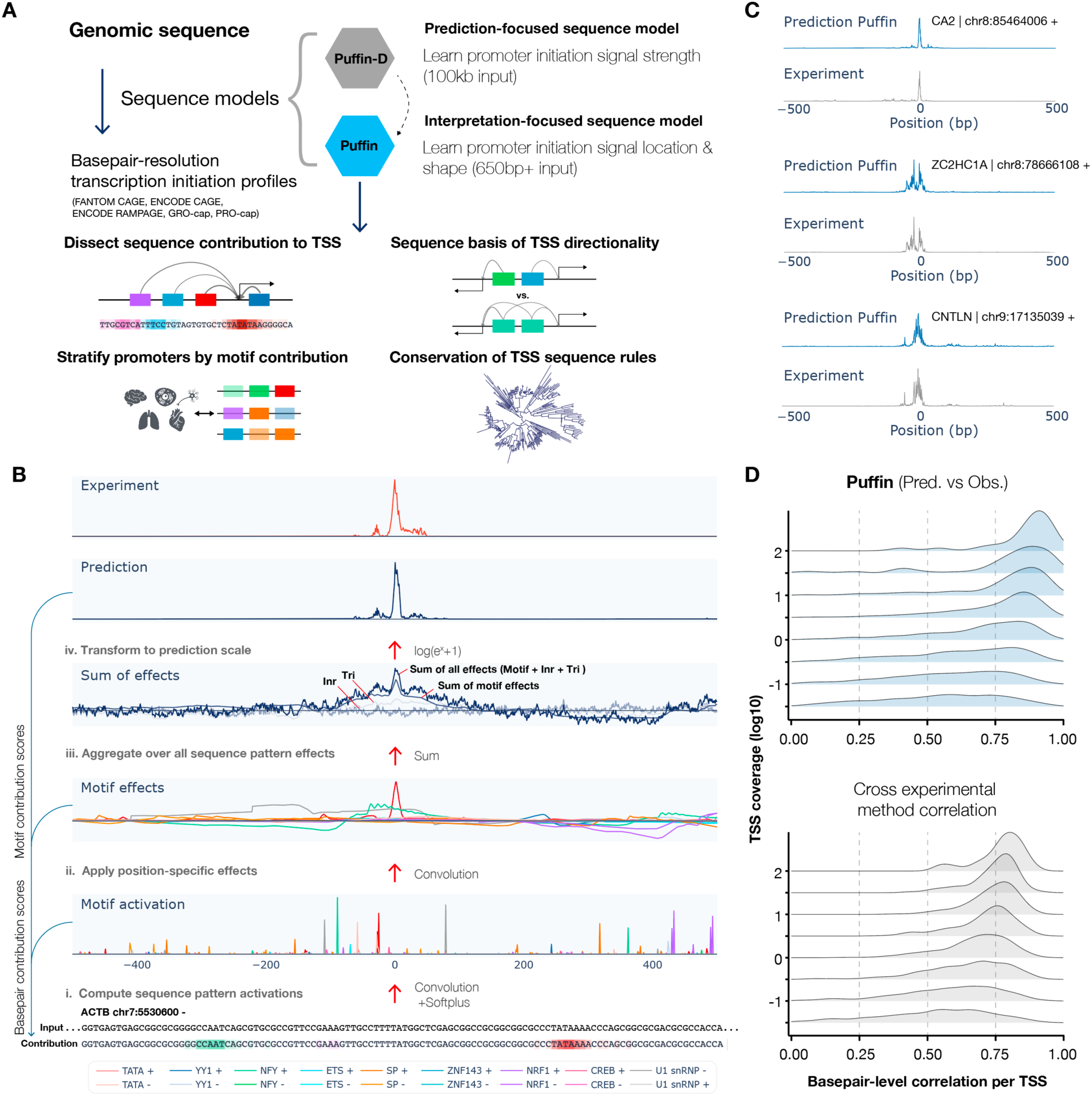
Dissect the sequence basis of transcription initiation with sequence models. (**A**) Schematic overview of sequence-based models of transcription initiation. The prediction-focused Puffin-D and interpretation-focused Puffin models were trained to predict basepair-resolution transcription initiation signals from sequence. These sequence models enabled analyses of transcription start site (TSS) motif composition, directionality, regulatory properties, and sequence rule conservation. (**B**) Step-by-step illustration of the Puffin promoter sequence model. The input sequence is first converted to activation scores of learned sequence patterns. Sequence pattern effects are computed next based on the activations. The motif effects are then summed together with initiator and trinucleotide sequence pattern effects, and transformed into the prediction. The computation of motif contribution scores and basepair contribution scores are illustrated on the left. The motif name legend is shown at the bottom. (**C**) Example prediction of basepair resolution transcription initiation signal from promoter sequences on holdout chromosomes. The x-axis indicates the position relative to the annotated transcription start site and the y-axis is shown in log10 scale. (**D**) Basepair-level correlation (x-axis) between Puffin (top panel) and experimental measurement (FANTOM CAGE) within 1kb window of each annotated TSS. TSS were grouped by coverage level with the y-axis indicating the lower bound of each group (the upper bound of the group is the lower bound of the next group). The bottom panel shows the correlations between ENCODE CAGE and FANTOM CAGE, the most correlated pair of techniques, which also provides a reference for the expected decrease in correlation due to lower coverage.

Importantly, Puffin shows that a simple set of rules can explain the majority of human promoter sequences (**Fig. 1D**). Trained to predict transcription initiation signal shape, Puffin achieves higher basepair-level correlation with experimental measurement than the most correlated pair of experimental techniques, when evaluated on test chromosomes (**Fig. 1C-D**, Fig. S3). To focus on promoter sequence dependencies, Puffin uses only proximal sequence context and focuses on the shape but not the scale of the transcription initiation signal.

Puffin captured the sequence dependencies of transcription initiation with two major steps of computation. First, it computes basepair-resolution *activation scores* for all sequence patterns it learned (*activation score is analogous to motif match score with non-matches set to zero, Methods)*; next, all sequence pattern activations’ position-specific *effects* on transcription initiation are computed and combined additively in log scale (which is equivalent to multiplicative combination in count scale) (Methods, **Fig. 1B**).

With a data-driven design (Supplementary Text 1), the model learns three types of sequence patterns to capture different types of sequence dependencies (**Fig. 2A-C**, Fig. S4): 1. *Motifs,* the main sequence drivers of transcription initiation site and shape; 2. *Initiators*, which tune the local basepair-level preference to transcription initiation *within* the initiator sequence patterns; 3. *Trinucleotides*, which capture the residual local sequence dependencies. Thus, with a small number of sequence patterns (10 motifs + 10 initiators + all trinucleotides) and a simple additive/multiplicative rule, we can for the first time predict basepair resolution transcription initiation signals from sequence in strong agreement with experiments.

**Fig. 2.**
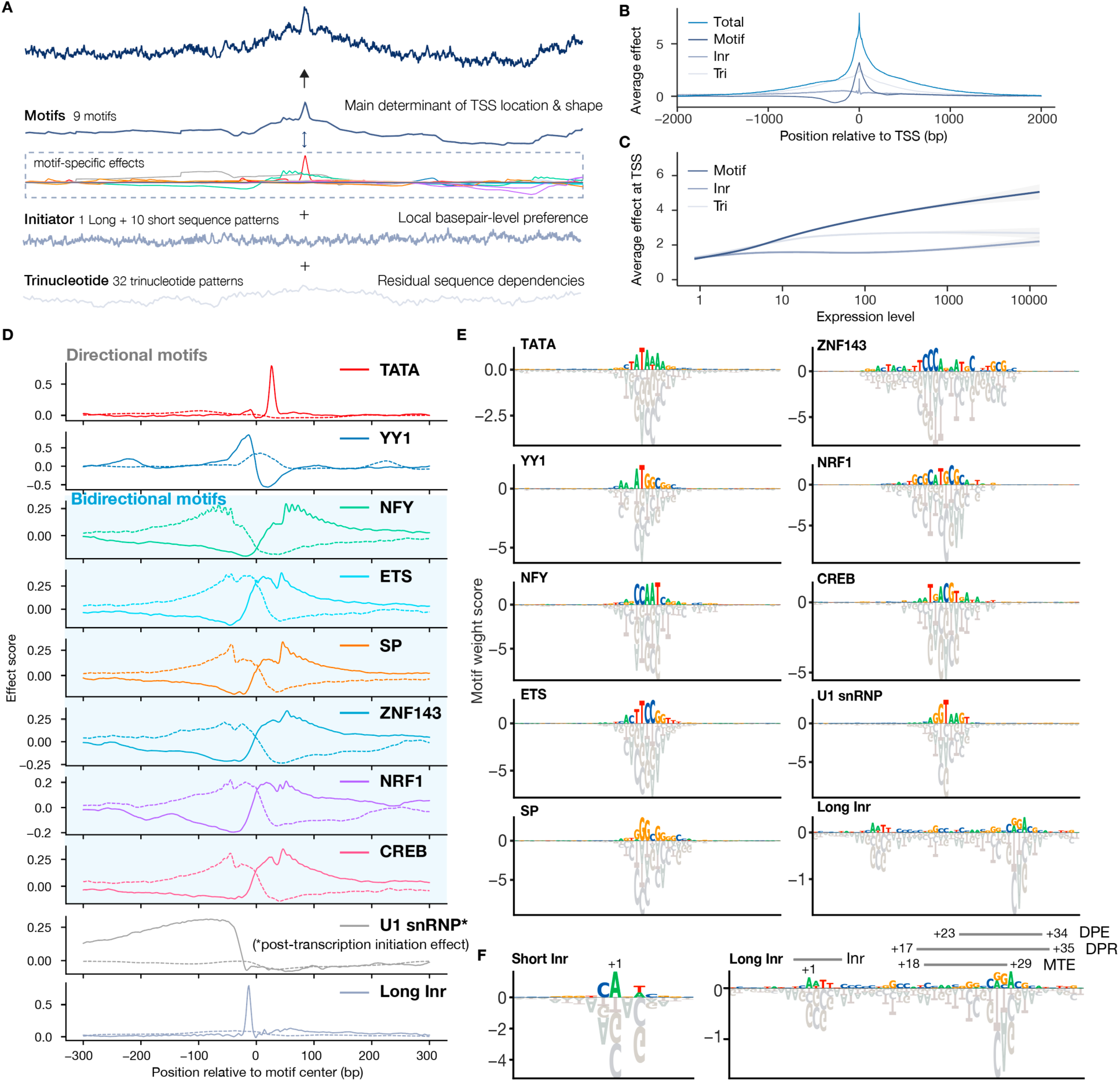
Sequence patterns with position-specific effects on transcription initiation. (**A**) Overview of the three sequence pattern types that the Puffin model learns. (**B**). The average position-specific effect of each sequence pattern type over the top 40,000 TSS by FANTOM CAGE signal (top panel). (**C**) The average sequence pattern effect at TSS (y-axis) varies with TSS expression levels (x-axis). Generalized additive model fitted curves and 95% confidence intervals are shown. (**D**) Position-specific effects of all motifs. The bidirectional motifs are indicated with blue background and all other motifs are direction-specific. The dotted lines show the motif effect on reverse strand transcription initiation. U1 snRNP effects are post-transcription initiation effects. (**E-F**). All transcription initiation motifs learned by the Puffin model with assigned names. Known promoter motifs overlapping with the Long Inr motif are indicated in (**F**). The height of each base represents the motif score (convolution kernel weight) for that nucleotide.

### Position- and strand-specific sequence effects on transcription initiation

The core of the transcription initiation sequence model is the position-specific effect curves of sequence patterns. The model has learned a distinct curve for each motif that represents the activation and repression effects of the motif at different locations relative to the motif (**Fig. 2D**). Each position-specific effect curve is a transcriptional signature of the motif and likely reflects its mechanism of action.

All ten motifs included in the Puffin model are robustly discovered with >0.95 correlation across training replicates (**Fig. 2D-E**, Supplementary Data 2), which we call individually identifiable sequence patterns. While all motifs were learned from scratch using only sequence and transcription initiation signal data, many of them match known motifs(*10*). However, their position-specific effect curves on transcription initiation have never been characterized. Thus, the position-specific motif effects were characterized by Puffin for the first time. Importantly, these effects are well supported by experimental perturbations in the next section and by evolutionary conservation in the last section.

We assigned each motif a name and an ID (Table S1). By symmetry of motif effects, the motifs can be divided into two groups: a group of strand-specific, or directional motifs that have strong effects on the forward strand and much weaker or no effect on the reverse strand: TATA, YY1, U1 snRNP, and Long Initiator (Long Inr) (**Fig. 2D-F**), and a group of non-strand-specific, or bidirectional motifs with almost symmetrical effects on both strands: SP, NFY, ETS, ZNF143, NRF1, and CREB (**Fig. 2D-F**, Fig. S5). Below, we summarize each group of motifs separately, as well as initiator sequence patterns and trinucleotide patterns.

#### Direction-specific promoter motifs

Puffin identified the well-established TATA motif as expected, and it also has the most position-specific and strand-specific effects in Puffin. The YY1 motif is also estimated to be highly strand-specific. The YY1 motif has one of the most distinctive transcription initiation effect patterns estimated by Puffin, activating transcription at the immediate upstream and ∼200bp upstream of the first peak (**Fig. 2D**). YY1 is known to bind promoter sequence in vitro and a role in promoter has been suggested(*5*, *11*), but its position-specific effect on transcription initiation has not been previously resolved. The U1 snRNP motif (or the 5’ splice site motif) was the only motif that exhibited a strong estimated positive effect on CAGE/RAMPAGE data, experiments that detect total mRNA, but not on PRO/GRO-cap data, where only nascent transcript are detected. This suggests that U1 snRNP motif exerts its impact after transcriptional initiation, consistent with recent findings that it promotes transcription elongation(*12–15*). The last direction-specific motif, the Long Initiator (Long Inr), had more in common with the initiator sequence patterns and is described in more detail below. Direction-specific motifs are thus expected to contribute to the strand preference of transcription initiation sites.

#### Bidirectional promoter motifs

The bidirectional group motifs are expected to bind the trimeric NF-Y TF (NFY), KLF/SP family TFs (SP), ETS family TFs (ETS), the zinc finger TF ZNF143 (ZNF143), CREB/ATF family TFs (CREB), and the homodimeric TF NRF1 (NRF1) respectively. Interestingly, while each motif in the bidirectional group has a highly distinguishable position-specific effect pattern, they also share apparent similarities in position-specific effect profiles, which may indicate similarities in their transcription activation mechanisms (**Fig. 2D**). The NFY motif showed the most distinctive position-specific effect pattern with ∼10.5bp periodicity. 10.5bp matches the helical periodicity of the DNA double-strand and may indicate a more rigid physical interaction with the Pol II preinitiation complex. Similar, albeit weaker 10.5bp periodicity was also observed in the position-specific effect curves of other motifs. The CG-rich SP motif was the most common motif at promoters (Fig. S6). The NRF1 motif was the only palindromic motif among all motifs with symmetric effects. As we will discuss later, bidirectional motifs are likely the basis of the bidirectional transcription initiation at most human TSS. Moreover, all bidirectional motifs activate transcription away from the motif on both strands, avoiding the formation of double-stranded RNA.

#### Initiator patterns tune the local propensity for transcription initiation

The initiator sequence patterns are named after the initiator (Inr) element(*16*), one of the first core promoter elements identified. Initiator sequence patterns in Puffin are expected to tune the local propensity for the exact nucleotide(s) that transcription starts from *within* the sequence pattern itself (**Fig. 2A-B**). We refer to the initiator sequence pattern that matches the Inr as Short Inr (**Fig. 2F)**, because Puffin also identified a new related motif Long Inr, which is an extended Short Inr motif that contains several downstream core promoter elements including MTE, DPE, and DPR (**Fig. 2F**). Similar to these motifs, Long Inr may represent the nucleotide preferences of TFIID, which binds to the core promoter to position the Pol II. In addition, initiator sequence dependencies are more complex than Long and Short Inr, as other initiator sequence patterns we identified were able to explain about half of the variance of initiators. The sum of initiator effects are also highly reproducible across training replicates (Fig. S7).

#### Trinucleotide patterns capture residual local sequence dependencies

Residual local sequence dependencies are captured by the trinucleotide patterns, (Methods, Fig. S8-10, Supplementary Text 2). Both individual and total trinucleotide effects are highly reproducible across training replicates (Fig. S7,11-12). The trinucleotide patterns with the strongest contribution to TSS are CpG-containing patterns (Fig. S8). The effects of CpG islands are likely explained by both CG-rich motifs like SP and trinucleotide effects in Puffin.

As shown by the average effect of each sequence pattern type (**Fig. 2B**), motifs, initiators, and trinucleotides define transcription start site at three spatial scales. Motifs are the most important contributor to the TSS activity, and their effect ranges are the longest. Moreover, genes with higher expression are characterized by stronger contributions from motifs (**Fig. 2C**). Initiators tune the local basepair-level preference of transcription initiation based on local sequences and mostly only affect the exact base of transcription initiation. While trinucleotides capture mostly local sequence dependencies within 50bp, a broad region of a few kilobases near TSS is enriched in trinucleotide patterns preferable for transcription initiation (**Fig 2B**, Fig. S8-10). Overall, motifs, initiators, and trinucleotides capture different aspects of sequence dependencies, which together explain basepair resolution transcription initiation in most human promoters. Next, we focus on motifs and their roles in transcription initiation, gene regulation, and evolution.

### Experimental perturbations validated position-specific motif effects on transcription initiation

To directly validate the motif effects with independent data, we analyzed experimental data of TF depletion effects on basepair-resolution transcription initiation patterns and compared them with model predictions (**Fig. 3A-C**). Puffin was developed to have the capability of performing in silico TF knock-out in the model to predict the effects of depleting a specific TF, by turning off the activation and effects of the corresponding motif (**Fig. 3A**).

**Fig. 3.**
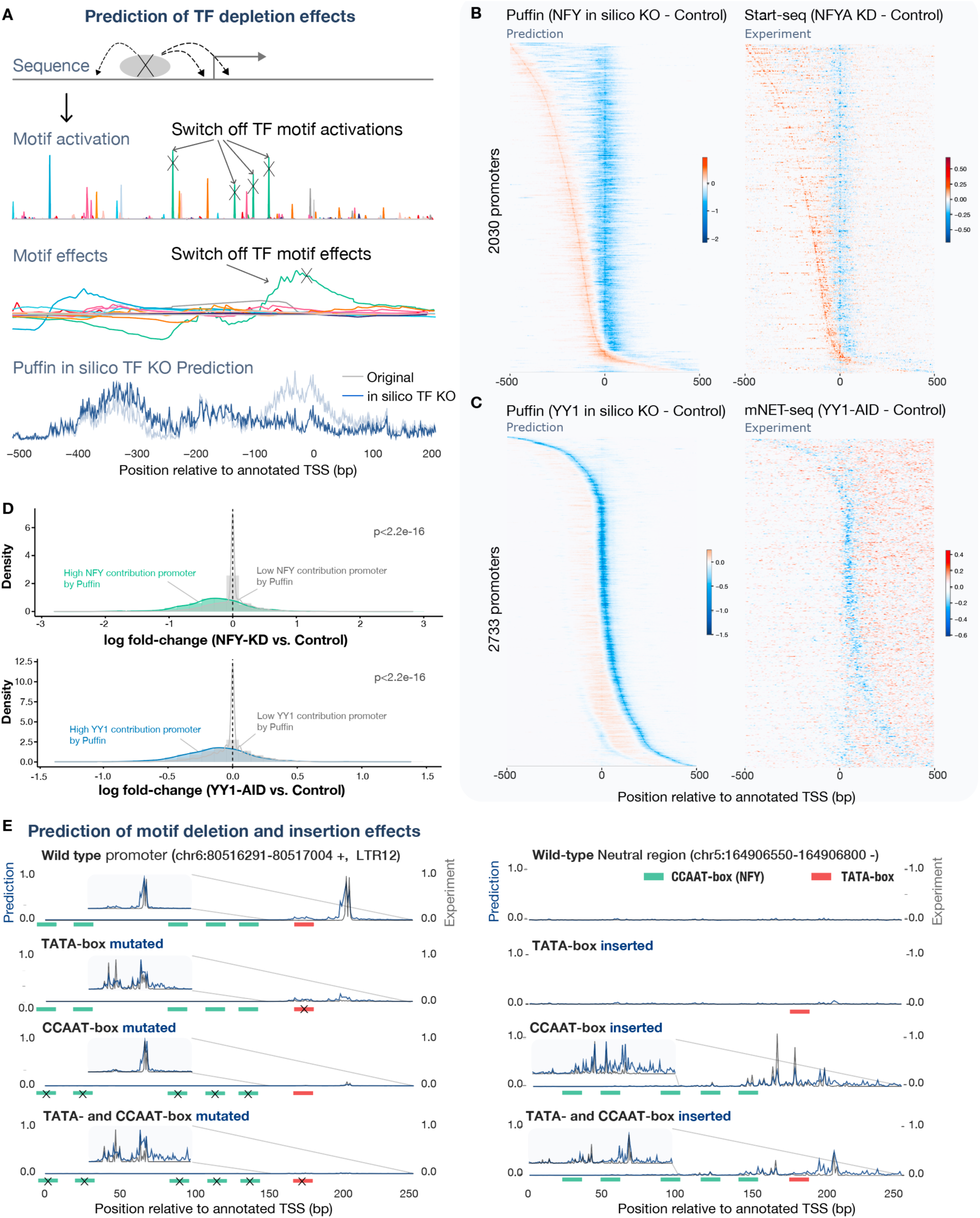
Experimental TF and motif perturbation effects are recapitulated by Puffin prediction. (**A**) Schematic illustration of predicting TF depletion effect by in silico knock-out (KO) with Puffin. To predict the effects of TF depletion, we set the activation and consequently the effects of the corresponding motif to 0 in the Puffin model, and predict the transcription initiation pattern. (**B**) In silico KO prediction for NFY compared with experimental measurement of NFYA knockdown with Start-seq (subtracting control). Promoters with strong NFY contributions were selected and sorted by the predicted shifted TSS positions. (**C**) In silico KO prediction for YY1 compared with the experimental measurement with mNET-seq after induced depletion of YY1 (subtracting control). Promoters with strong YY1 contributions were selected and sorted by the position with the strongest predicted decrease in transcription initiation. The matrices shown in (**B**) and (**C**) heatmaps were smoothed with a small rectangular filter of size 5×1 for prediction and 5×3 for experimental data. (**D**) TSS with high NFY (top panel) or YY1 (bottom panel) contributions are significantly more influenced by corresponding TF depletion than those with low contributions (p-values derived from two-sided Wilcoxon rank sum test). (**E**) Example prediction and experimental measurements for TATA- and CCAAT-box (NFY) motif insertion and deletion experiments. All prediction and experimental values were shown in count scale and scaled to maximum 1. Positions of motifs inserted or deleted were indicated by colored bars.

Puffin estimated that the depletion of NF-Y transcription factor not only reduces transcription initiation at promoters with strong NFY contribution but also leads to an upstream shift of transcription initiation location in some TSS (**Fig. 3B**). Indeed the knockdown of NFYA(*17*) (encoding a subunit of NF-Y) led to an upstream shift of the TSS at locations exactly as predicted by Puffin (**Fig. 3B**, the experiment is performed in mouse and prediction is based on mouse genome sequence). Moreover, only NFY-contributing promoters based on Puffin showed strong changes in transcription while promoters with no strong NFY contribution are largely unaffected (**Fig. 3D**).

Puffin predicted that depleting YY1 has a unique effect of reducing transcription initiation at the immediate upstream of the motif, and a weaker effect about 200bp upstream of the first peak. We tested this effect using mNET-seq experimental data of induced YY1 depletion by auxin-inducible degron (AID)(*18*). Analysis of this data identified a previously unreported reduction of transcription initiation at the expected positions predicted by the model, including even the second weaker upstream peak of the YY1 motif effect (**Fig. 3C**). The transcription decrease was also selectively observed for YY1-contributing promoters by Puffin (**Fig. 3D**).

Next, we tested the model’s prediction of sequence edit effects on basepair resolution transcription initiation signals (**Fig. 3E**). Specifically, effects for transcription initiation signal changes with single basepair resolution were measured for TATA and NFY motifs deletion from promoters carrying these motifs, and insertions into neutral sequences with no promoter activity(*19*). The Puffin prediction not only recapitulated effects on expression level caused by sequence alterations (TSS-level correlation 0.88, Fig. S13A), but also displayed close to 0.9 basepair-level correlation with experimental data for TSS with strong expression which allowed basepair-level comparison (Fig. S13B). For example, deletion of both TATA or NFY motifs strongly decreased transcription in an LTR12 promoter (chr6:80516291-80517004, **Fig. 3E**), insertion of NFY motifs into a neutral sequence (chr5:164906550-164906800) alone was sufficient to drive transcription, and inserting both NFY and TATA motifs created a more focused promoter, consistent with experimental data (**Fig. 3E**). Puffin prediction also captured the shape of the transcription initiation signal, as well as changes of the shape upon motif insertions and deletions (**Fig. 3E**).

Taken together, experimental perturbations corroborated the Puffin-estimated position-specific motif effects on transcription initiation.

### Human promoters display diverse motif compositions that affect expression patterns

Puffin allows quantifying motif contributions to each promoter based on effects from each motif type, and we first characterized the basic properties of promoter motif compositions (**Fig. 4A-B**, Fig. S14, Table S2). We observed that promoters display very diverse motif combinations, and no motif is necessary for all promoters (**Fig. 4A-B**). Interestingly, we did not identify any strongly preferred or underrepresented motif combination relative to expectation based on single motif contribution (Fig. S14B), which suggests that motifs can be rather flexibly combined in human promoters. We estimated that the effective number of contributing motif types to each promoter is most commonly 2-3 (Fig. S14C).

**Fig. 4.**
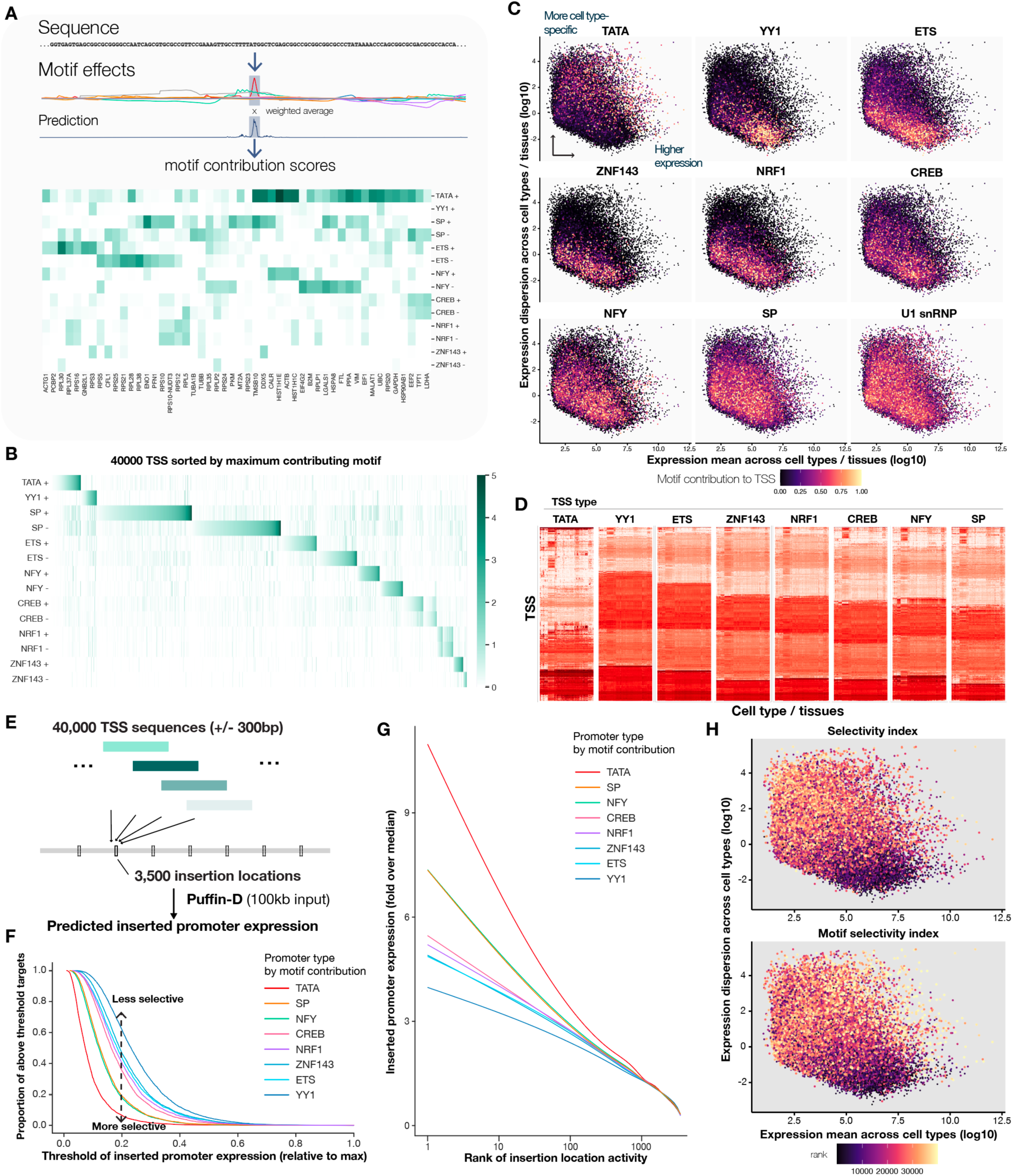
Motif compositions of promoters are linked to gene expression selectivity. (**A**) Schematic illustration of motif contribution score. The motif contribution score is computed by the weighted average of motif effects within +/-20bp to the annotated TSS, the weighting function is the prediction. Example contribution scores of 100 TSS with the highest expression are shown. (**B**) Motif contribution scores across 40,000 TSS with the highest expression based on FANTOM CAGE, sorted first by the maximum contribution motif type and then by the contribution score of that motif. (**C-D**) Motif contribution has a strong impact on expression dispersion across cell types and tissues. Scatterplots (**C**) show log expression dispersion (y-axis) and log mean (x-axis) with each dot indicating a TSS. The dots are colored by the contribution score of a motif type in each subpanel. Expression matrices (cell type / tissue x TSS) for representative TSS per motif type (TSS classified by maximum contributing motif type and filtered to TSS with top 2000 contribution scores from the classified motif type). (**E**) schematic overview of the promoter TSS insertion virtual screen with Puffin-D. 40,000 TSS sequences with 600bp length (+/− 300bp to the TSS) were inserted into 3500 target locations uniformly spaced over a 7Mb region (chr8:22964801-29964801). (**F-G**) Motif contribution affects selectivity to insertion targets in the virtual screen. (**F**) For each motif type (color), the top 1000 TSS by motif contribution scores were selected and the average predicted expression scores at each target location were computed, the proportions of target locations (y-axis) with predicted expression higher than the thresholds shown in the x-axis (scaled to proportion relative to the average of top-3 predicted expression levels across targets) were shown. (**G**) Generalized additive model-fitted curves of inserted promoter expression (scaled to fold over median expression across targets, y-axis) versus log rank of target activity (average target expression across 40,000 TSS) were shown. (**H**) The promoter selectivity index in the virtual screen (top panel) and motif selectivity index (bottom panel) are linked to gene expression dispersion across cell types.

Next, we asked how promoter composition is linked to gene expression properties. Analyzing TSS-level expression across >200 cell types and tissues from the FANTOM project, we found that motifs have remarkable effects on expression variation across cell types (**Fig. 4C-D**). Promoters with higher TATA contributions were most likely to be cell type-specific or have high dispersion index (variance / mean) across cell types and tissues. This is consistent with the previously known association between TATA and tissue-specific genes(*20*). Moreover, the prevalence of TATA motif within cell type-specific promoters was not due to a preference for gene expression in any specific tissue (Fig. S15). Interestingly, other motifs possess varying degrees of preference for ubiquitous expression patterns. Most strikingly, promoters with high YY1 contributions are most likely to be ubiquitously expressed or have low dispersion indices (**Fig. 4C-D**). High contributions from ETS, ZNF143, NRF1, and CREB motifs also favor more ubiquitous expression patterns. NFY and SP motifs are nearly neutral with only a slight preference for more ubiquitous expression patterns. U1 snRNP, which we estimated to mostly affect transcript abundance after transcription initiation exhibited neutral tissue-specificity. All promoter motifs are expected to bind at least one ubiquitously expressed TFs. Because the link between promoter motif and cell type-specificity cannot be explained by individual motif’s preference for any specific tissue, we hypothesize that promoter motifs influence how promoters respond to cell type-specific transcriptional regulatory signals including enhancers.

To explore the potential mechanism of this link, we studied the promoter response to context sequences and discovered a similar link between motif contribution and promoter selectivity for context sequences. We estimated promoter selectivity by inserting promoters into different genomic contexts and predicting the expression using the deep learning sequence model Puffin-D which takes 100kb sequence as input. In addition to accurately predicting the shape of transcription initiation signal, Puffin-D also excels at predicting the expression level of TSS, with >0.9 TSS-level and gene-level correlation with experimental data (Fig. S1-2). Thus, Puffin-D’s capability of utilizing long sequence context makes it well suited for studying the response of promoter to context sequences.

Specifically, for estimating promoter selectivity, each of 40,000 human promoter sequences (600bp) was inserted into 3,500 target locations that represent a diverse set of genomic contexts, and the expression levels of all inserted promoters were predicted (**Fig. 4G**). More selective promoters are defined as being highly expressed in a smaller subset of target locations and vice versa. Analysis of the 40,000 x 3,500 insertions revealed that motif contribution has a strong link to the selectivity of promoters (**Fig. 4H-J**, Fig. S16). Remarkably, motifs that demonstrated high and low selectivity are the same motifs that are linked to high and low expression dispersion across cell types/tissues (**Fig. 4E-F**).

Moreover, promoter selectivity from the virtual insertion screen is predictive of expression dispersion (**Fig. 4K**). By training a linear model to predict promoter selectivity from motif contribution, we obtained a sequence-based score of promoter selectivity, which is also predictive for high dispersion, tissue-specific expression patterns (Table S3).

Taken together, based on our results we propose that motif contributions determine the response curve of a promoter to external transcriptional activation signals, for example, TATA promoters are much more responsive than YY1 promoters to strong transcription activation signals. This mechanism can also explain the similarities between promoter selectivities to context sequences and cell-type-specific expression patterns. Thus promoter sequence composition likely plays a key role in determining gene expression patterns in conjunction with distal regulatory sequences.

### Sequence contributors underlying bidirectional transcription at promoters

Bidirectional transcription initiation is observed at most human promoters when measured at nascent transcript level(*13*, *14*, *21*, *22*). Whether such bidirectional transcription shares a common sequence basis has remained an open question(*23–25*).

The Puffin model provides an explanation of the sequence basis of bidirectional transcription initiation (**Fig. 2D** and **5A**). Bidirectional motifs with symmetric effects in both directions are the main contributors for most promoters (Table S2), thus they lead to substantial transcription initiation on the reverse strand, specifically at the level of nascent transcript. At the same time, directional motifs including TATA, YY1, and Long Inr contribute to preferential transcription initiation on a single strand, but even directional motifs can have activating effects on the other strand, such as YY1, or be partly palindromic such as TATA. On the other hand, most promoters are strongly directional at the mature transcript level even if they are bidirectional at nascent transcript level. As mentioned earlier, U1 snRNP motif exhibited a unique mechanism by contributing to the production of mature transcripts unidirectionally via post-transcription initiation effects (**Fig. 5A**), in consistency with prior results(*12–15*).

**Fig. 5.**
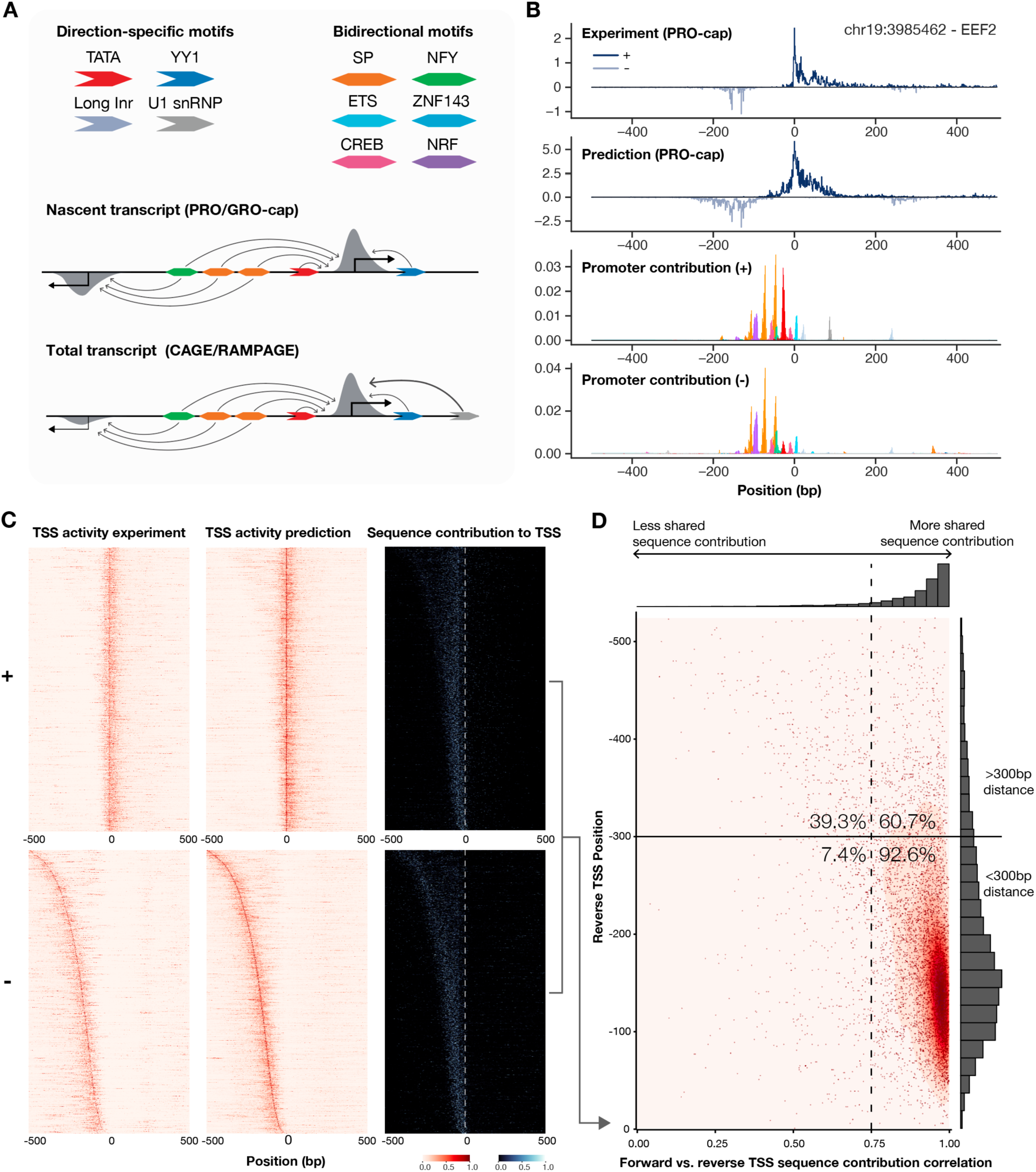
Sequence-basis of bidirectional transcription initiation at human promoters. (**A**) Schematic illustration of a sequence-based model of bidirectional transcription initiation. Directional motifs preferentially contribute to transcription initiation on the forward strand, and bidirectional motifs contribute to transcription initiation on both strands. Promoters with a high proportion of contribution from bidirectional motifs are bidirectional at the nascent transcript level. Most promoters are strongly directional at the mature transcript level, and the U1 snRNP motif contributes to a directional mature transcript outcome. (**B**). Basepair contribution analysis to an example TSS with bidirectional transcription initiation. The forward (+) and reverse strand (-) prediction and experimental measurements in log scale were shown in the top two panels. Reverse strand values were taken a negative sign. The bottom two panels showed per-motif basepair contribution scores to forward- and reverse-strand transcription respectively. All rows were scaled to maximum 1. (**C**) Basepair contribution scores for forward (top panel) and reverse (bottom panel) strand transcription of 8,216 promoters, sorted by reverse TSS position, scaled by the sum of positive basepair contribution scores per TSS and strand. The three columns are experimentally measured TSS activity, predicted TSS activity (PRO-cap), and basepair contribution score (PRO-cap) respectively. (**D**) High correlations between forward and reverse basepair contribution scores (x-axis) for reverse TSS with distance to forward TSS (y-axis) within about 300bp. The proportions of forward-reverse TSS pairs with less and greater than 0.75 correlation were indicated, for both pairs with less and greater than 300bp distance.

Beyond qualitative explanation, our model also allows quantifying the degree of shared sequence contribution for any forward-reverse TSS pair (**Fig. 5B-D**). To quantify how much sequence contribution is shared between forward and reverse strand TSS pairs, we selected 8,216 TSS with high expression levels in both directions based on PRO-cap (Methods). Puffin prediction well recapitulated forward and reverse directional transcription initiation site positions (**Fig. 5C**), which allowed us to further analyze basepair level sequence contribution to transcription initiation on both strands. Comparing basepair contribution scores for both strands (**Fig. 5D**, Fig. S17), the correlations were high for most TSS (87.7% with r > 0.75). More specifically, most of the reverse TSS that were within 300bp of the forward TSS have high correlations (92.6% with r > 0.75), while less shared sequence contribution was more common for reverse TSS that are further than 300bp away (60.7% with r > 0.75). Thus, the majority of bidirectional transcription initiation site pairs in close proximity (<300bp) share a major proportion of contributing sequence.

### Evolutionary conservation of promoter sequence determinant across mammal species

Finally, we asked whether the sequence dependencies captured by Puffin were conserved across mammalian species. Since high-coverage transcription initiation datasets are also available for mouse, we first assessed the cross-species generalization of transcription initiation models between human and mouse using the CAGE data from FANTOM project(*4*) (**Fig. 6A**). The models trained on mouse data found identical core motifs as the human model (Supplementary Data 3). Predictions of the human model and mouse model on the same sequences are also highly similar (median correlation of 0.967, **Fig. 6B**). Moreover, applying the human Puffin model to mouse sequences achieves almost identical performance as the mouse model (**Fig. 6B**). Thus the transcription initiation sequence dependencies learned by Puffin between human and mouse are nearly interchangeable.

**Fig. 6.**
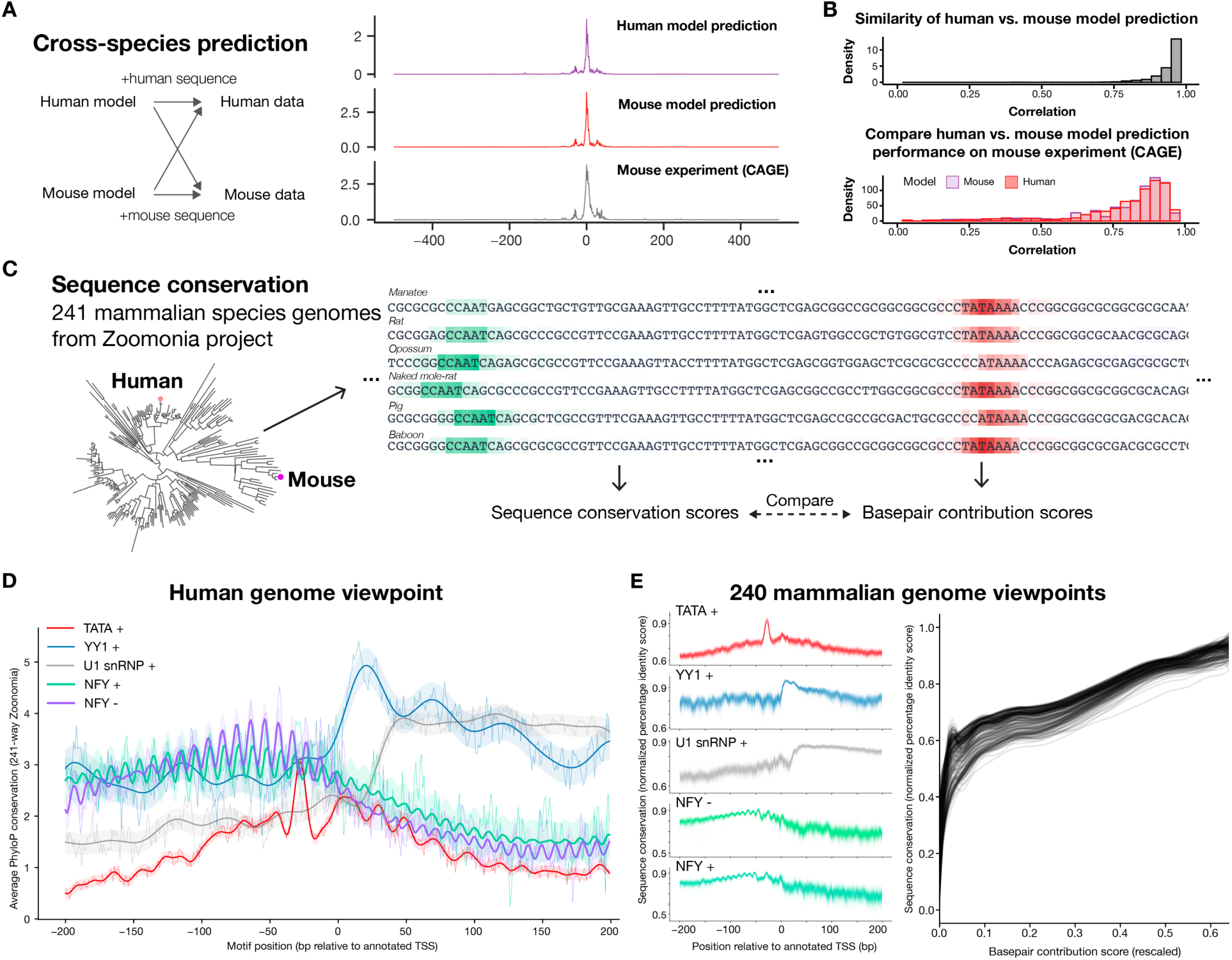
Cross-species generalization and conservation of promoter sequence rules across mammals. (**A**) Schematic illustration of cross-species prediction comparison of human and mouse Puffin transcription initiation models. Models trained on human and model data respectively are evaluated on predicting holdout promoters. (**B**) Human and mouse models have highly similar predictions (top panel) and almost equal performances (bottom panel) on the mouse genome. The distribution of correlations between human and model predictions on a mouse is shown in the top panel and the basepair level correlation for the human model (red) and mouse model (purple) are shown in the bottom panel. All comparisons are made on mouse holdout sequences (Methods). (**C**) Schematic illustration of sequence rule conservation analysis across 241 mammalian species. For each species, motif-specific basepair contribution scores are computed and compared with sequence conservation scores. (**D**) Position-specific sequence conservation scores for motifs share a similar pattern as our estimated position-specific motif effects, from a human genome viewpoint. X-axis shows the position (bp) relative to annotated TSS (human genome viewpoint) and the y-axis shows average PhyloP scores for each motif, computed with the average weighting by motif activation scores. This TSS-centered pattern is mirrored versus the motif-centered view. The bold line and shades indicate posterior mean and 95% credible intervals of Gaussian process regression (RBF kernel + White kernel + Periodic kernel for NFY, RBF kernel + White kernel for other curves). (**E**) Position dependencies of sequence conservation scores are also observed from the viewpoint of each of the 240 non-human mammalian species (left panel), with normalized percentage identity score (y-axis, Methods) with other species as the conservation metric. Basepair contribution scores computed by Puffin (right panel, x-axis, scores were transformed to the power of 0.25) are strong predictors of sequence conservations measured by percentage identity scores (y-axis) in all 241 mammalian species.

We next analyzed the conservation across mammalian species using 240 species genomes from the Zoonomia project(*7*) (**Fig. 6C**). We tested the hypothesis that position-specific evolutionary conservation patterns of the motifs are consistent with the TSS-centered position-specific motif effects inferred by the model. Specifically, we computed the average evolutionary conservation PhyloP scores for each motif across all positions relative to TSS (**Fig. 6D**). The spatial patterns of evolutionary conservation indeed showed striking similarities with position-specific motif effects. For example, TATA motifs are most conserved at ∼30bp upstream of TSS, YY1 motifs were most conserved immediately downstream of TSS, U1 snRNP motifs were most conserved starting from 50bp downstream of TSS, even the NFY motif’s ∼10.5bp periodic effect patterns were reflected in evolutionary conservation scores (conservation patterns for all motifs shown in Fig. S18). These results are all consistent with the motif’s estimated position-specific effects. Thus, evolutionary conservation provides another independent line of evidence for the position-specific motif effect patterns.

We showed that similar conservation patterns can be observed from the viewpoint of any of the 241 mammalian species (**Fig. 6E**). To assess the conservation of transcription initiation sequence rules across species, we analyzed each of the other 240 mammalian genomes, using the genome sequence for each species as input and sequence identities with all other species per base as the conservation measure. Moreover, we found that the basepair-level contribution score by Puffin is a strong predictor of evolutionary conservation in all 241 species (**Fig. 6E**). Thus, we expect that sequence dependencies of transcription initiation captured by Puffin will be widely applicable across mammalian species.

## Discussion

We have created the first model that explains the transcription initiation activity of most human promoter sequences at basepair level. This was achieved through the development of a deep learning-inspired explainable modeling approach, which allowed for learning a compact model that provides insights into the sequence basis transcription initiation. Puffin both recapitulated known biology and provided new predictions that are well supported by experimental results and evolutionary conservation. Importantly, we discovered that sequence determinants of transcription initiation can be effectively described using a small set of rules, ultimately making systematic and quantitative analysis of this process tractable.

Thus, our model and analyses shed light on many questions related to promoter sequence and function that we set out to answer: for most human promoters, we can now identify individual basepairs and motifs that contribute to transcription initiation; a simple additive / multiplicative model of sequence pattern effects is sufficient to recapitulate most of the transcription initiation activity; we discovered new connections between promoter sequence compositions, cell type-specificity of genes, and promoter selectivity; we provided an explanation for the sequence-basis of bidirectional transcription and strand preference in human promoters; finally, with Puffin we can now show that the sequence rules of promoters are conserved across mammalian species.

Moreover, we expect that the sequence-level understanding of transcription initiation and the molecular mechanism-level understanding will eventually converge, and future research will likely discover the molecular and structural underpinnings of each motif’s position-specific effects.

Overall, our study provides systematic insights into the sequence determinants of transcription initiation in the human genome and beyond, as well as a powerful tool for understanding and engineering promoter sequences and gene regulation across species. Our findings also underscore the scientific importance of explainable machine learning and computational modeling for unraveling the sequence-based mechanisms governing genome regulation, which we expect to further unravel the sequence rules underlying diverse genomic functions.

## Methods

### Processing of transcription initiation signal datasets

All transcription initiation datasets (FANTOM CAGE, ENCODE CAGE, ENCODE RAMPAGE, GRO-cap, PRO-cap) were downloaded from published datasets listed in Supplementary Data 1. As individual signal profiles do not have enough coverage for providing accurate estimates at basepair resolution except for the most highly expressed TSS, we aggregated all signal profiles measured by the same technique. Specifically, the basepair-resolution count profiles were averaged after applying log_10_(*x* + 1) transformation where *x* is the read count, with plus and minus strand profiles aggregated separately. The addition of the pseudocount can be interpreted as applying a uniform Dirichlet prior and obtaining the posterior mean. In this manuscript, we refer to this aggregated signal as the log scale and its inverse transformed value by 10^#^ − 1 as the count scale.

We next addressed the known bias for poly-T sequence in the FANTOM CAGE aggregated signal profile, which is specific to the HelioScope CAGE protocol used for FANTOM CAGE datasets(*26*), but does not affect ENCODE CAGE datasets. After analyzing FANTOM CAGE signal dependency on the number of consecutive “T”s across the genome (Fig. S19), we applied a filter with a threshold of >=8 consecutive “T”s, and masked the signals in [-6, +10) interval relative to the end of the poly-T sequence. The filtered FANTOM CAGE signal profile is replaced with ENCODE CAGE signal rescaled to the average signal level of FANTOM CAGE. This correction only affects a minor fraction of the genome (0.46%).

We obtained human TSS annotation from the FANTOM-CAT catalog at the “robust” level(*27*) and removed TSS with inconsistent expression levels across datasets. We then rank all TSS by expression level in the aggregated FANTOM CAGE profile, quantified by the sum over +/− 20bp at TSS in count scale (we refer to this ranking when we used top-N TSS in this manuscript). Specifically, for removing TSS with inconsistent expression levels, non-protein-coding TSS that are >40-fold lower in expression value than FANTOM CAGE in both datasets, after scaling by total expression values across all TSS for each dataset, were removed. In addition, for performance evaluation, we selected high-confidence TSS that have consistent basepair-resolution transcription initiation profiles across experimental techniques. Specifically, for the 1000bp interval centered at the TSS position, only TSS with FANTOM CAGE profile with high correlation with at least one other dataset (>0.5 for ENCODE CAGE, >0.4 for ENCODE RAMPAGE, GRO-cap, or PRO-cap datasets) were preserved. The filtered list of TSS with high-confidence TSS labels is provided in Table S4.

### Interpretation-focused Puffin model for transcription initiation

The Puffin model consists of two learnable layers, a sequence pattern activation layer, and a transcription initiation effect layer (Fig. S4). Both layers were trained from scratch based on only sequence and transcription initiation signal data, without using any known motifs. Both layers’ computations can be represented by convolution layers. The convolution kernels of the first layer represent sequence patterns’ position-specific weights and the convolution kernels of the second layer represent sequence pattern activations’ position-specific effects on transcription initiation.

Since Puffin uses three sequence pattern types with different sizes (kernel size in the first layer) and effect ranges (kernel size in the second layer), each layer of the model contains several parallel convolution layers each corresponding to a sequence pattern type. Specifically, the sequence pattern sizes of motifs, initiators, and trinucleotides are 51bp, 15bp, and 3bp respectively, and the effect ranges are +/−300bp, +/−7bp, and +/− 300bp respectively. The sequence pattern sizes and effect ranges specified in the model are the maximum that the model can learn and the effective sequence pattern size and effect ranges learned are usually lower. We note that the initial design of Puffin uses only the motif sequence patterns, and more parsimonious sequence patterns with smaller sizes or effect ranges including initiators and trinucleotides were introduced in a data-driven process to obtain a more compact and interpretable design while maintaining the performance (Supplementary Text 1).

The first layer uses the softplus activation function to compute the sequence pattern activations. Moreover, the activations of each reverse-complement sequence pattern were also computed, which doubles the number of channels from the first layer output. For the second layer, we used FFT-based convolution for computing motif and trinucleotide effects because the large kernel size makes FFT-based convolution more efficient than regular convolution implementation. Finally, the second layer computes sequence pattern effects for 10 targets, which corresponds to 5 different techniques on both forward and reverse strands, thus the second layer learns a separate set of sequence pattern effects for each target.

The sum of sequence pattern effects per position were transformed by the softplus function to the final prediction. The softplus function is deliberately chosen here to make the input and output interpretable. In this formulation, the pre-activation sum of sequence pattern effects *x* is comparable to ln(*s*) where *s* is the transcription initiation signal in count scale, because we train the model by fitting the scaled softplus function output log_10_(exp(*x*) + 1) to the log scale transcription initiation signal log_10_(*s* + 1). Thus, the additive combination of sequence pattern effects in *x* space can be considered as a multiplicative combination in *s* space. Similarly, the final Puffin model prediction is comparable to natural log scale transcription initiation signal ln(*s* + 1) and can be converted to count scale using exponential minus 1 transform.

The main training loss function is the Kullback-Leibler(KL) divergence loss

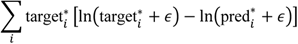

Where 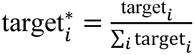, 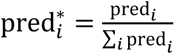, *i* is the position index, and *ε* is set to 1e-10. The target is processed as introduced in the previous section. The KL divergence loss is sensitive to the shape but not the scale of the transcription initiation signal. This is intended as the overall abundance is dependent on not only the promoter. The prediction and target are both divided by the sum over the 4kb region before computing KL divergence loss. Since the KL-divergence loss does not constrain the scale of the prediction, we added an auxiliary loss that matches the exact transcription signal values. This auxiliary loss also increased the interpretability of the prediction by allowing it to be considered as a prediction of transcription initiation signals, even though the training emphasizes learning the shape rather than the scale of the transcription initiation signal. Specifically, the auxiliary loss is

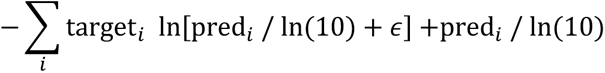

where *i* indicates the position and *ε* is 1e-10. We refer to this loss as the pseudo-Poisson loss as it has the same form as the Poisson loss function even though the target values are not counts. The auxiliary loss is weighted by a factor of 1e-3. The losses for all ten targets were averaged. Additional regularization terms were added to the loss function including L1 regularization to kernel weights in both layers, and L2 smoothness regularization between spatially adjacent kernel weights in the second layer, for motifs and trinucleotides.

The models were trained to predict from one-hot encoded sequence to transcription initiation signals on both strands for 5 experimental techniques. Specifically, the model was trained with the task of predicting transcription initiation signal in the 4kb region surrounding each TSS with a random strand selected for each training sample. Top 40,000 high-confidence TSS ranked by expression level as described in the previous section were used. We divide the genome into the training set: all chromosomes except for chr8, 9, and 10, the validation set: chr10, and the test set: chr8 and 9. The training data was retrieved on-the-fly during training and the strand is selected randomly for each training sample.

We trained the Puffin model in three stages to make sure that all motifs included were reproducibly discovered across multiple training replicates. In the first stage, we trained 12 replicates with different random seeds. The first stage model differs from the final Puffin architecture in that it learns 40 motif sequence patterns and 10 initiator sequence patterns, and the SiLU activation function was used for the first layer because it facilitates motif discovery. A nonredundant set of 10 motifs that were reproducible (> 0.95 maximum cross-correlation across >7 replicates) were chosen as the consensus motifs. In the second stage, we use the final Puffin model architecture as described, and sequence patterns and effects were initialized by the consensus sequence patterns from the first stage. Since not all sequence patterns learned were located at the center of the 51bp motif kernel, in the third stage, we centered the motif sequence patterns and continued training.

The model can process variable-length input, and the raw model output size equals the input sequence length. But since predictions near the edge can be affected by padding, to remove the effect of padding, valid predictions were trimmed from each end by 325bp. Therefore, for example, to obtain N-bp prediction we use N + 650bp long sequence as an input.

For evaluation of the prediction performance, we computed Pearson correlation between experiment and prediction per TSS for 1kb windows centered at each TSS. High confidence TSS on test chromosomes among the top 100,000 TSS were used for evaluation. For downstream analysis, FANTOM CAGE prediction and contribution scores were used unless otherwise indicated.

### Visualization of Puffin motifs and position-specific effect curves

For visualization of sequence patterns such as motifs, the position-specific weight matrices were directly obtained from the kernel weights of the first layer. We note that adding or subtracting a value to all four bases at the same position does not change the layer output (up to a constant which can be canceled out by bias term), thus for standardization we processed the motif position-specific weight matrices per position by first subtracting the mean and then subtracting 0.7x average absolute value per position (the later subtraction highlights the most positive base in visualization, otherwise typically more than one base were highlighted due to subtraction by mean).

We visualize the position-specific effect curve of each motif in motif-centered coordinates, which were obtained by reversing the spatial dimension of the kernel weights of the second layer.

### Prediction-focused Puffin-D model for transcription initiation

Puffin-D is the prediction-focused architecture that captures sequence dependencies up to 100kb. Different from Puffin which is trained with KL divergence loss to predict the shape of the transcription initiation signal, Puffin-D is trained to predict the exact value of the transcription initiation signal. The training loss function is the pseudo-Poisson loss, which is the same as the auxiliary loss function for training Puffin except for that it was computed for 100kb intervals.

To utilize long-range sequence information efficiently, the Puffin-D architecture uses an architecture that iteratively propagates information across sequence locations via two upward-downward passes. The upward passes integrate long-range sequence information hierarchically and the downward passes distribute integrated information to local sequence representations (Fig. S20). Residual connections between the same levels of the upward and downward passes allow information at all spatial resolutions to be preserved. Individual convolution blocks were modified over our previous design(*28*) by adding size 1 convolution layers, replacing ReLU with SiLU activation function, and replacing max pooling with strided convolution.

To evaluate Puffin-D performance, we generated predictions for entire test chromosomes chr8 and chr9, with a sliding window step size of 50kb, and the center 50kb of each 100kb prediction was used. Regions within 1kb to unknown bases or 25kb to chromosome ends were excluded. In addition to basepair level correlation, we computed correlations at the transcript level and gene level. At the transcript level, we aggregated prediction and experimental signal at count scale within 400bp window to each annotated transcription start site; at gene level, we further summed all transcript-level prediction and experimental signal per gene.

### Sequence contribution scores

An important advantage of the interpretation-focused Puffin architecture allows quantitatively analyzing sequence contribution to transcription initiation at motif and basepair levels. Here we describe the definition and interpretation for each motif and basepair-level contribution score:

#### Motif contribution score

Motif contribution score represents the amount of contribution from a motif type to any position or window surrounding TSS. The motif contribution to any position can be directly represented by motif effects in the Puffin model, because sequence pattern effects are additively combined, and the baseline for motif effects is zero due to regularization that drives the motif effect to zero at long distance. Thus, motif contribution is almost synonymous with motif effects in Puffin model, but motif contribution scores can also be computed by aggregating over a window in a weighted or unweighted fashion.

In this manuscript, the motif contribution scores for each TSS were computed by the weighted average motif effects within the 20bp window centered at the TSS, and the weights are the predicted transcription initiation signals. We provide the full contribution scores in Table S2. For downstream analysis, we summed the motif contribution scores for forward and reverse directional motifs for bidirectional motifs, and for directional motifs, only the + direction motif contribution scores were retained. U1 snRNP (post-transcription initiation effect) and Long Inr (initiator sequence pattern) were usually excluded from motif contribution analysis, because U1 snRNP has mostly post-transcriptional initiation effects, and Long Inr effects are initiator-like and do not have zero baselines like other motifs.

#### Basepair contribution score to transcription initiation

We also refer to this score as the basepair contribution score, which represents the amount of contribution to transcription initiation from each basepair. Moreover, basepair contribution score can also be decomposed to per motif type scores, allowing dissecting basepair level sequence contribution by motifs. The basepair contribution score equals the sum of basepair contribution scores for all motifs.

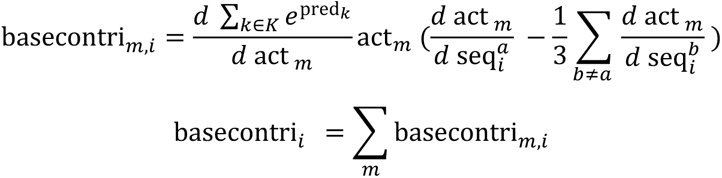

Where basecontri*_m,i_* indicates the basepair contribution score for motif *m* and the base at position *i*, pred*_k_* indicates the model prediction at the position *k*, and *K* indicates the window of transcription initiation prediction that basepair contribution scores are compute for. For notational simplicity we use act *_m_* to denote the vector of motif activation across all positions for the motif 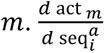 indicates the gradient with respect to the true base *a* at sequence position *i*, while 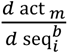 indicates the gradient with respect to any other base *b* at sequence position *i*. The basepair contribution score can be computed for any of the 10 targets that Puffin predicts by choosing the corresponding predictions.

The basepair contribution score can be computed with respect to transcription initiation signals in any position or window. In this manuscript we computed it for 1kb windows surrounding each TSS, using 1650bp sequences. Moreover, in all analyses, we further scale the basepair contribution score as defined above by dividing the sum of positive basepair contribution scores within the 1kb window per TSS. This score was used to compare sequence contributions underlying bidirectional TSS pairs on both strands and to analyze the relationship between basepair contribution score and sequence conservation across species.

#### Basepair contribution score to motif activation

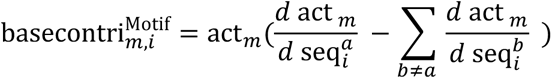

The basepair contribution score to motif activation represents the amount of contribution to motif activation from each basepair, which is computed using only the first layer of Puffin. This score was used in the analysis of position-specific evolutionary conservation patterns of each motif and notably it does not involve the second layer or any learned motif effects to compute.

### Puffin in silico knock-out for TF perturbation effect prediction

As the Puffin model dissects sequence dependencies into effects from individual sequence patterns, which are mapped to TFs, we can simulate the effects of TF depletion by removing the effects from that motif using the Puffin model. We refer to this technique as “in silico KO”, which mimics the effect of acutely removing the TFs that bind to any motif. To perform in silico KO, we set the motif activation scores for the corresponding motif to 0 and continue the subsequent computations, which also set the effects of that motif effect to 0.

We evaluated in silico KO predictions by comparisons with experimental data from NFY and YY1-depletion datasets(*17*, *18*). We first selected for TSS with a strong contribution from NFY and YY1, quantified by the sum of absolute predicted difference over the 1kb window centered at each TSS. A threshold of 20 and 55 was used for YY1 and NFY respectively. We demonstrated that the expression level of TSS above this threshold was significantly decreased compared to TSS without YY1 or NFY contribution with a threshold of 1, with a two-sided Wilcoxon rank sum test. The expression levels of TSS were quantified by −50 to +50bp for NFY Start-seq and −50 to +100bp for YY1 mNET-seq data (mNET-seq signals tend to be shifted downstream from the TSS). We then compared the predicted basepair resolution effect of TF depletion with experimental data for the selected TSS with heatmap visualization.

### Puffin prediction of motif insertion and deletion effects

We compared Puffin prediction with the motif perturbation dataset published by(*19*), which measured the effects of TATA and NFY motif mutations in human promoters and insertions of these motifs into neutral sequences. The transcriptional activities of wildtype and mutants were measured using STAP-seq in 500 oligomers with 5 replicates for every sample. The oligomer sequence was obtained from the data and the surrounding sequence was retrieved from the human STAP-seq screening vector sequence (Addgene ID: 125150). Both basepair level and TSS-level prediction and experimental measurements were compared. TSS-level quantification was computed by summing count scale predictions or signals over the 250bp oligonucleotide.

### Promoter motif contribution and expression selectivity

Motif contribution scores for TSS were computed as described in the “Sequence contribution scores” section. TSS-level expression values across >200 cell types and tissues were obtained from FANTOM CAGE profiles quantified by the sum over +/− 20bp at TSS in count scale. CAGE profiles for samples for the same cell type or tissue were summed at count scales. The raw TSS-level counts for each cell type / tissue were then normalized by dividing the size factors estimated by the DESeq2 package(*29*). The mean and dispersion of TSS-level expression across cell types and tissues were then computed from normalized counts and compared with motif contribution scores.

For comparison of TSS expression patterns across TSS types by motif contribution, we selected the top 5% TSS by motif contribution score for each motif type among the top 40,000 TSS. The expression matrix of all 40,000 TSS were hierarchically clustered, and the row and column orders were preserved in the visualization of individual TSS types.

To estimate promoter selectivity to genome contexts, we performed an in silico insertion screen with Puffin-D model. For insertion sequences, we used 600bp centered at each of the top 40,000 TSS, and for target locations, we selected 3500 locations uniformly spaced in the genomic interval chr8:22964801-29963540 with the step size 2000bp. The target locations are on a chromosome that was held out from the model training. For each of the 40,000 x 3,500 in silico insertion experiments, we replaced 600bp of the target genome sequence with the 600bp insertion sequence. The prediction for the transcription initiation signal was made using the Puddin-D model. The predicted transcription initiation signal, or expression level, was measured by the mean FANTOM CAGE prediction in count scale within +/−20bp from the center position.

The selectivity score for each TSS was defined as the proportion of insertion targets with predicted expression levels below the threshold. The threshold for selectivity score computation for each TSS was defined as 1/3 of the mean of the top-3 predicted expression levels across target locations. Motif selectivity score is estimated by training an L2 regularized linear regression model to predict log-odds of selectivity score from motif contribution scores, and the regularization parameter was selected by leave-one-out cross-validation in closed form.

### Basepair contribution scores for bidirectional TSS

TSS with high expression levels in both strands were selected based on the aggregated PRO-cap transcription initiation signal in count scale on both strands. Specifically, both the sum of the forward strand PRO-cap signal within −250 to +250bp window, and the sum of reverse strand signal within −500 to 0bp window are >50. Moreover, we selected TSS for which the maximum signal position on the reverse strand is upstream of the maximal signal position on the forward strand, the maximal signal position on the forward strand is within 50bp to the annotated TSS position, and the maximal signal position on the reverse strand is upstream of the annotated TSS position. Basepair contribution scores to transcription initiation for PRO-cap on both strands were computed and compared by correlation (within +/−500bp window to the annotated TSS).

### Comparison of promoter sequence dependencies in human and mouse

To compare human and mouse sequence dependencies captured by Puffin, we trained Puffin models on mouse FANTOM CAGE data(*4*), aggregated with the same procedure as human FANTOM CAGE data. The consensus motifs from mouse training replicates were analyzed in the same process as for human and showed nearly identical motifs. We next compared predictions between human and mouse models applied to the mouse genome. To ensure that the comparison is appropriately done, we liftover the mouse TSS annotation from the FANTOM project from mm10 to hg38. The top 40,000 mouse TSS ranked in the same process as described for the human TSS were used for this analysis. The mouse models were trained on TSS for which the liftover coordinates were in human training chromosomes. Similarly, the evaluations were performed on mouse TSS for which the liftover coordinates in hg38 locate in the human holdout chromosomes. We compared human and mouse Puffin models after stage 1 training, and the average prediction across 12 training replicates was used for both human and mouse. Human and mouse model predictions for 1kb regions centered at TSS that belong to the test set were also computed and compared with mouse FANTOM CAGE signals.

### Evolutionary conservation across 241 mammalian genomes

We obtained 241-way mammalian genome alignment from the Zoonomia project and downloaded the PhyloP scores from the UCSC genome browser. To analyze position-specific evolutionary conservation for each motif from the human genome viewpoint, for every basepair position relative to the annotated TSS from −200bp to +200bp, we computed the weighted average PhyloP score across top 4000 TSS for each motif. The weight used is the basepair level motif activation score for that motif.

To analyze evolutionary conservation from the viewpoint of each of the other 240 genomes, human genome TSS positions were liftover to each genome, and the 1650bp sequences centered at the TSS in each genome assembly were retrieved. Basepair contribution scores to motif activation and transcription initiation were computed separately for sequences from each genome. We used percentage identity with the rest of the 240 genomes as the raw sequence conservation score. Percentage identity scores per basepair were computed based on multi-sequence alignment generated by MAFFT(*30*). Since each species has a different distribution of evolutionary distance from other species, the raw percentage identity scores are not directly comparable in scale with each other. To make percentage identity scores comparable across species we applied quantile normalization by linearly scaling. The linear scaling matches each species’ 0.1 and 0.9 quantiles to the human quantiles. For each species, in addition to computing position-specific evolutionary conservation scores with normalized percentage identity scores, we also fitted species-specific curves representing the relationship between scaled basepair contribution scores (transformed to the power of 0.25) and normalized percentage identity scores by generalized additive model.

## Data availability

The human GRCh38/hg38 reference genome was used for training the Puffin and Puffin-D models. The mouse models were trained with the mm10 reference genome. All coordinates in the manuscript refer to GRCh38/hg38 unless otherwise indicated. All transcription initiation signal datasets used for model training (FANTOM CAGE, ENCODE CAGE, ENCODE RAMPAGE, GRO-cap, PRO-cap) are listed in Data S1. The experimental validation datasets are obtained from NCBI GEO accessions GSE178982 (YY1-AID), GSE115110 (NFYA knockdown), and GSE156741 (TATA / NFY motif perturbation). The 241 Zoonomia genomes were obtained from the data portal https://zoonomiaproject.org/the-data/. All data used in this manuscript were also deposited into Zenodo repository https://zenodo.org/record/7954971.

## Code availability

The code and model for Puffin and Puffin-D are available at https://github.com/jzhoulab/puffin. The code for reproducing the analyses in the manuscript is available at https://github.com/jzhoulab/puffin_manuscript. A user-friendly website for using Puffin is available at https://tss.zhoulab.io.

## Acknowledgments

The authors thank Ronald C. Conaway, Kathleen M. Chen, Christopher Y. Park, and Jian Xu for feedbacks to this manuscript. This works is conducted on the high-performance computing facility BioHPC at the University of Texas Southwestern Medical Center. This work is supported by National Institutes of Health grant DP2GM146336, Cancer Prevention and Research Institute of Texas grant RR190071, and UTSW Endowed Scholar award to JZ.

## References

1. S. T. Smale, J. T. Kadonaga, The RNA polymerase II core promoter. Annu. Rev. Biochem. 72, 449– 479 (2003).

2. A. Sandelin, P. Carninci, B. Lenhard, J. Ponjavic, Y. Hayashizaki, D. A. Hume, Mammalian RNA polymerase II core promoters: insights from genome-wide studies. Nat. Rev. Genet. 8, 424–436 (2007).

3. Y.-L. Wang, G. A. Kassavetis, J. T. Kadonaga, Others, The punctilious RNA polymerase II core promoter. Genes Dev. 31, 1289–1301 (2017).

4. A. R. R. Forrest, H. Kawaji, M. Rehli, J. K. Baillie, M. J. L. De Hoon, V. Haberle, T. Lassmann, I. V. Kulakovskiy, M. Lizio, M. Itoh, R. Andersson, C. J. Mungall, T. F. Meehan, S. Schmeier, N. Bertin, M. Jørgensen, E. Dimont, E. Arner, C. Schmidl, U. Schaefer, Y. A. Medvedeva, C. Plessy, M. Vitezic, J. Severin, C. A. Semple, Y. Ishizu, R. S. Young, M. Francescatto, I. A. Altschuler, D. Albanese, G. M. Altschule, T. Arakawa, J. A. C. Archer, P. Arner, M. Babina, S. Rennie, P. J. Balwierz, A. G. Beckhouse, S. Pradhan-Bhatt, J. A. Blake, A. Blumenthal, B. Bodega, A. Bonetti, J. Briggs, F. Brombacher, A. M. Burroughs, A. Califano, C. V. Cannistraci, D. Carbajo, Y. Chen, M. Chierici, Y. Ciani, H. C. Clevers, E. Dalla, C. A. Davis, M. Detmar, A. D. Diehl, T. Dohi, F. Drabløs, A. S. B. Edge, M. Edinger, K. Ekwall, M. Endoh, H. Enomoto, M. Fagiolini, L. Fairbairn, H. Fang, M. C. Farach-Carson, G. J. Faulkner, A. V. Favorov, M. E. Fisher, M. C. Frith, R. Fujita, S. Fukuda, C. Furlanello, M. Furuno, J. I. Furusawa, T. B. Geijtenbeek, A. P. Gibson, T. Gingeras, D. Goldowitz, J. Gough, S. Guhl, R. Guler, S. Gustincich, T. J. Ha, M. Hamaguchi, M. Hara, M. Harbers, J. Harshbarger, A. Hasegawa, Y. Hasegawa, T. Hashimoto, M. Herlyn, K. J. Hitchens, S. J. H. Sui, O. M. Hofmann, I. Hoof, F. Hori, L. Huminiecki, K. Iida, T. Ikawa, B. R. Jankovic, H. Jia, A. Joshi, G. Jurman, B. Kaczkowski, C. Kai, K. Kaida, A. Kaiho, K. Kajiyama, M. Kanamori-Katayama, A. S. Kasianov, T. Kasukawa, S. Katayama, S. Kato, S. Kawaguchi, H. Kawamoto, Y. I. Kawamura, T. Kawashima, J. S. Kempfle, T. J. Kenna, J. Kere, L. M. Khachigian, T. Kitamura, S. P. Klinken, A. J. Knox, M. Kojima, S. Kojima, N. Kondo, H. Koseki, S. Koyasu, S. Krampitz, A. Kubosaki, A. T. Kwon, J. F. J. Laros, W. Lee, A. Lennartsson, K. Li, B. Lilje, L. Lipovich, A. Mackay-sim, R. I. Manabe, J. C. Mar, B. Marchand, A. Mathelier, N. Mejhert, A. Meynert, Y. Mizuno, D. A. L. De Morais, H. Morikawa, M. Morimoto, K. Moro, E. Motakis, H. Motohashi, C. L. Mummery, M. Murata, S. Nagao-Sato, Y. Nakachi, F. Nakahara, T. Nakamura, Y. Nakamura, K. Nakazato, E. Van Nimwegen, N. Ninomiya, H. Nishiyori, S. Noma, T. Nozaki, S. Ogishima, N. Ohkura, H. Ohmiya, H. Ohno, M. Ohshima, M. Okada-Hatakeyama, Y. Okazaki, V. Orlando, D. A. Ovchinnikov, A. Pain, R. Passier, M. Patrikakis, H. Persson, S. Piazza, J. G. D. Prendergast, O. J. L. Rackham, J. A. Ramilowski, M. Rashid, T. Ravasi, P. Rizzu, M. Roncador, S. Roy, M. B. Rye, E. Saijyo, A. Sajantila, A. Saka, S. Sakaguchi, M. Sakai, H. Sato, H. Satoh, S. Savvi, A. Saxena, C. Schneider, E. A. Schultes, G. G. Schulze-Tanzil, A. Schwegmann, T. Sengstag, G. Sheng, H. Shimoji, Y. Shimoni, J. W. Shin, C. Simon, D. Sugiyama, T. Sugiyama, M. Suzuki, N. Suzuki, R. K. Swoboda, P. A. C. ’T Hoen, M. Tagami, N. T. Tagami, J. Takai, H. Tanaka, H. Tatsukawa, Z. Tatum, M. Thompson, H. Toyoda, T. Toyoda, E. Valen, M. Van De Wetering, L. M. Van Den Berg, R. Verardo, D. Vijayan, I. E. Vorontsov, W. W. Wasserman, S. Watanabe, C. A. Wells, L. N. Winteringham, E. Wolvetang, E. J. Wood, Y. Yamaguchi, M. Yamamoto, M. Yoneda, Y. Yonekura, S. Yoshida, S. E. Zabierowski, P. G. Zhang, X. Zhao, S. Zucchelli, K. M. Summers, H. Suzuki, C. O. Daub, J. Kawai, P. Heutink, W. Hide, T. C. Freeman, B. Lenhard, L. V. B. Bajic, M. S. Taylor, V. J. Makeev, A. Sandelin, D. A. Hume, P. Carninci, Y. Hayashizaki, A promoter-level mammalian expression atlas. Nature (2014), doi:10.1038/nature13182.

5. H. Xi, Y. Yu, Y. Fu, J. Foley, A. Halees, Z. Weng, Analysis of overrepresented motifs in human core promoters reveals dual regulatory roles of YY1. Genome Res. 17, 798–806 (2007).

6. C. G. de Boer, E. D. Vaishnav, R. Sadeh, E. L. Abeyta, N. Friedman, A. Regev, Deciphering eukaryotic gene-regulatory logic with 100 million random promoters. Nat. Biotechnol. 38, 56–65 (2020).

7. Zoonomia Consortium, A comparative genomics multitool for scientific discovery and conservation. Nature. 587, 240–245 (2020).

8. Y. Hayashizaki, Cap Analysis Gene Expression (CAGE). Cap-Analysis Gene Expression (Cage) (2019), pp. 1–6.

9. J. E. Moore, X.-O. Zhang, S. I. Elhajjajy, K. Fan, H. E. Pratt, F. Reese, A. Mortazavi, Z. Weng, Integration of high-resolution promoter profiling assays reveals novel, cell type–specific transcription start sites across 115 human cell and tissue types. Genome Res. 32, 389–402 (2022).

10. M. T. Weirauch, A. Yang, M. Albu, A. G. Cote, A. Montenegro-Montero, P. Drewe, H. S. Najafabadi, S. A. Lambert, I. Mann, K. Cook, H. Zheng, A. Goity, H. van Bakel, J. C. Lozano, M. Galli, M. G. Lewsey, E. Huang, T. Mukherjee, X. Chen, J. S. Reece-Hoyes, S. Govindarajan, G. Shaulsky, A. J. M. Walhout, F. Y. Bouget, G. Ratsch, L. F. Larrondo, J. R. Ecker, T. R. Hughes, Determination and inference of eukaryotic transcription factor sequence specificity. Cell (2014), doi:10.1016/j.cell.2014.08.009.

11. E. Seto, Y. Shi, T. Shenk, YY1 is an initiator sequence-binding protein that directs and activates transcription in vitro. Nature. 354, 241–245 (1991).

12. H. Vlaming, C. A. Mimoso, A. R. Field, B. J. E. Martin, K. Adelman, Screening thousands of transcribed coding and non-coding regions reveals sequence determinants of RNA polymerase II elongation potential. Nat. Struct. Mol. Biol. 29, 613–620 (2022).

13. L. J. Core, A. L. Martins, C. G. Danko, C. T. Waters, A. Siepel, J. T. Lis, Analysis of nascent RNA identifies a unified architecture of initiation regions at mammalian promoters and enhancers. Nat. Genet. 46, 1311–1320 (2014).

14. A. E. Almada, X. Wu, A. J. Kriz, C. B. Burge, P. A. Sharp, Promoter directionality is controlled by U1 snRNP and polyadenylation signals. Nature. 499, 360–363 (2013).

15. C. A. Mimoso, K. L. Adelman, U1 snRNP increases RNA pol II elongation rate to enable synthesis of long genes. SSRN Electron. J. (2022), doi:10.2139/ssrn.4296553.

16. S. T. Smale, D. Baltimore, The “initiator” as a transcription control element. Cell. 57, 103–113 (1989).

17. A. J. Oldfield, T. Henriques, D. Kumar, A. B. Burkholder, S. Cinghu, D. Paulet, B. D. Bennett, P. Yang, B. S. Scruggs, C. A. Lavender, E. Rivals, K. Adelman, R. Jothi, NF-Y controls fidelity of transcription initiation at gene promoters through maintenance of the nucleosome-depleted region. Nat. Commun. 10, 3072 (2019).

18. T.-H. S. Hsieh, C. Cattoglio, E. Slobodyanyuk, A. S. Hansen, X. Darzacq, R. Tjian, Enhancer– promoter interactions and transcription are largely maintained upon acute loss of CTCF, cohesin, WAPL or YY1. Nature Genetics. 54 (2022), pp. 1919–1932.

19. C. Neumayr, V. Haberle, L. Serebreni, K. Karner, O. Hendy, A. Boija, J. E. Henninger, C. H. Li, K. Stejskal, G. Lin, K. Bergauer, M. Pagani, M. Rath, K. Mechtler, C. D. Arnold, A. Stark, Differential cofactor dependencies define distinct types of human enhancers. Nature. 606, 406–413 (2022).

20. P. Carninci, A. Sandelin, B. Lenhard, S. Katayama, K. Shimokawa, J. Ponjavic, C. A. M. Semple, M. S. Taylor, P. G. Engström, M. C. Frith, A. R. R. Forrest, W. B. Alkema, S. L. Tan, C. Plessy, R. Kodzius, T. Ravasi, T. Kasukawa, S. Fukuda, M. Kanamori-Katayama, Y. Kitazume, H. Kawaji, C. Kai, M. Nakamura, H. Konno, K. Nakano, S. Mottagui-Tabar, P. Arner, A. Chesi, S. Gustincich, F. Persichetti, H. Suzuki, S. M. Grimmond, C. A. Wells, V. Orlando, C. Wahlestedt, E. T. Liu, M. Harbers, J. Kawai, V. B. Bajic, D. A. Hume, Y. Hayashizaki, Genome-wide analysis of mammalian promoter architecture and evolution. Nat. Genet. 38, 626–635 (2006).

21. A. C. Seila, J. M. Calabrese, S. S. Levine, G. W. Yeo, P. B. Rahl, R. A. Flynn, R. A. Young, P. A. Sharp, Divergent transcription from active promoters. Science. 322, 1849–1851 (2008).

22. L. J. Core, J. J. Waterfall, J. T. Lis, Nascent RNA sequencing reveals widespread pausing and divergent initiation at human promoters. Science. 322, 1845–1848 (2008).

23. S. H. C. Duttke, S. A. Lacadie, M. M. Ibrahim, C. K. Glass, D. L. Corcoran, C. Benner, S. Heinz, J. T. Kadonaga, U. Ohler, Human promoters are intrinsically directional. Mol. Cell. 57, 674–684 (2015).

24. R. Andersson, Y. Chen, L. Core, J. T. Lis, A. Sandelin, T. H. Jensen, Human Gene Promoters Are Intrinsically Bidirectional. Mol. Cell. 60 (2015), pp. 346–347.

25. S. H. C. Duttke, S. A. Lacadie, M. M. Ibrahim, C. K. Glass, D. L. Corcoran, C. Benner, S. Heinz, J. T. Kadonaga, U. Ohler, Perspectives on Unidirectional versus Divergent Transcription. Mol. Cell. 60 (2015), pp. 348–349.

26. H. Kawaji, M. Lizio, M. Itoh, M. Kanamori-Katayama, A. Kaiho, H. Nishiyori-Sueki, J. W. Shin, M. Kojima-Ishiyama, M. Kawano, M. Murata, N. Ninomiya-Fukuda, S. Ishikawa-Kato, S. Nagao-Sato, S. Noma, Y. Hayashizaki, A. R. R. Forrest, P. Carninci, FANTOM Consortium, Comparison of CAGE and RNA-seq transcriptome profiling using clonally amplified and single-molecule next-generation sequencing. Genome Res. 24, 708–717 (2014).

27. C.-C. Hon, J. A. Ramilowski, J. Harshbarger, N. Bertin, O. J. L. Rackham, J. Gough, E. Denisenko, S. Schmeier, T. M. Poulsen, J. Severin, M. Lizio, H. Kawaji, T. Kasukawa, M. Itoh, A. M. Burroughs, S. Noma, S. Djebali, T. Alam, Y. A. Medvedeva, A. C. Testa, L. Lipovich, C.-W. Yip, I. Abugessaisa, M. Mendez, A. Hasegawa, D. Tang, T. Lassmann, P. Heutink, M. Babina, C. A. Wells, S. Kojima, Y. Nakamura, H. Suzuki, C. O. Daub, M. J. L. de Hoon, E. Arner, Y. Hayashizaki, P. Carninci, A. R. R. Forrest, An atlas of human long non-coding RNAs with accurate 5’ ends. Nature. 543, 199–204 (2017).

28. J. Zhou, Sequence-based modeling of three-dimensional genome architecture from kilobase to chromosome scale. Nat. Genet. 54, 725–734 (2022).

29. M. I. Love, W. Huber, S. Anders, Moderated estimation of fold change and dispersion for RNA-seq data with DESeq2. Genome Biol. 15, 550 (2014).

30. K. Katoh, D. M. Standley, MAFFT multiple sequence alignment software version 7: improvements in performance and usability. Mol. Biol. Evol. 30, 772–780 (2013).

