## Supplementary figures and images for "Sequence basis of transcription initiation in human genome"

### Supplementary Data 2

TATA

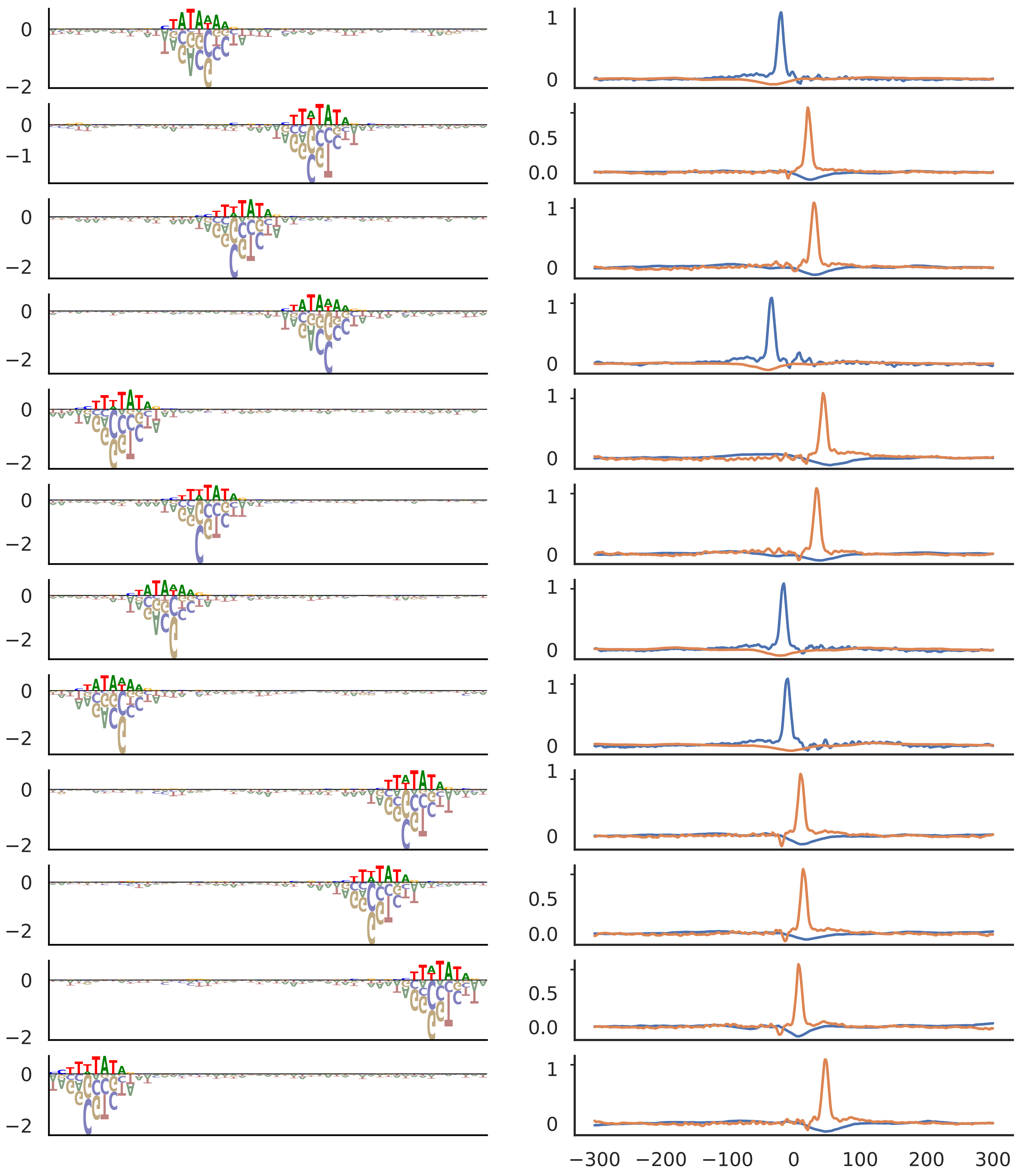

YY1

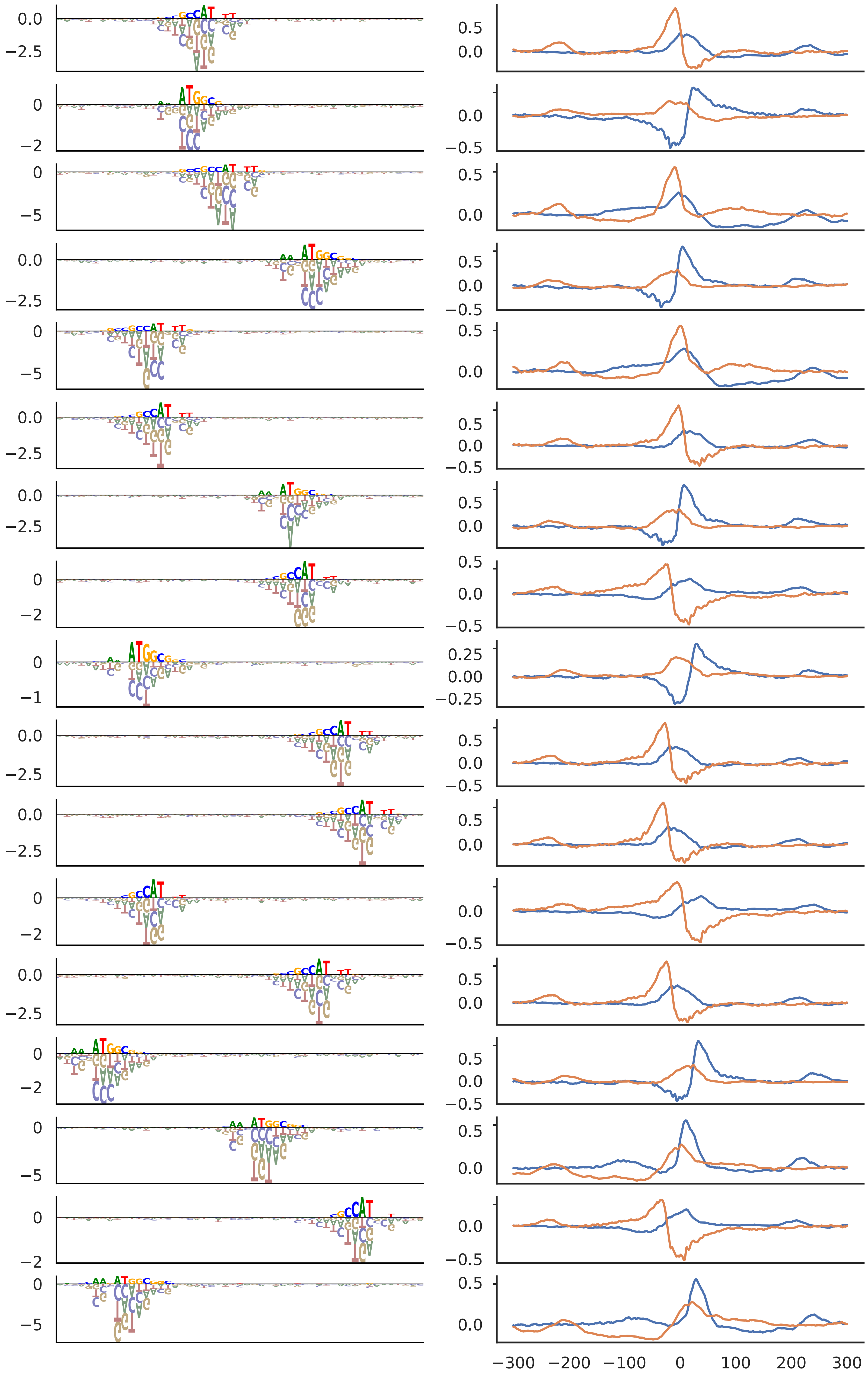

NFY

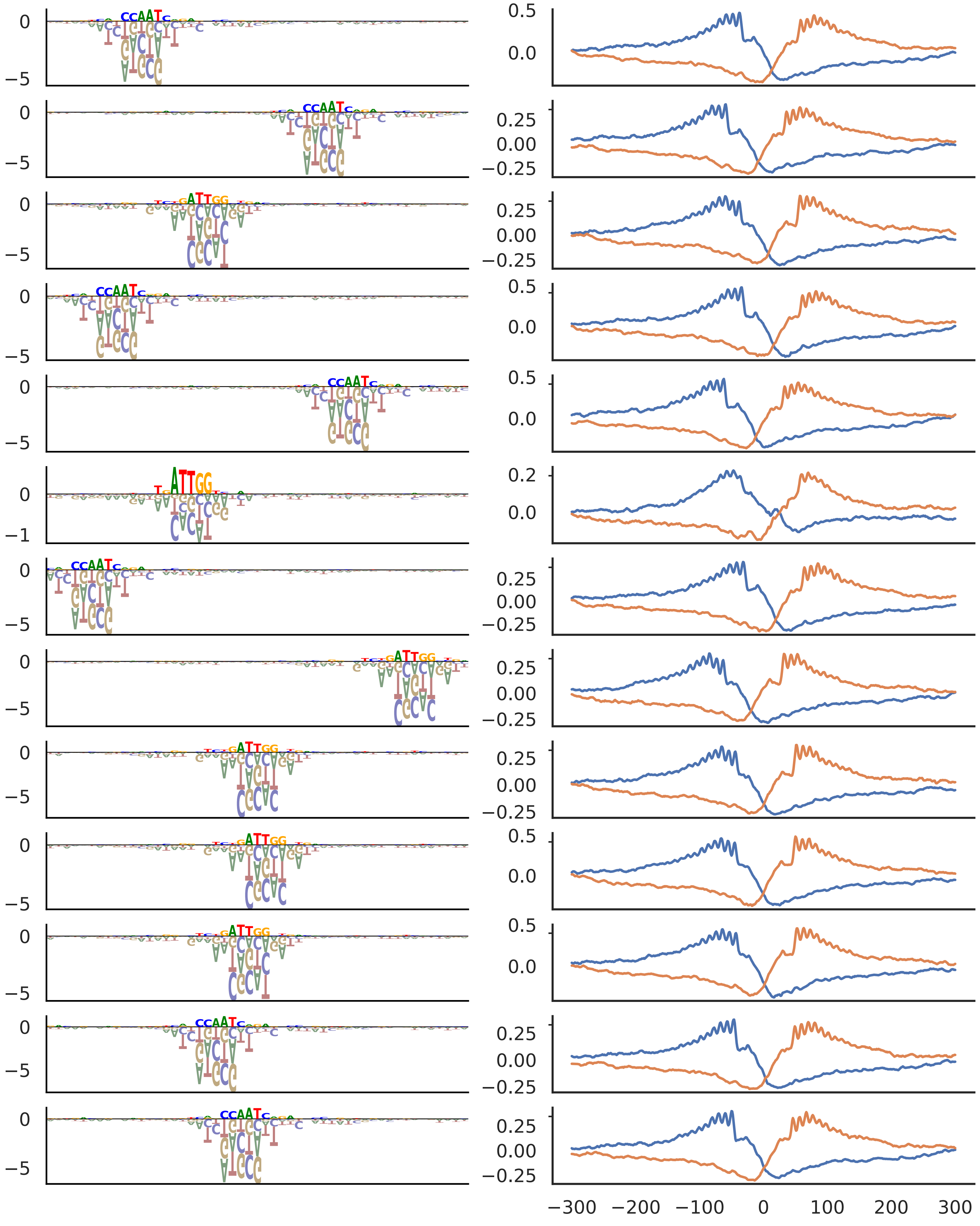

ETS

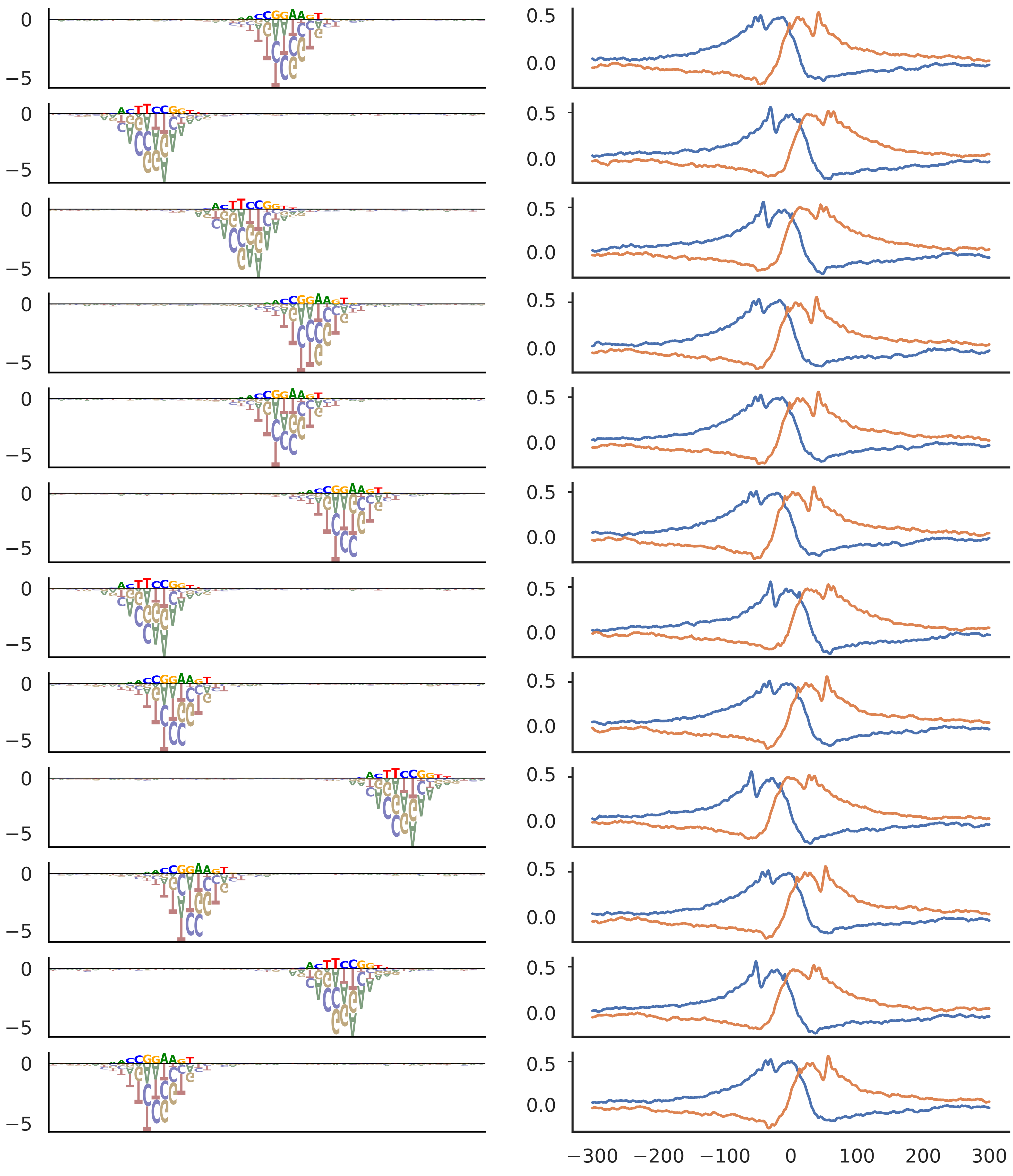

SP

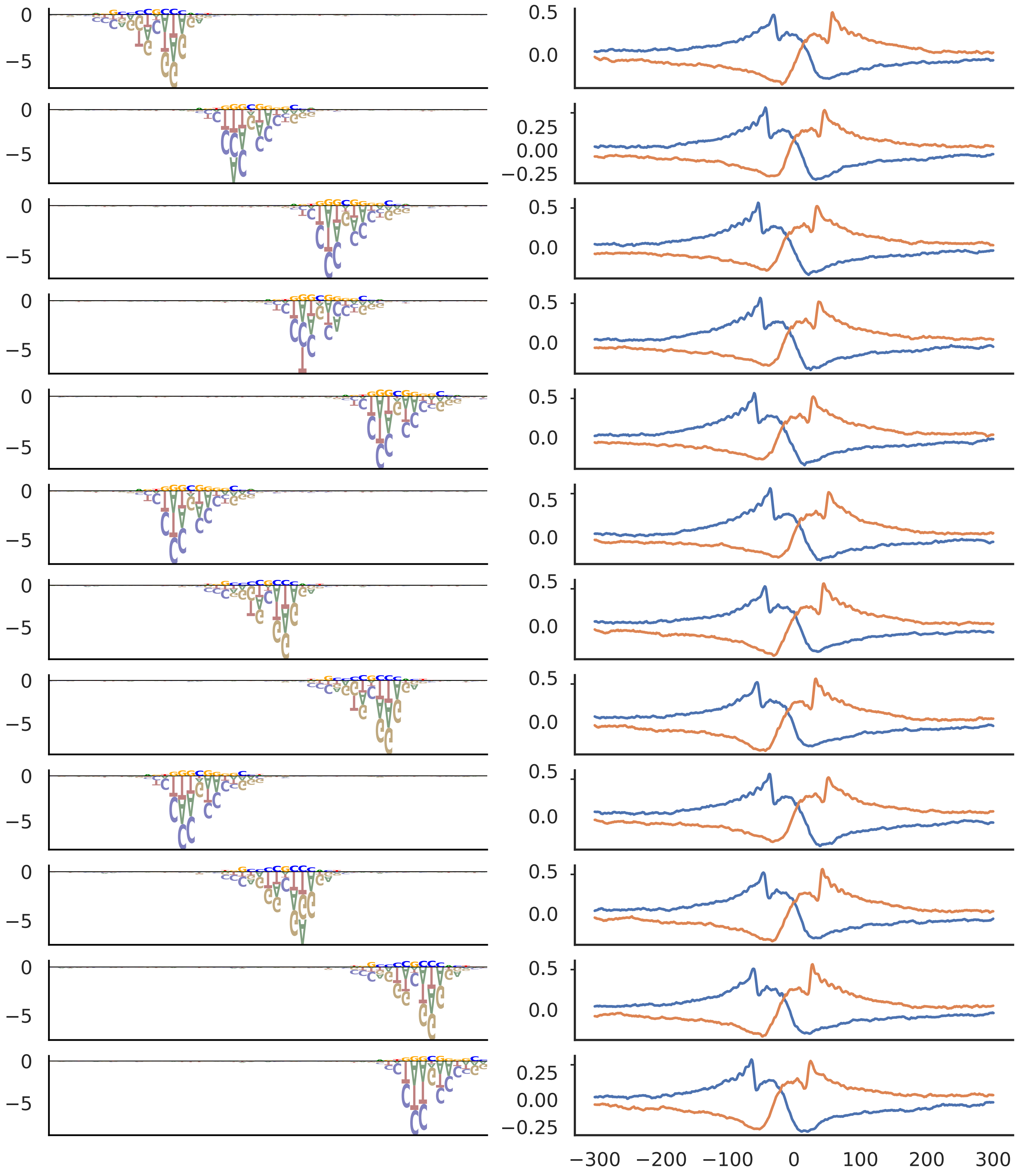

ZNF143

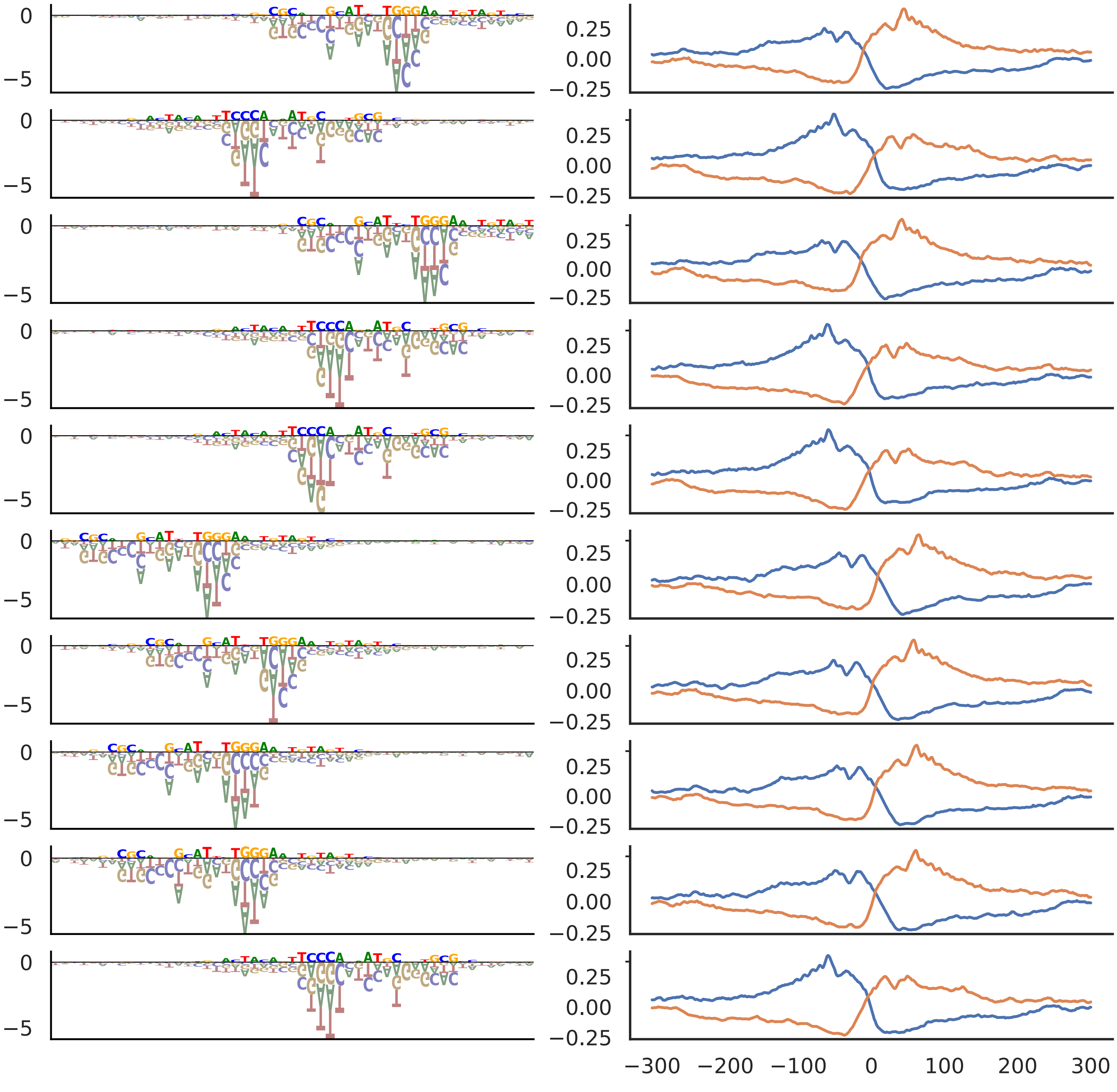

NRF1

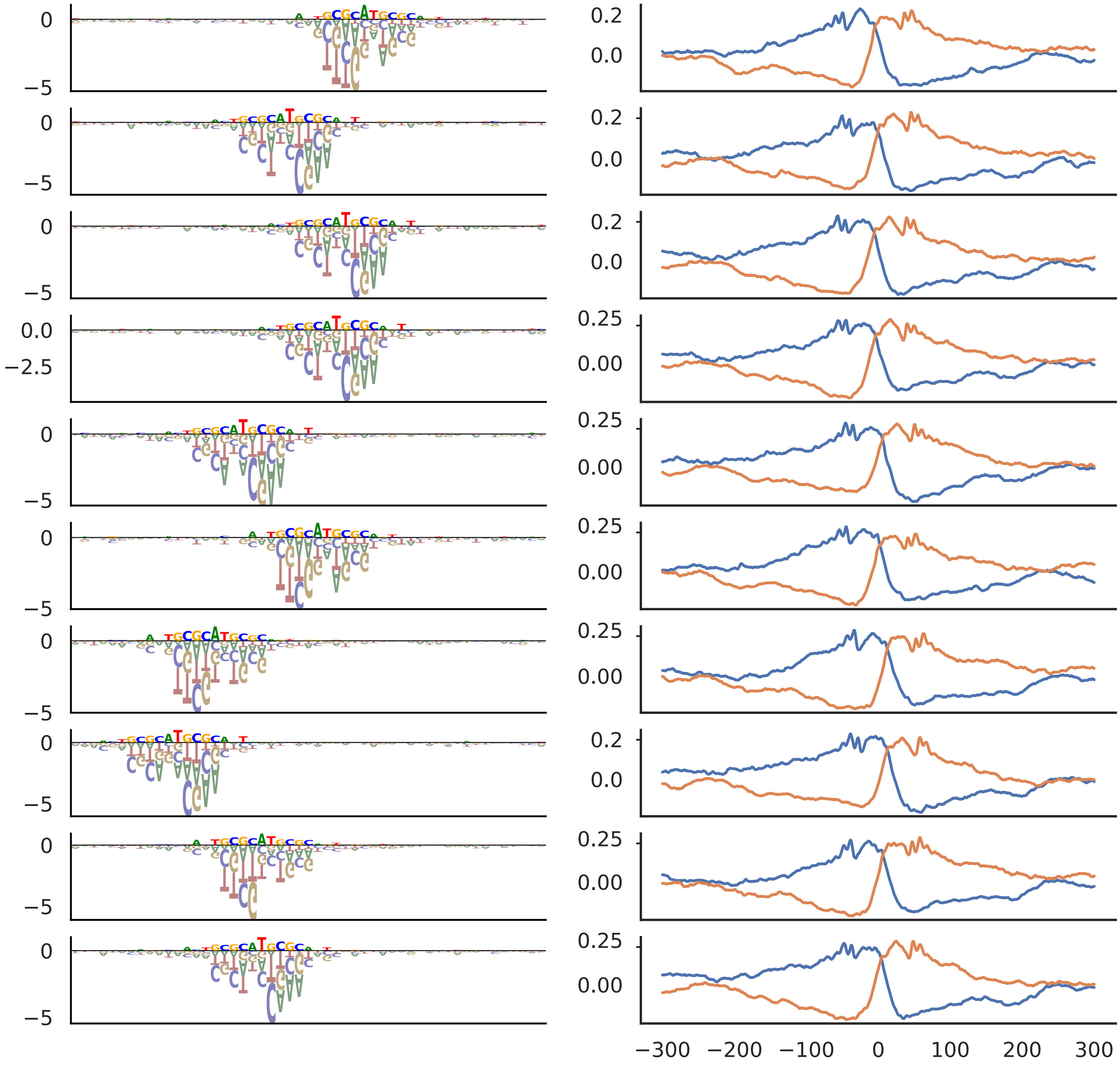

CREB

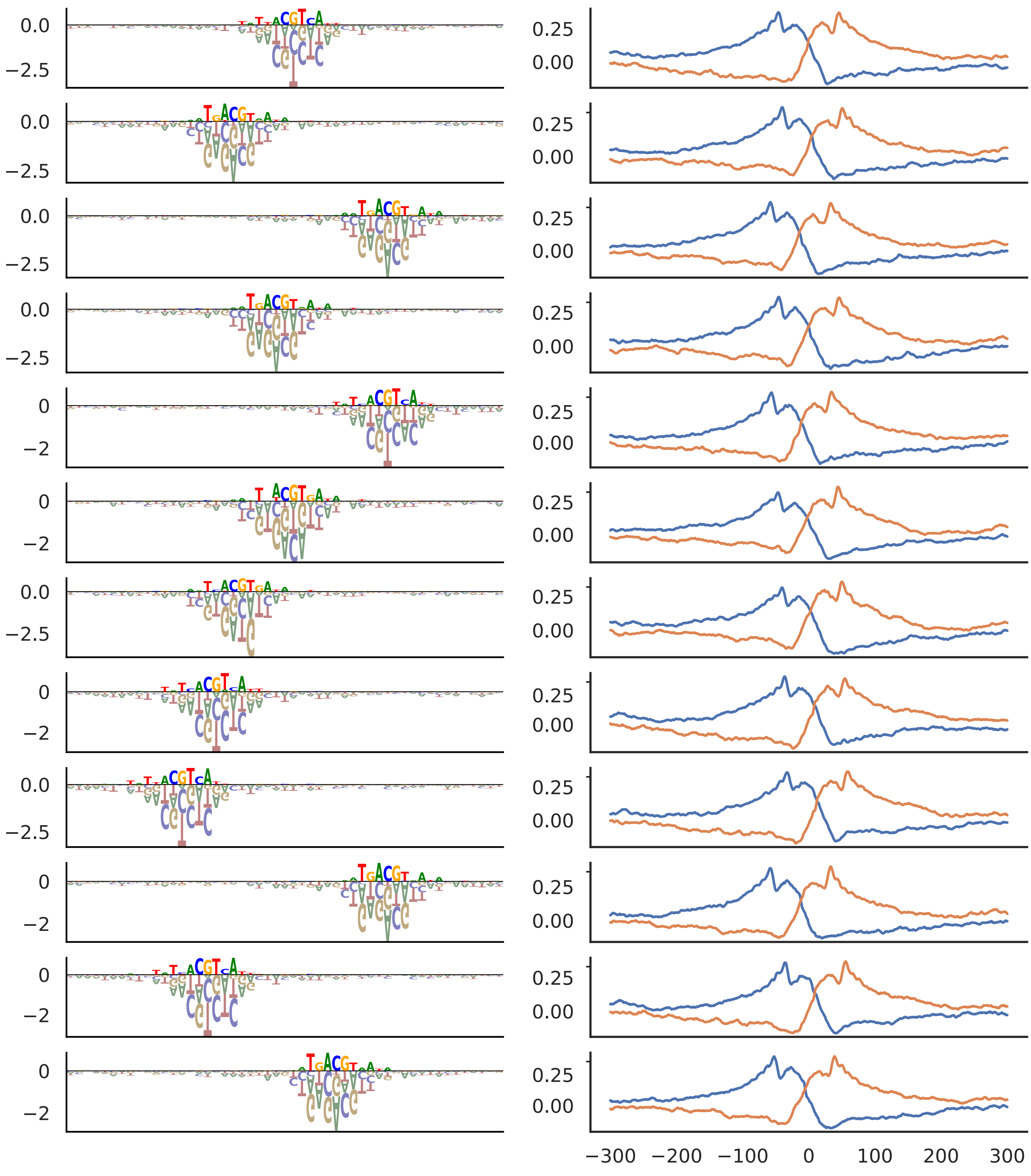

U1 snRNP

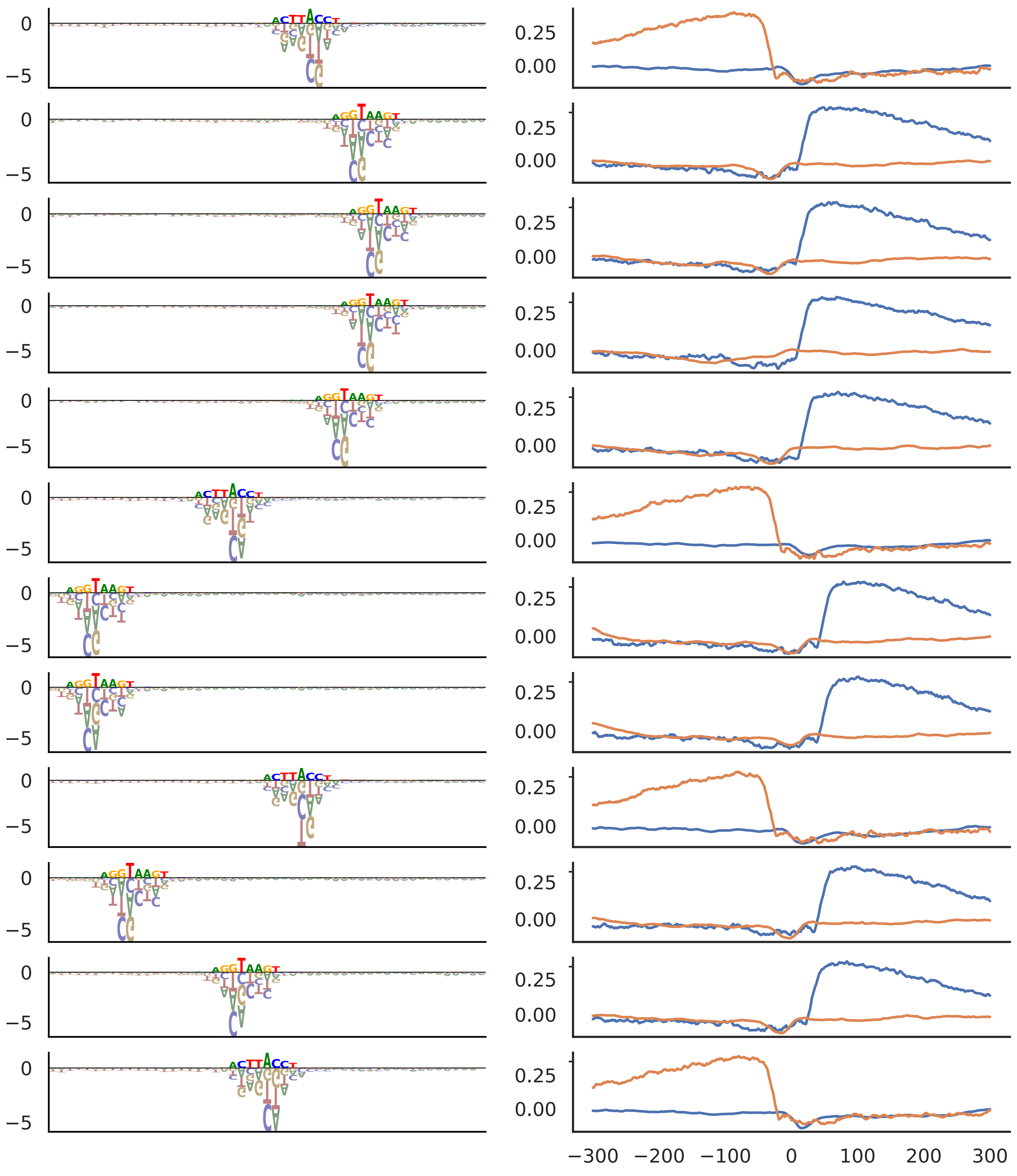

#### Long Inr

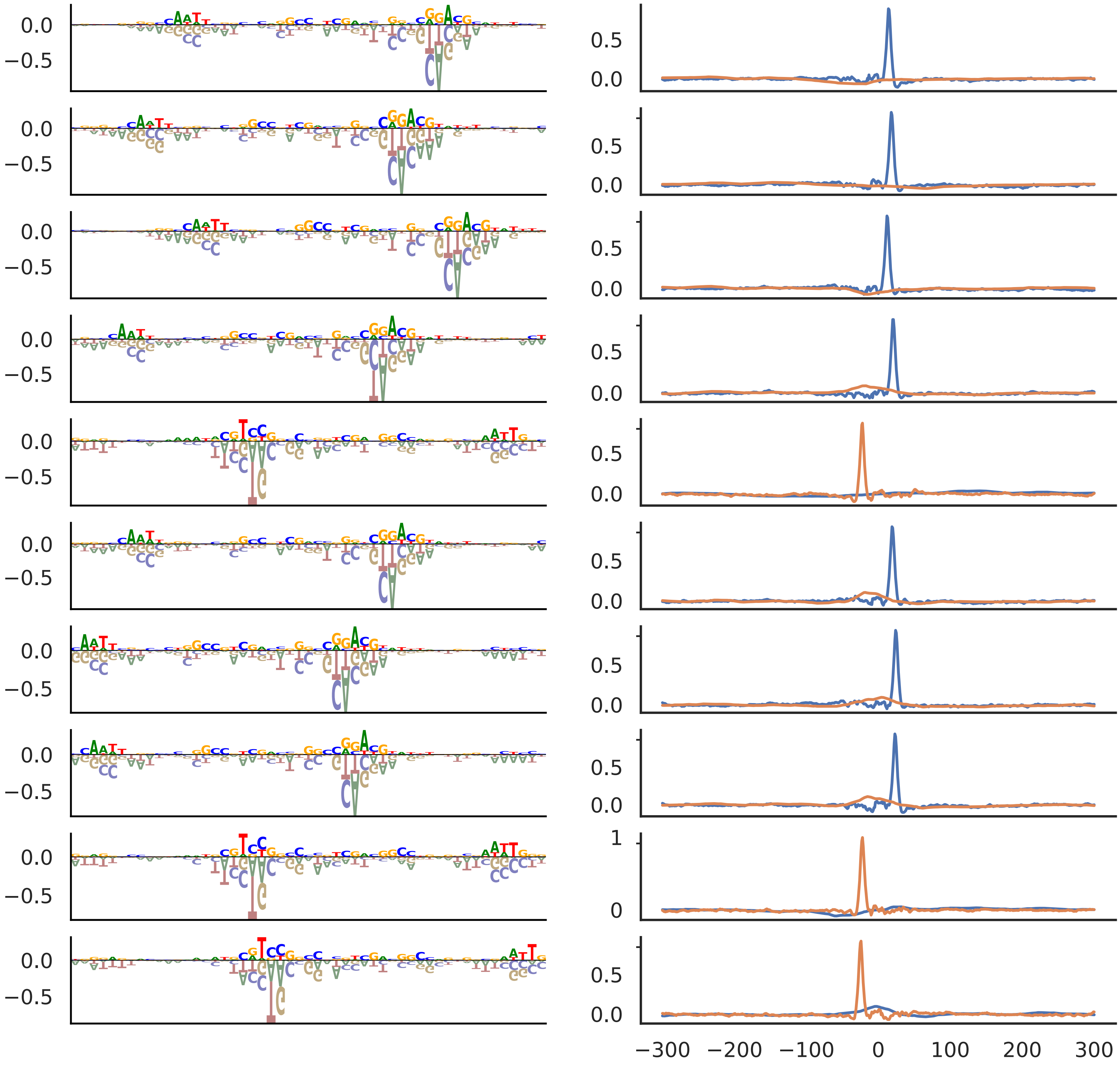

### Supplementary Data 3

TATA

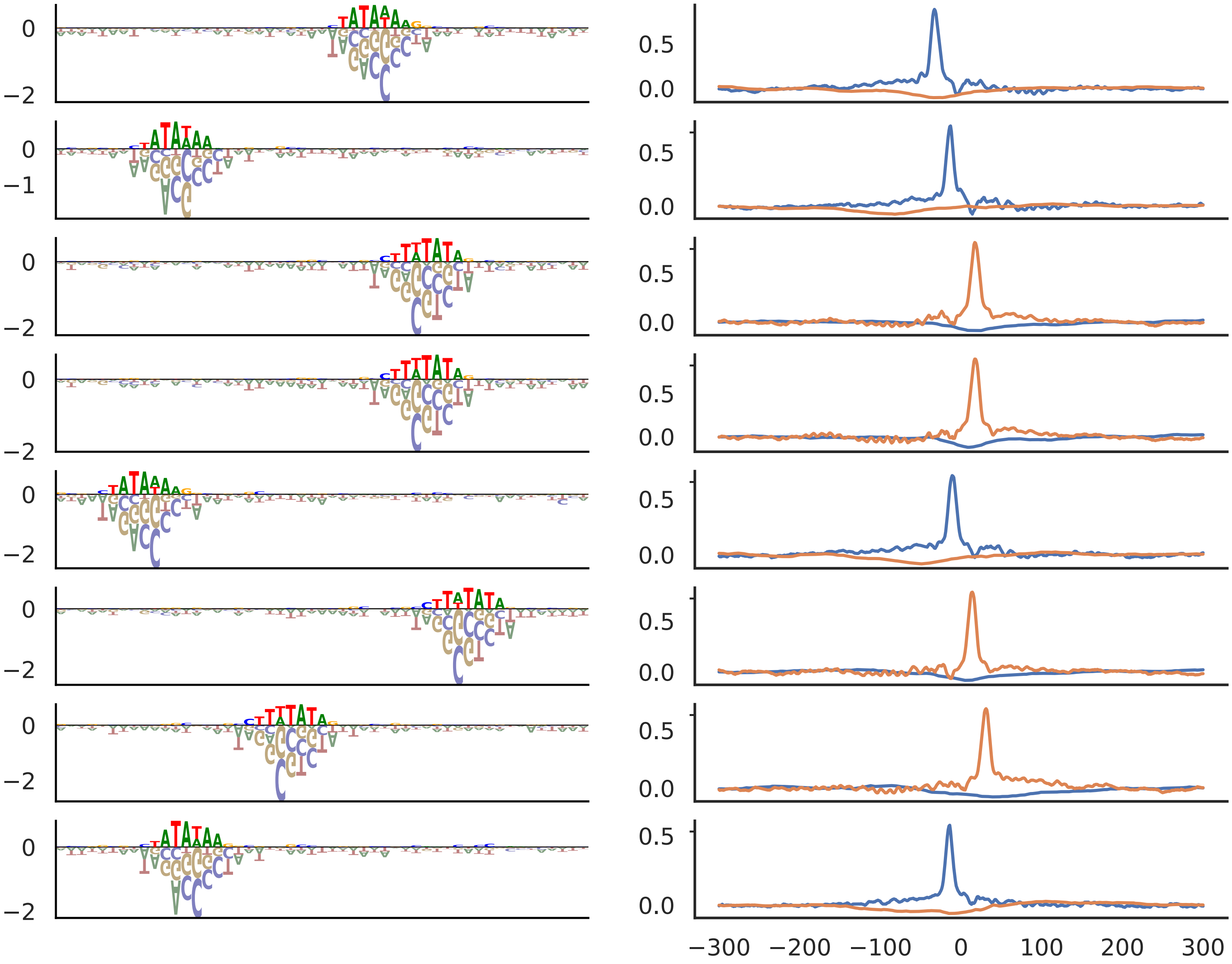

YY1

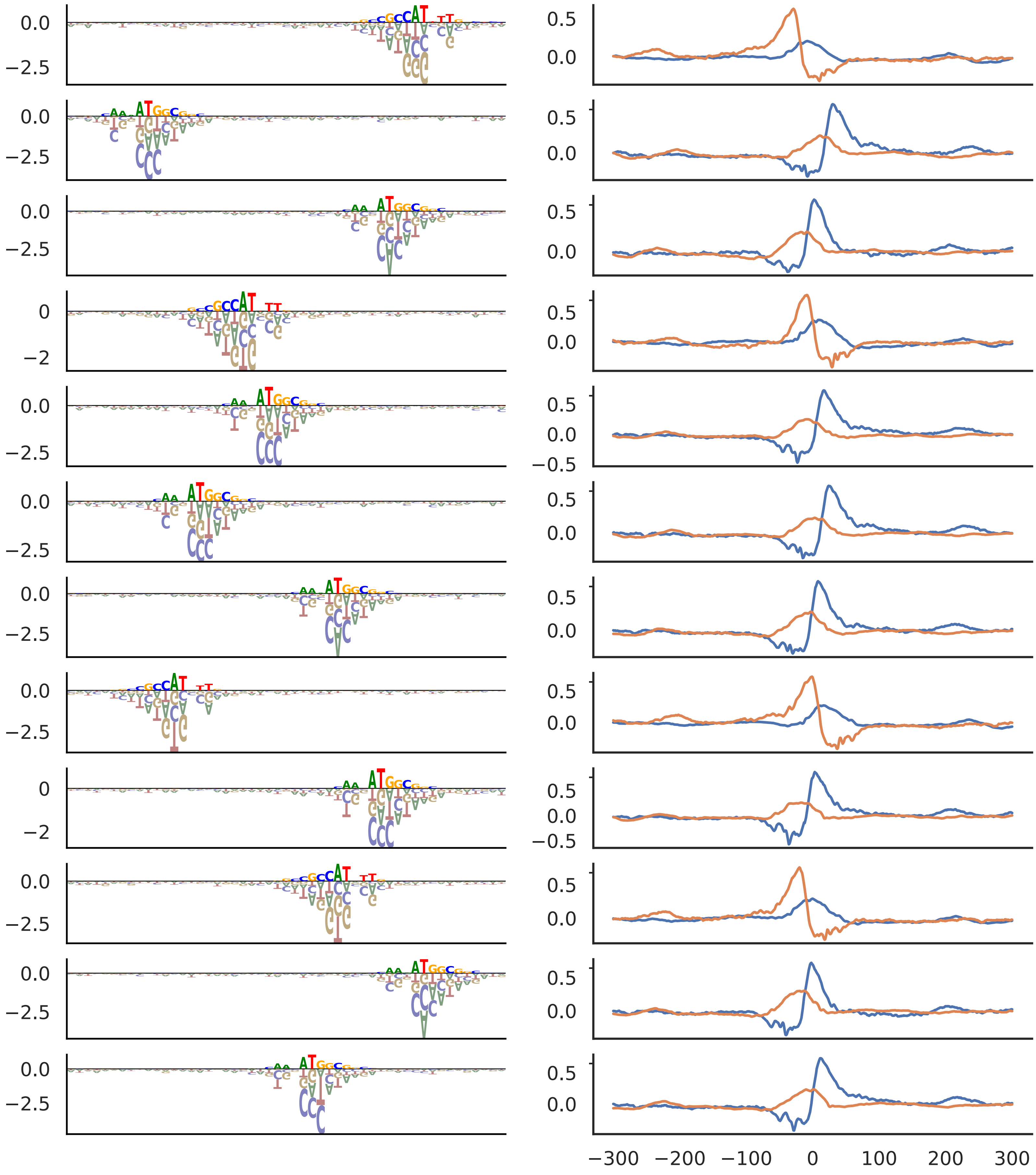

NFY

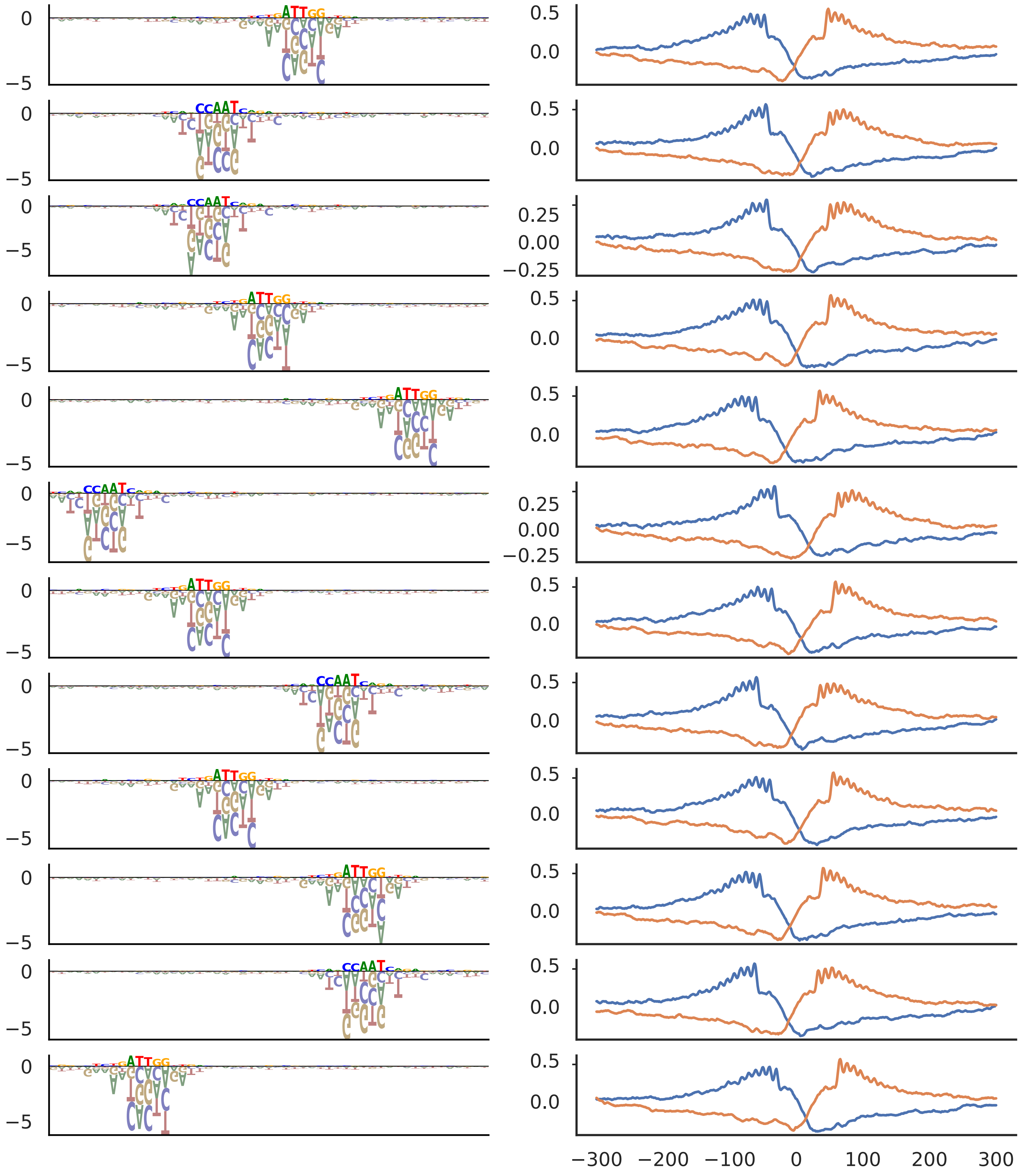

ETS

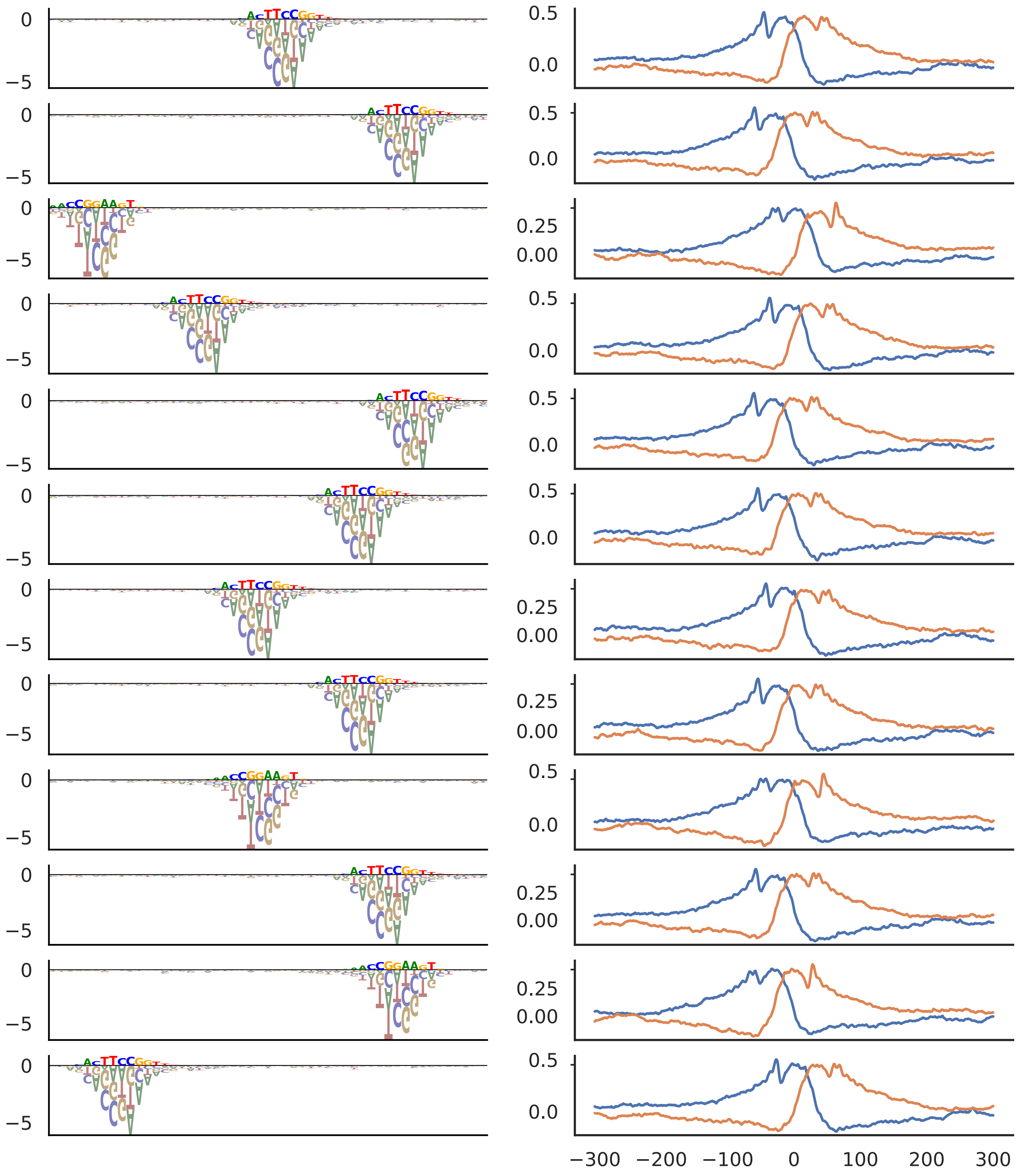

SP

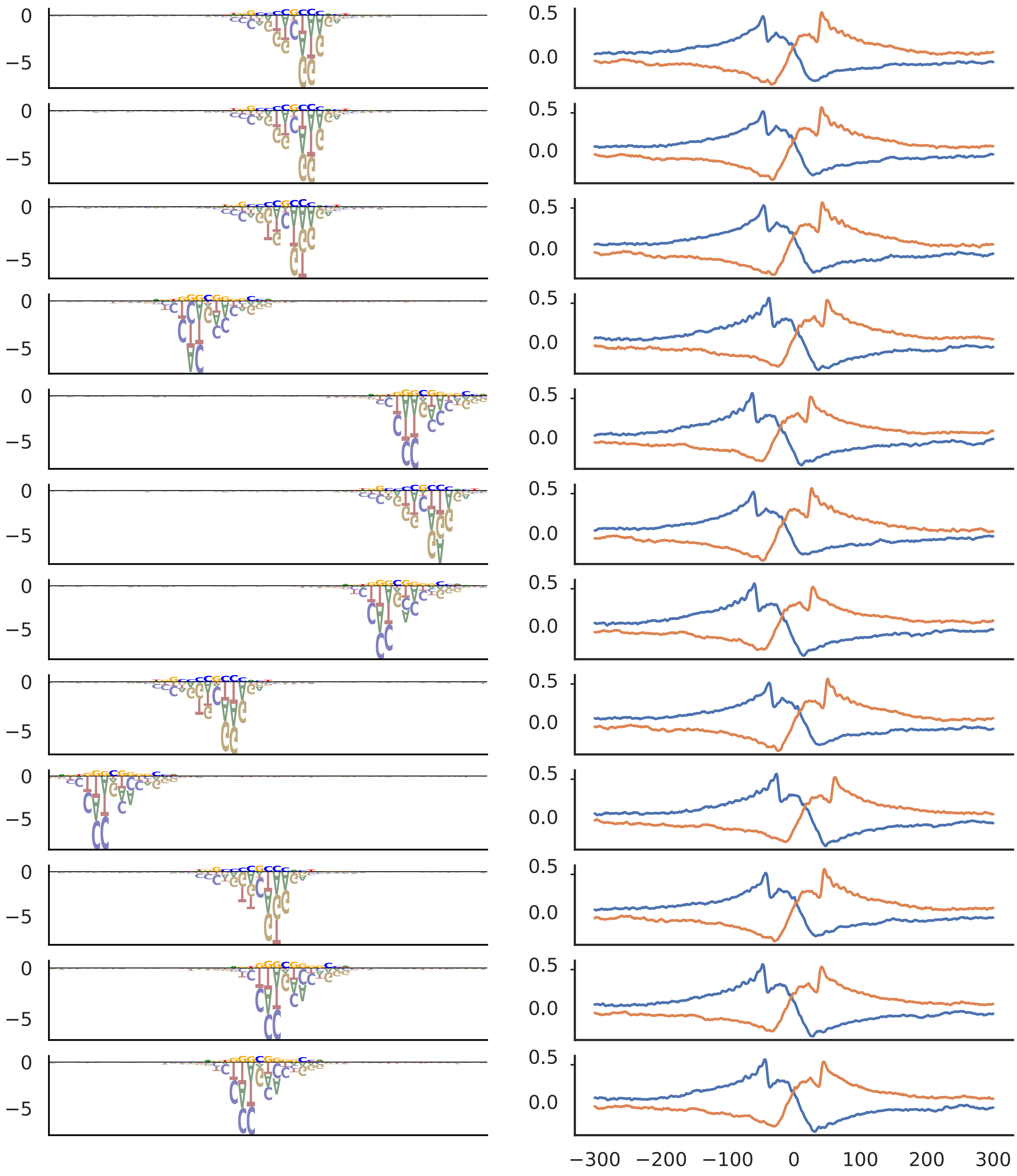

ZNF143

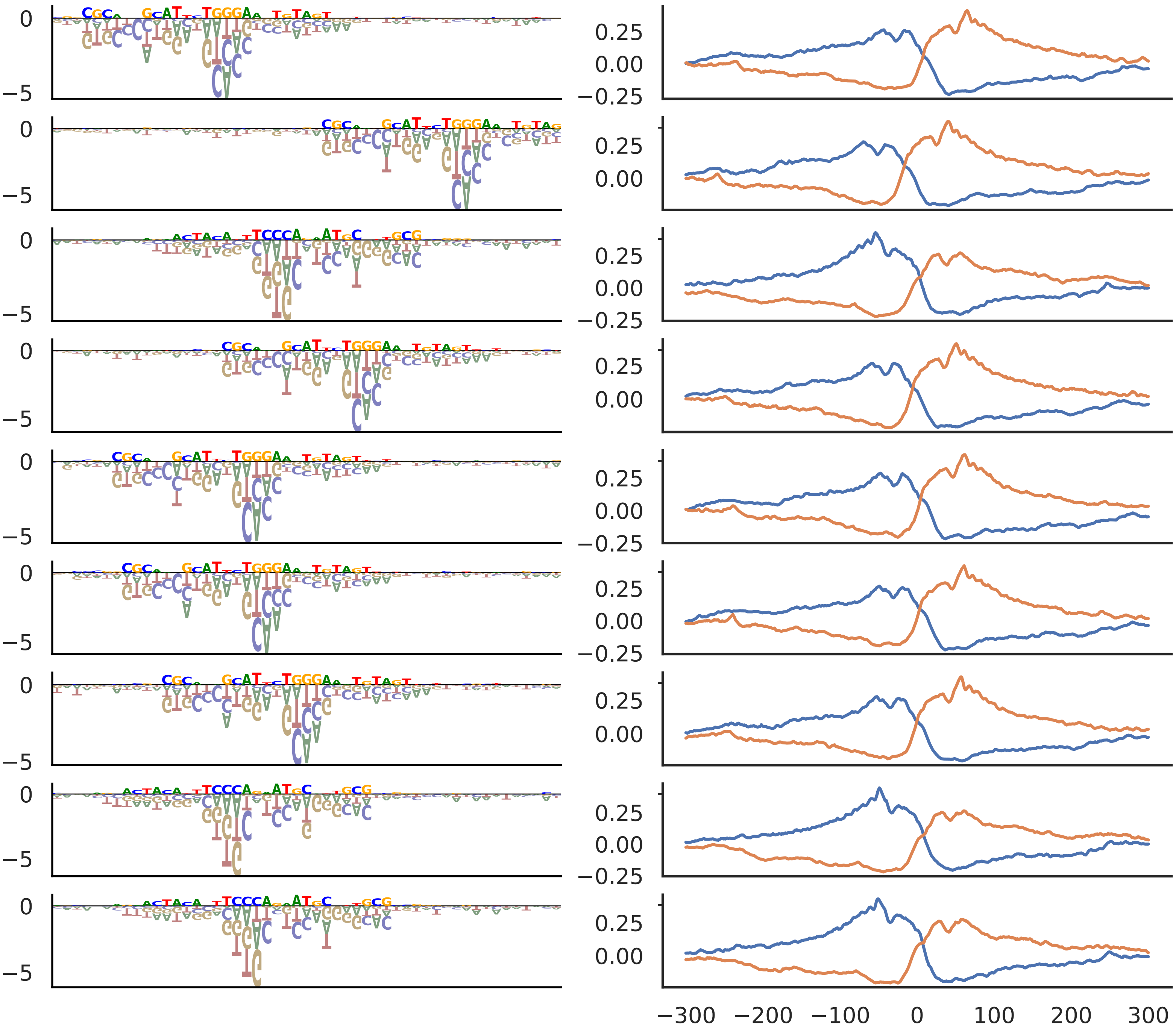

NRF1

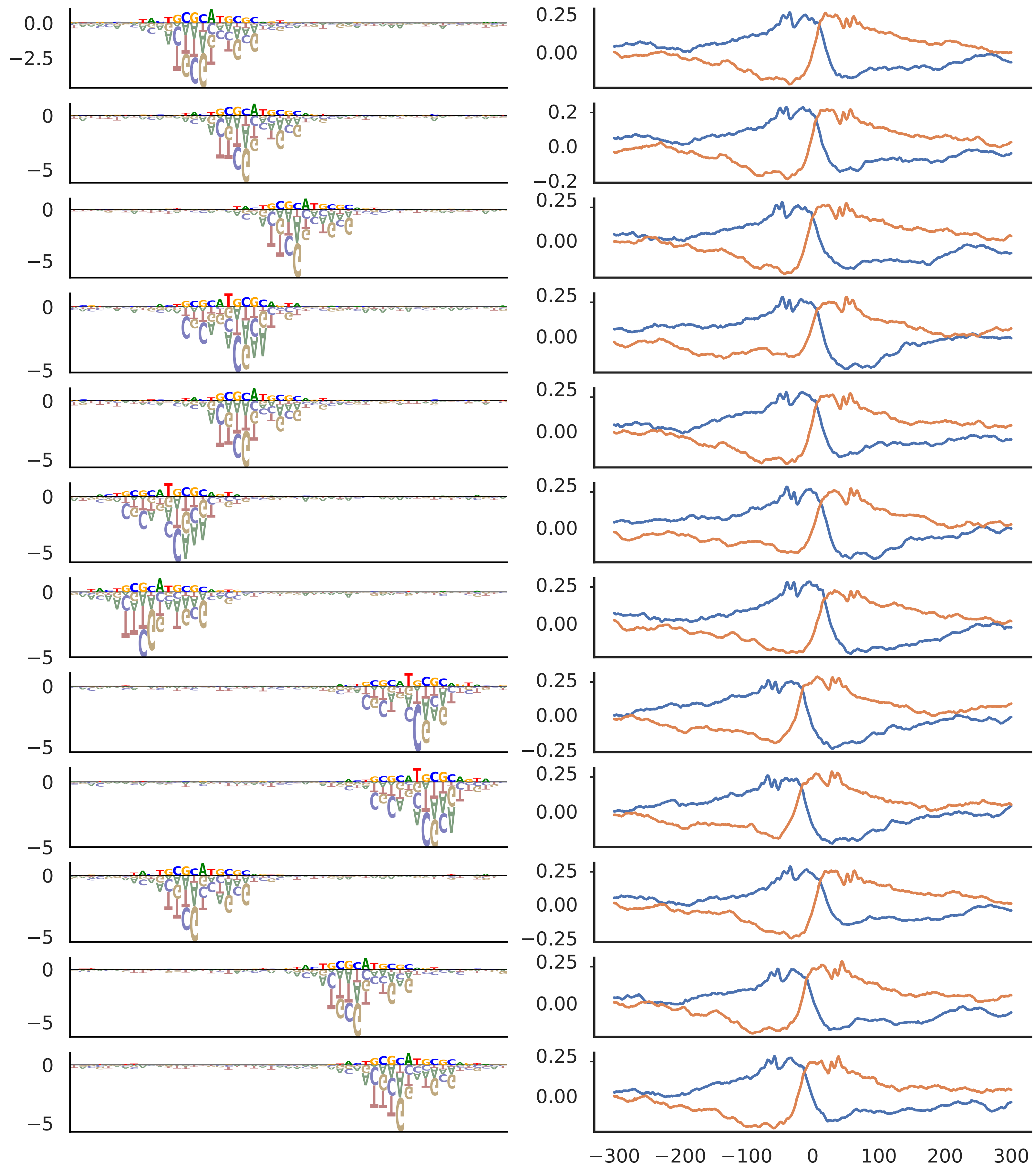

CREB

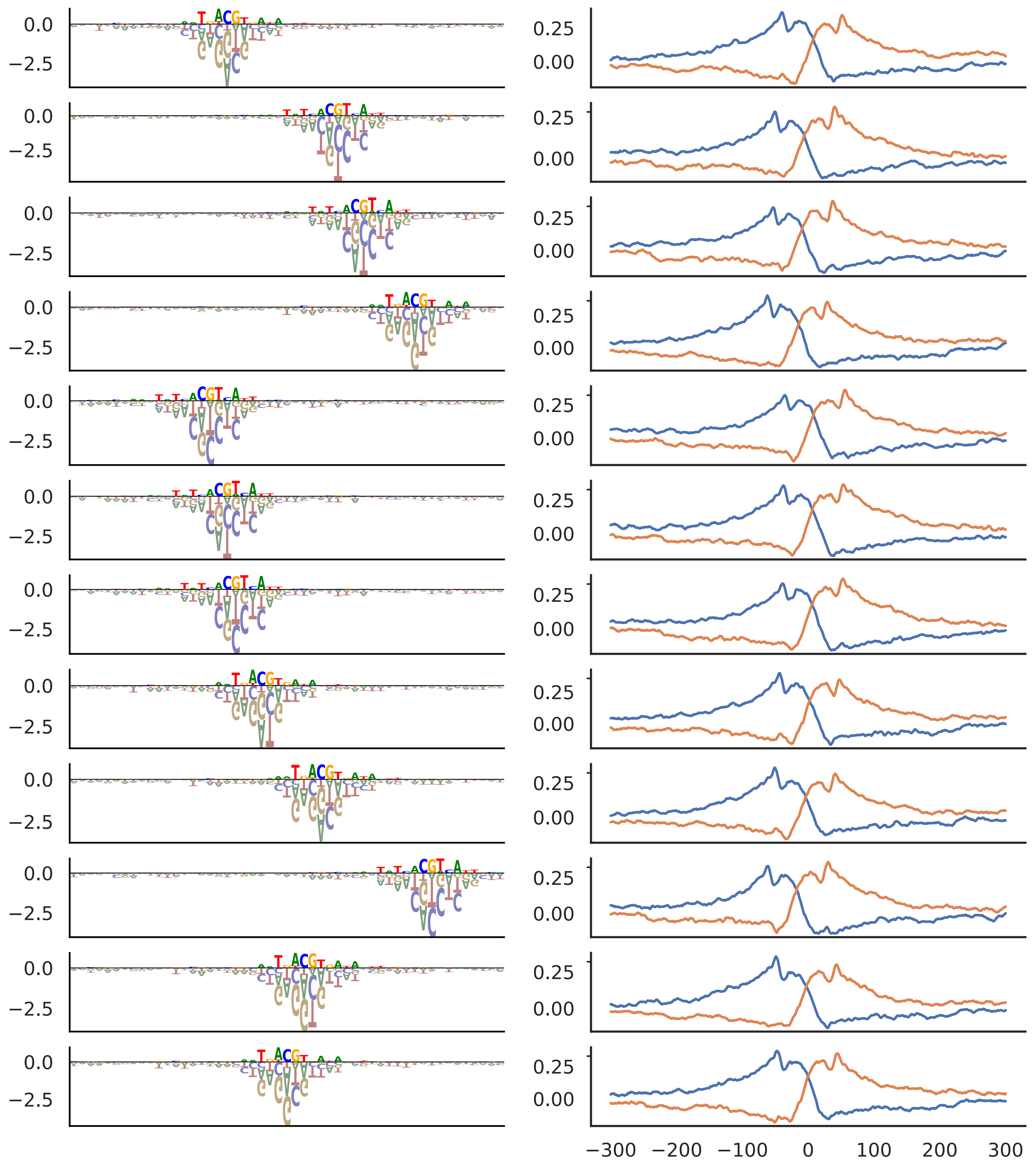

U1 snRNP

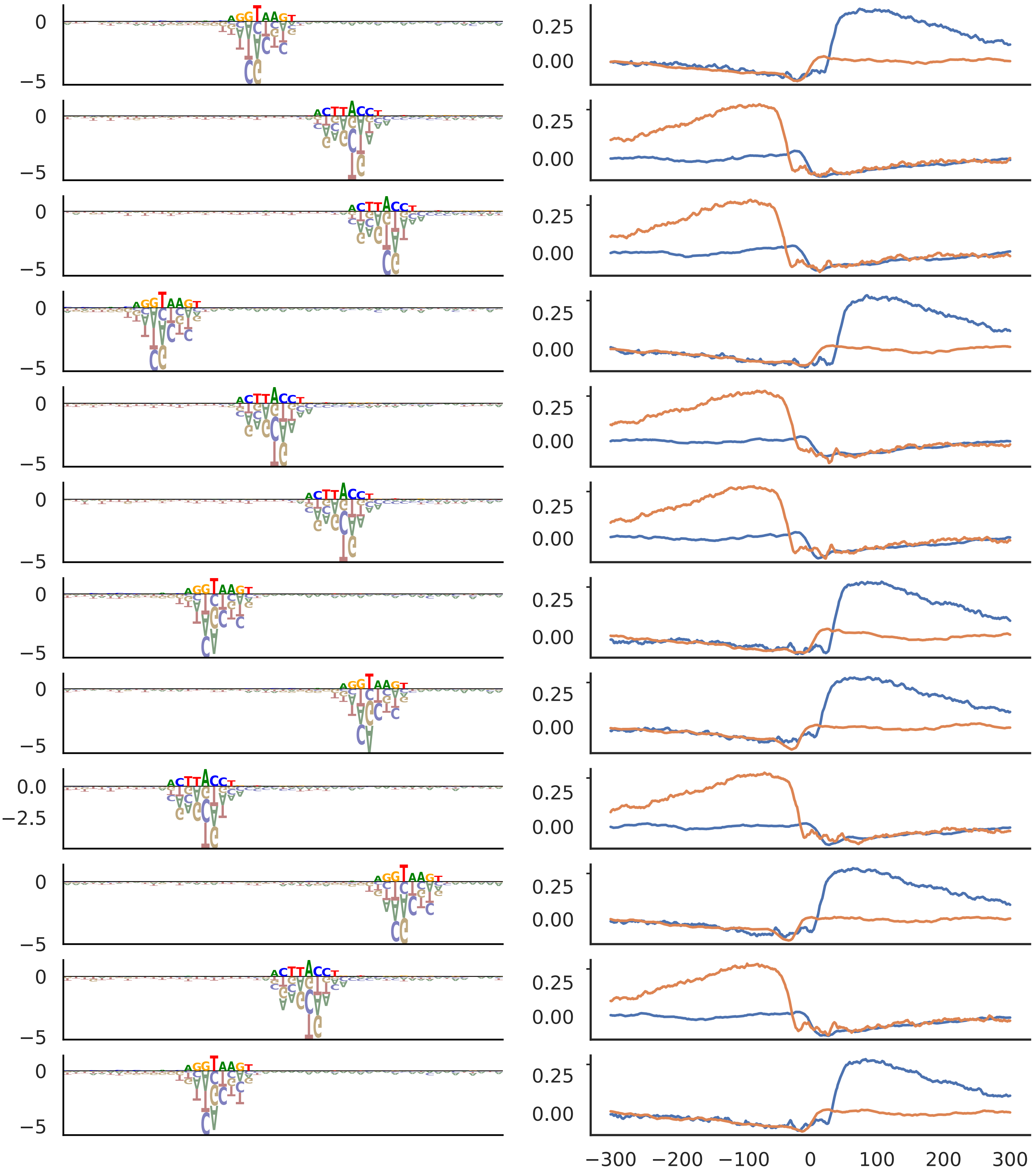

Long Inr

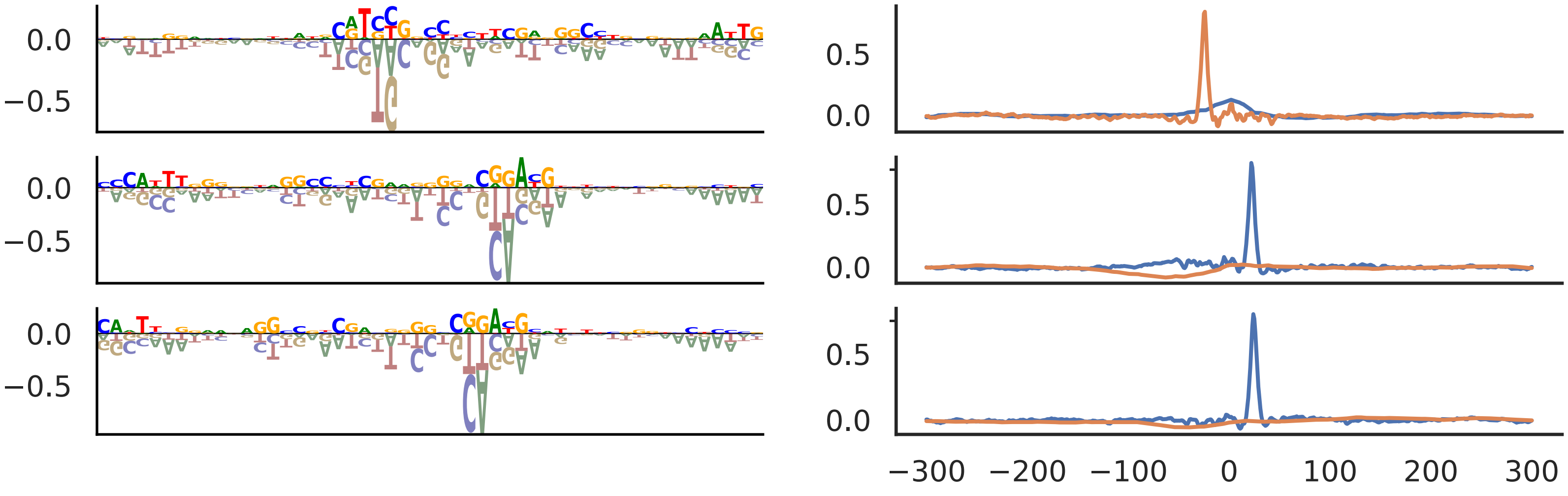
