## Supplementary materials for "Sequence basis of transcription initiation in human genome"

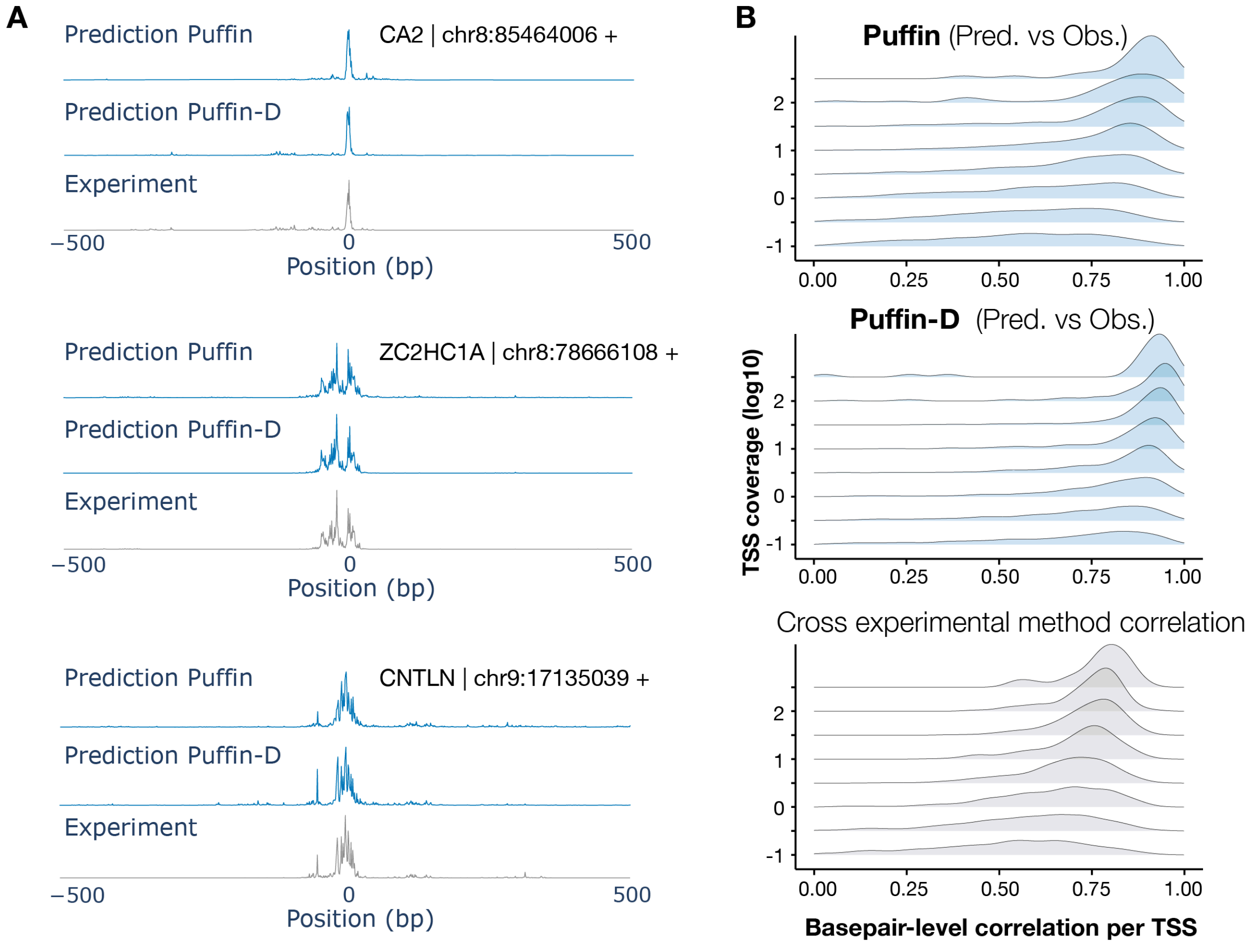


**Fig. S1. Puffin-D prediction performance on basepair-level transcription initiation signal near TSS.** (**A**) Example prediction of basepair resolution transcription initiation signal from promoter sequences on holdout chromosomes. The x-axis indicates basepair position relative to the annotated transcription start site and the y-axis is shown in log scale. (**B**) Basepair-level correlation (x-axis) between Puffin (top panel) or Puffin-D (middle panel) prediction and experimental measurement (FANTOM CAGE) within 1kb window of each annotated TSS. Puffin performance and cross-experimental method correlations are reproduced from Figure 1c-d for comparison.

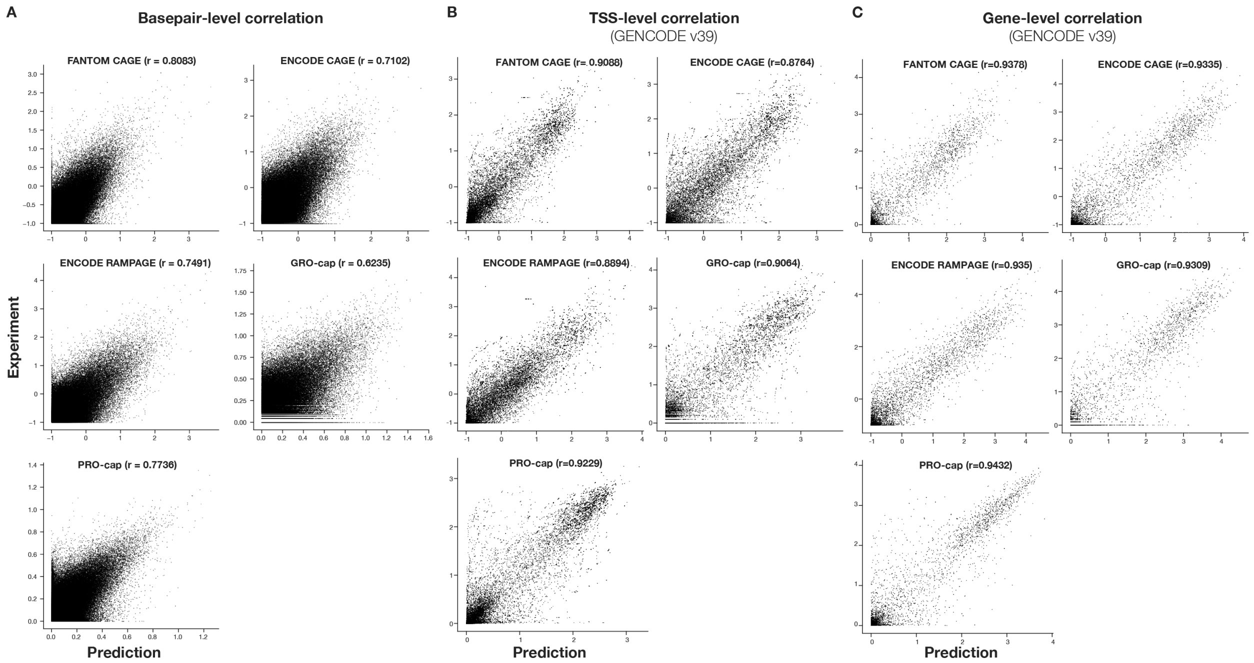


**Fig. S2. Sequence-based prediction of transcription initiation signal strength by Puffin-D.** The Puffin-D prediction was compared with experimental data at (**A**) basepair-level (each dot represents a basepair), (**B**) TSS-level (each dot represents a TSS defined as 400bp window surrounding the annotated TSS position), and (**C**) gene-level (each dot represents a gene, defined as the sum of all TSS for the gene) on test chromosomes, for five experimental techniques. The prediction (x-axis) and experimental data (y-axis) are both represented in log scale with pseudocount 0.1 for FANTOM CAGE, ENCODE CAGE, and ENCODE RAMPAGE and 1 for GRO/PRO-cap.

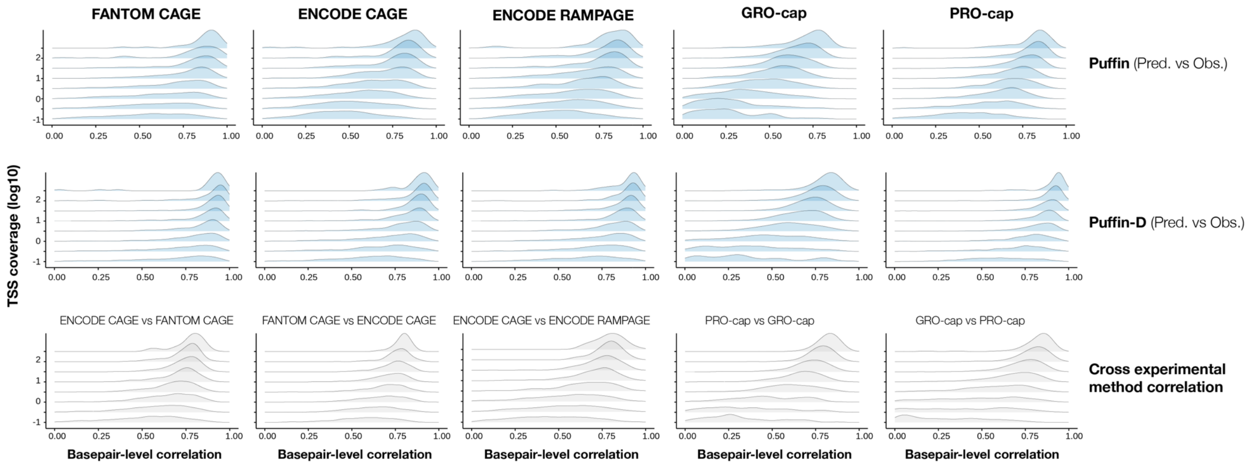


**Fig. S3. Puffin and Puffin-D prediction of transcription initiation signals achieve high correlation with experiment measurements at basepair-level.** Basepair-level correlation (x-axis) between Puffin (top panels) or Puffin-D (middle panels) predictions and experimental measurements within 1kb windows centered at annotated TSS. TSS were grouped by coverage level and the y-axis indicates the lower bound of each group (the upper bound of a group is the lower bound of the next group). Each column shows the performance of a different Puffin / Puffin-D prediction target. The bottom panels show the correlations between experimental measurement and its most correlated technique, which is indicated in the labels. The bottom panels provide a reference for the expected decrease in correlation due to lower coverage.

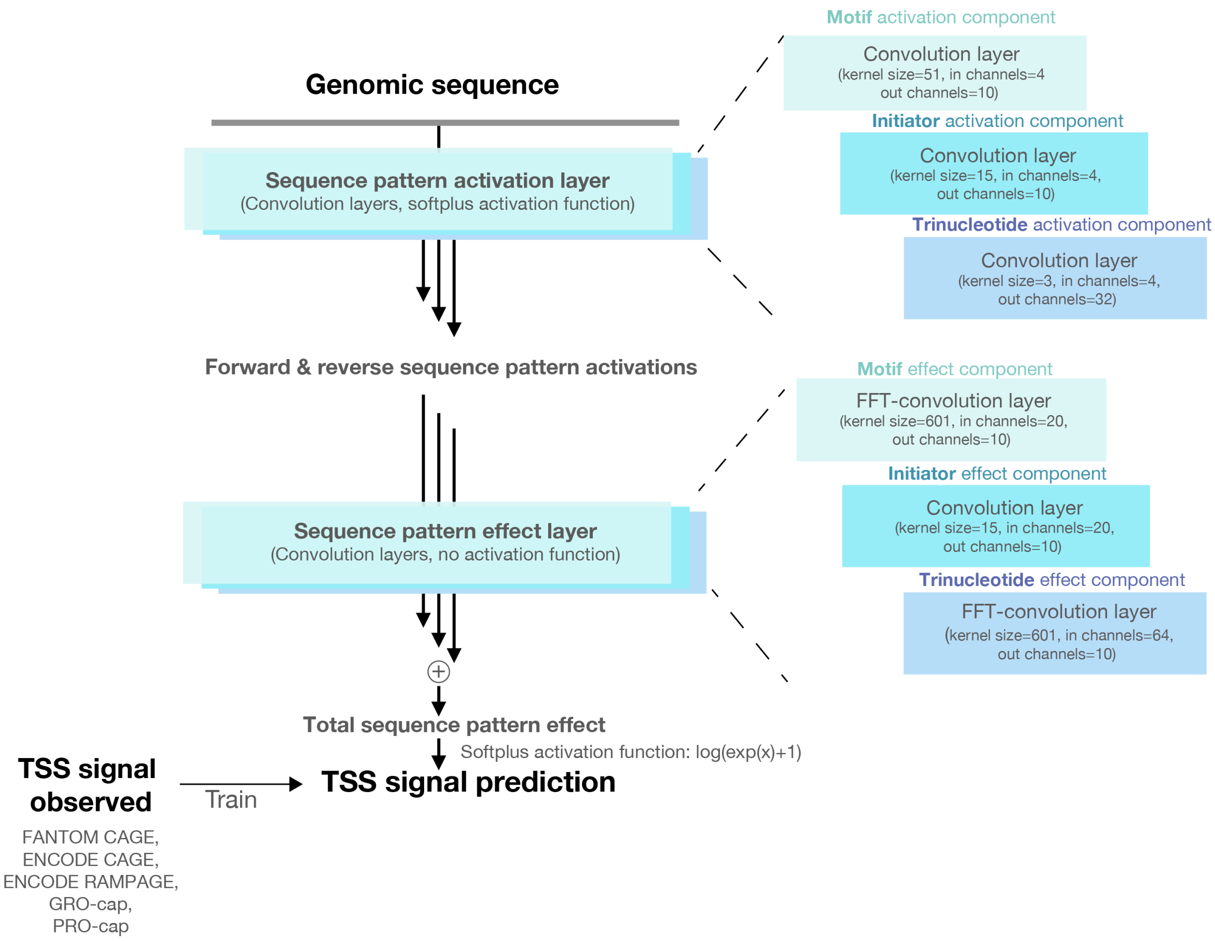


**Fig. S4. Schematic illustration of Puffin model architecture.** The architecture contains two learnable layers, corresponding to detection of sequence pattern and computing sequence pattern effects on transcription initiation respectively. Each layer has three components corresponding to motif, initiator, and trinucleotide.

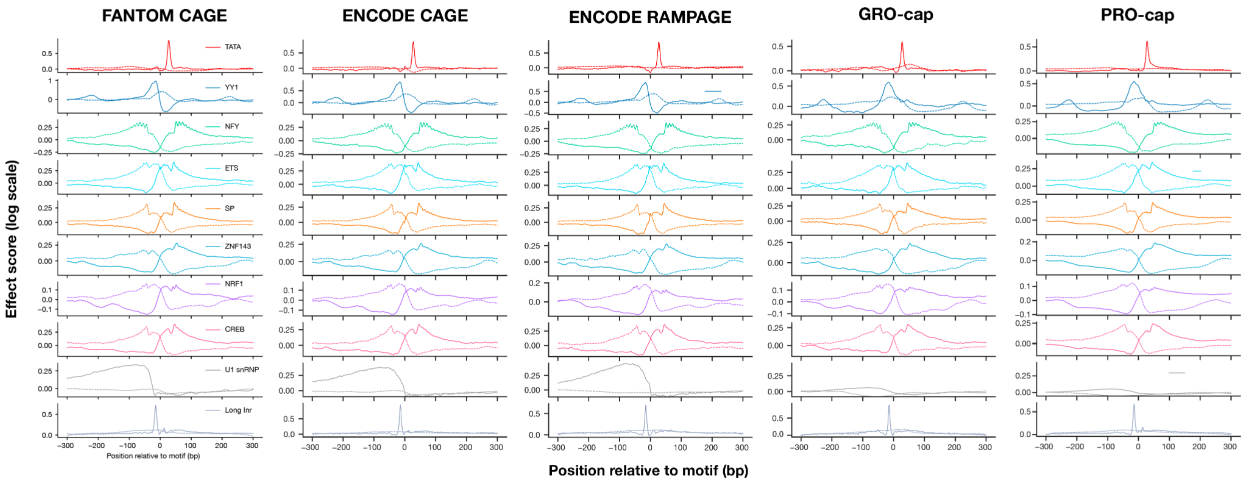


**Fig. S5. Position-specific motif effects estimated for all five experimental methods.** The solid lines show the transcription initiation effects on the forward strand and the dotted lines show the reverse strand. Notably the U1 snRNP motif only has strong effects on FANTOM CAGE, ENCODE CAGE, and RAMPAGE but not PRO/GRO-cap.

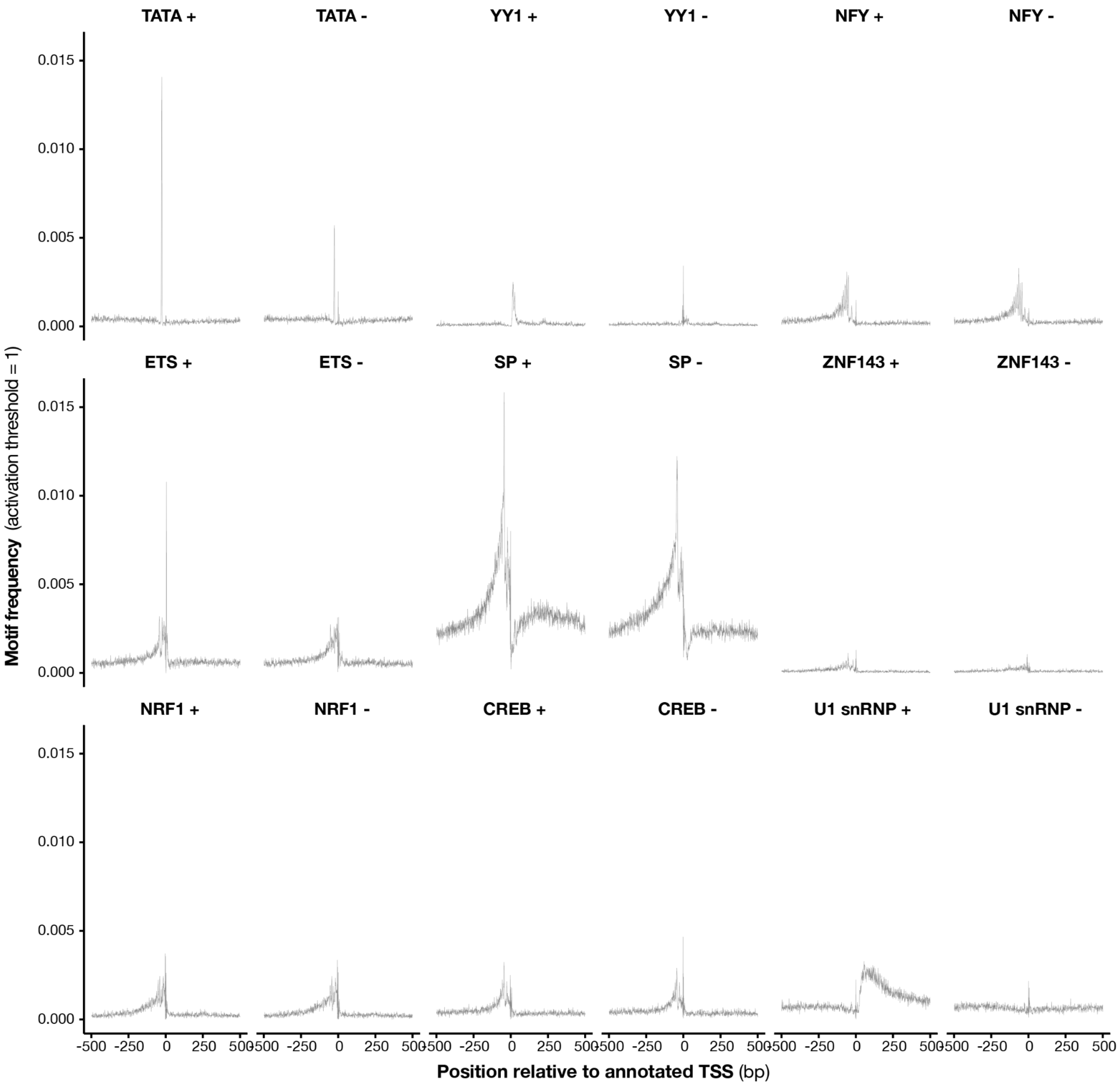


**Fig. S6. Motif activation frequencies at each basepair position near TSS.** For each motif type, the frequency of motif activation > 1 (y-axis) at each basepair position relative to TSS (x-axis) across the top 40,000 TSS ranked by FANTOM CAGE signals is shown.

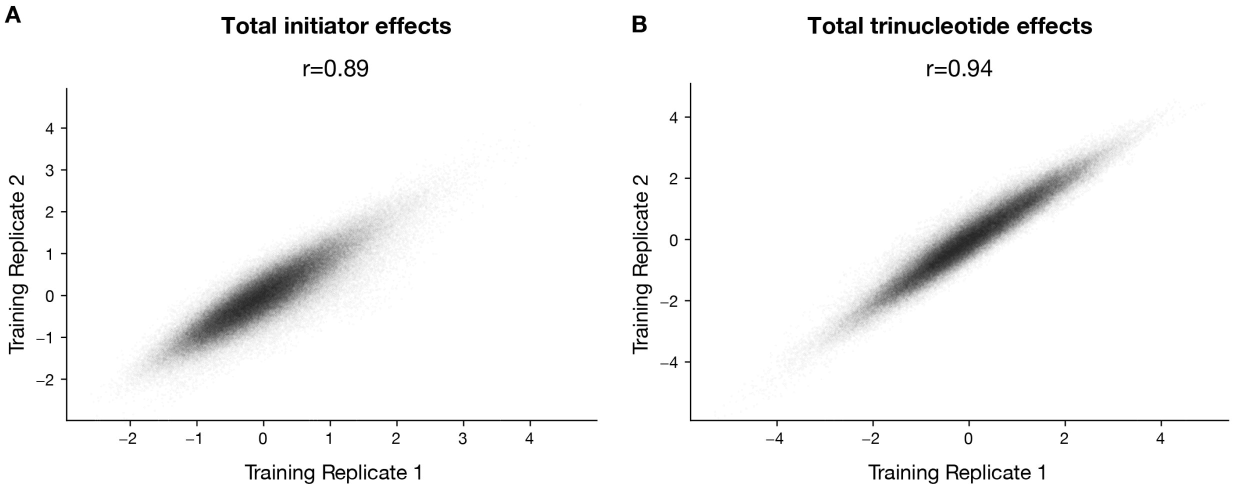


**Fig. S7. Total initiator and trinucleotide sequence pattern effects are highly reproducible across training replicates.** The total initiator effects (**A**) or trinucleotide effects (**B**) per basepair are highly reproducible across training replicates with different random seeds. A random subset of 100 TSS (1000 x 100 bps) among the top 40,000 TSS ranked by FANTOM CAGE signals was shown in the scatter plots.

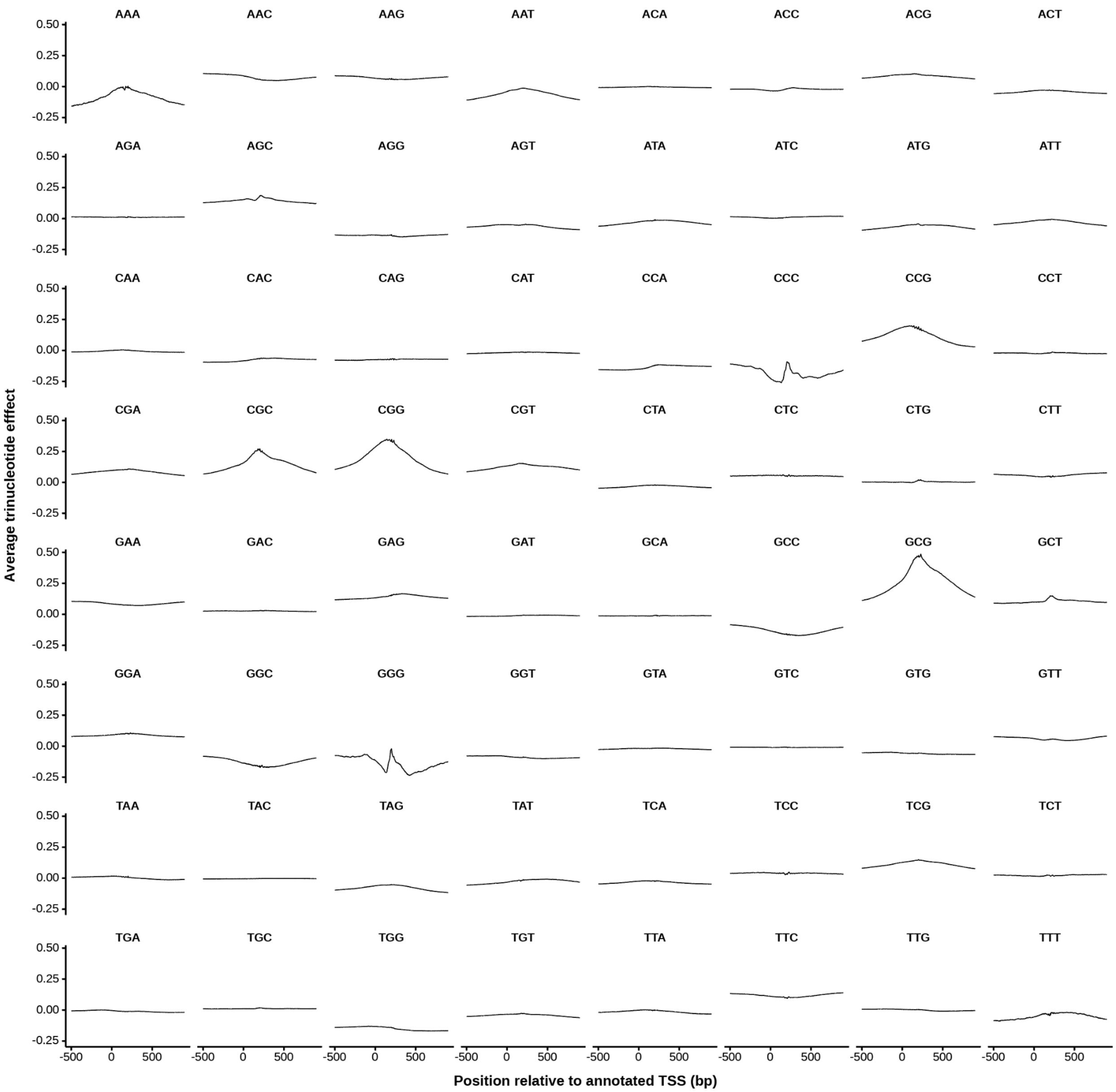


**Fig. S8. Average trinucleotide contribution to TSS for each trinucleotide type.** Average trinucleotide effects estimated by Puffin for each of the 64 trinucleotides across the top 40,000 TSS ranked by FANTOM CAGE signals were computed.

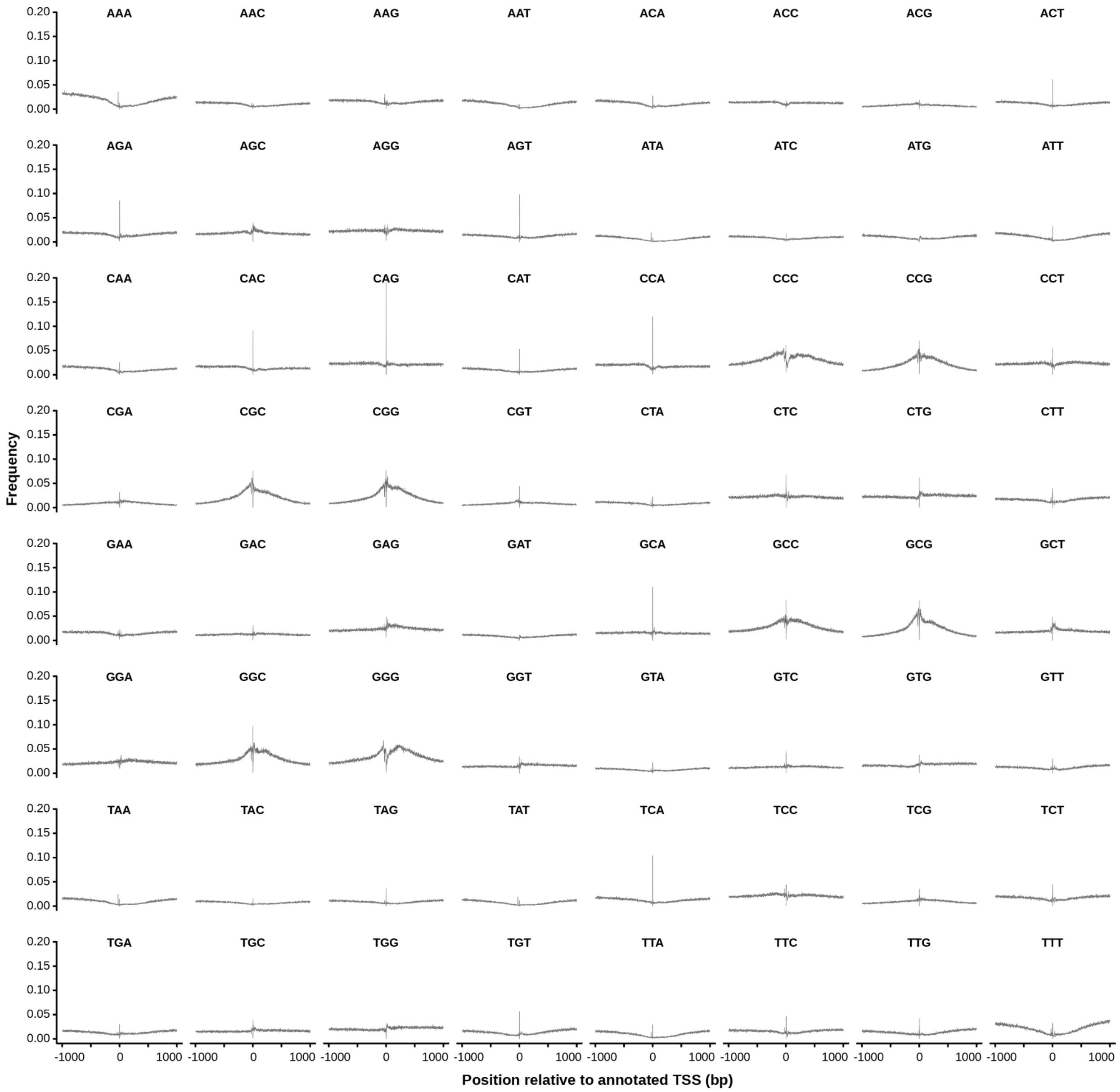


**Fig. S9. Trinucleotide frequencies at each basepair position near TSS.** Frequencies of each trinucleotide (y-axis) at every basepair position relative to TSS (x-axis), across the top 40,000 TSS ranked by FANTOM CAGE signal.

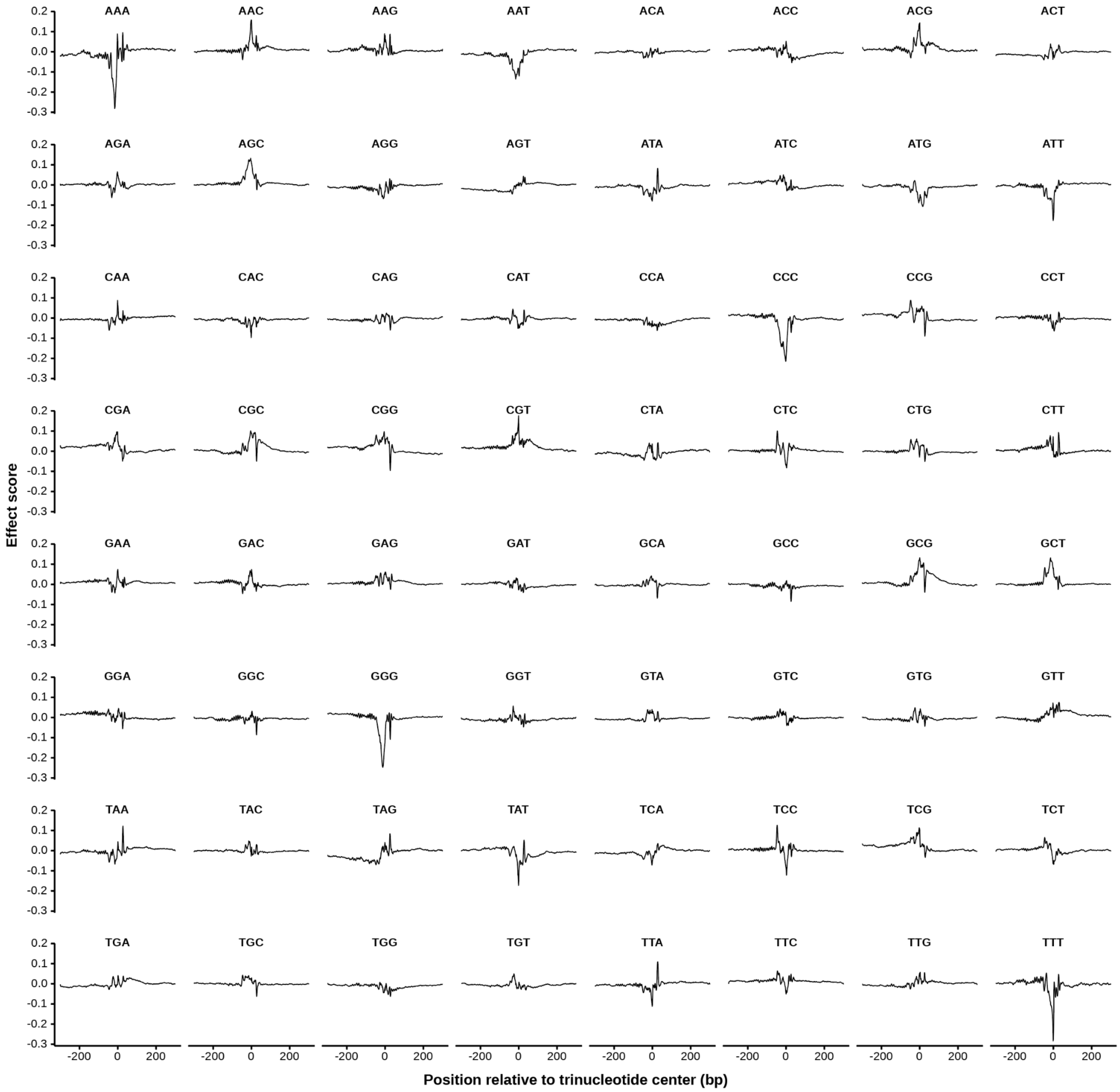


**Fig. S10. Position-specific effects of trinucleotides.** The normalized position-specific effect curves of each trinucleotide are shown.

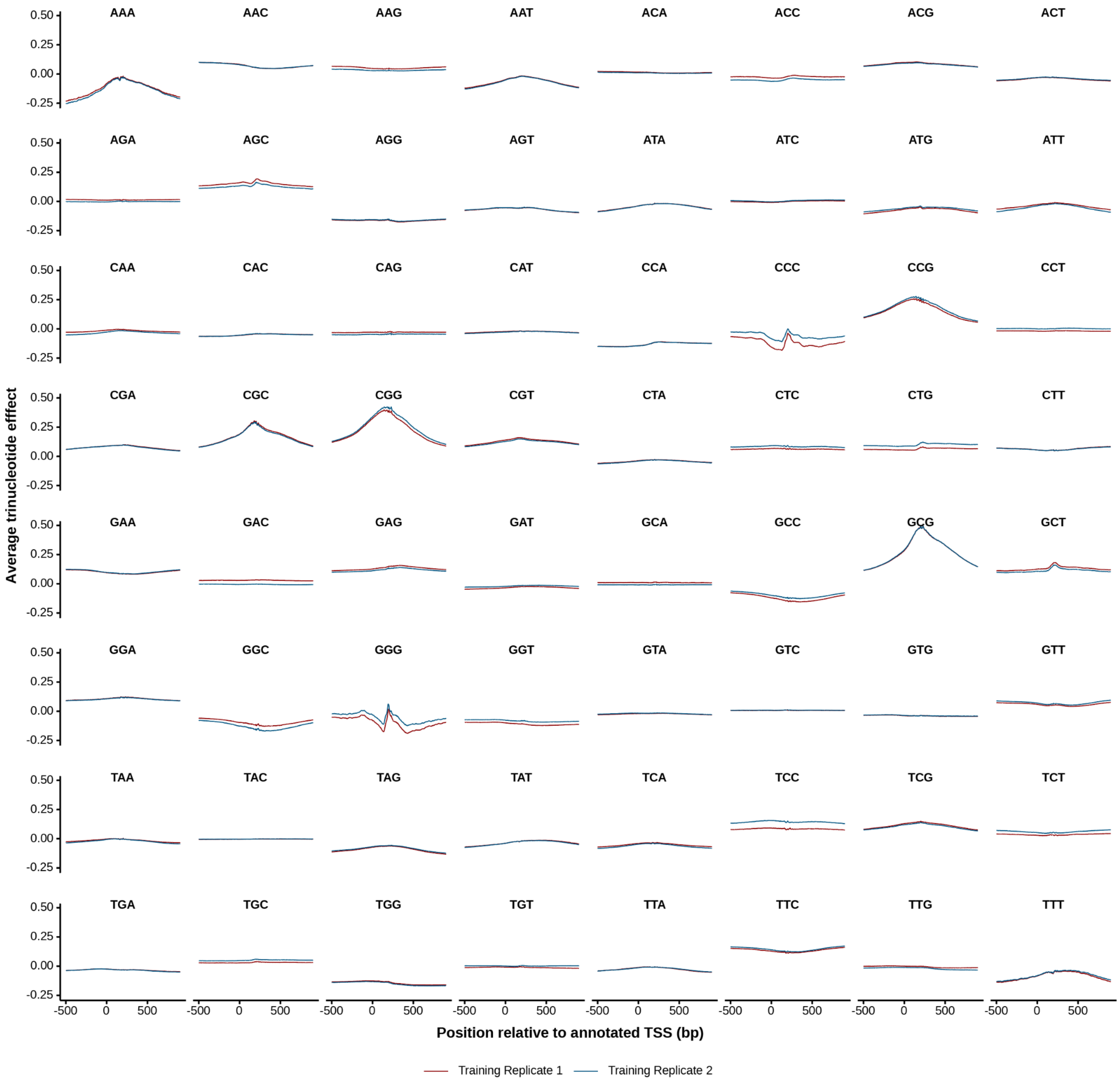


**Fig. S11. Comparison of average trinucleotide contribution to TSS between training replicates for each trinucleotide.** For each training replicate, average trinucleotide effects for each of the 64 trinucleotides across the top 40,000 TSS ranked by FANTOM CAGE signals were computed. Training replicates were indicated by color.

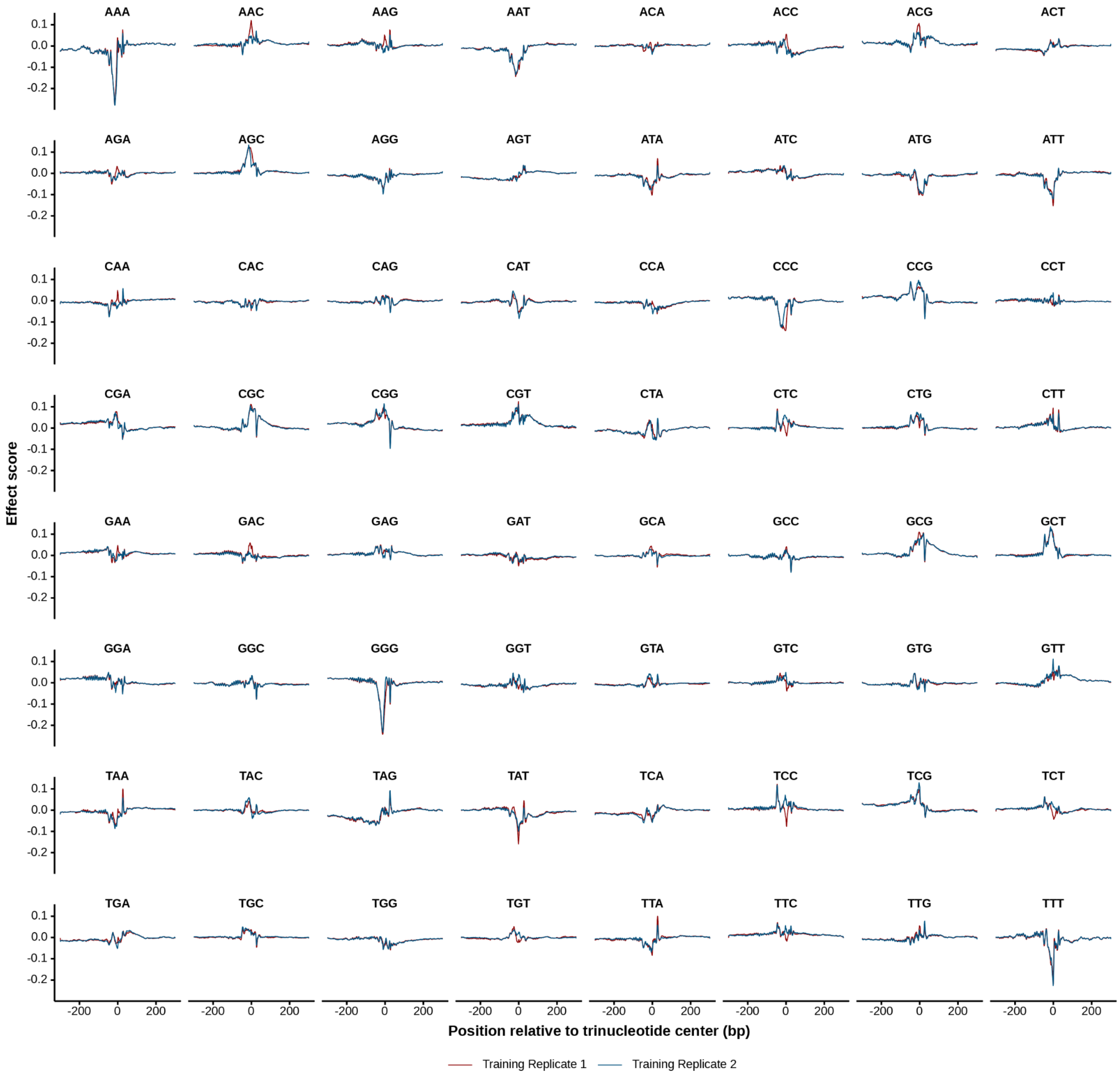


**Fig. S12. Comparison of position-specific effects of trinucleotides between training replicates.** The normalized position-specific effect curve of each trinucleotide is shown for each training replicate. Training replicates were indicated by color.

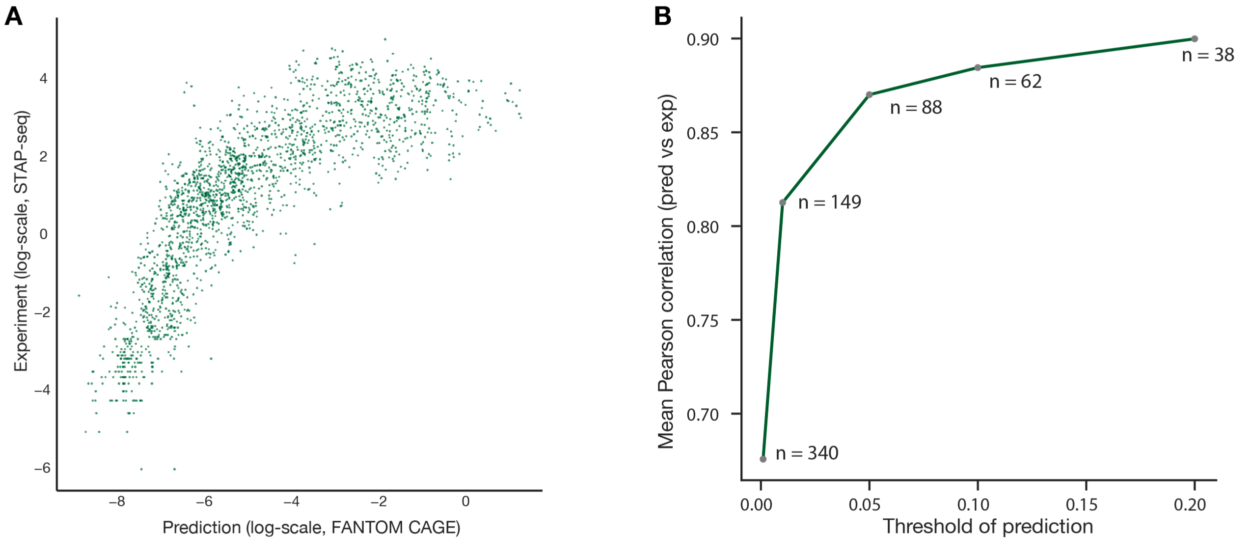


**Fig. S13. Quantitative comparison of Puffin prediction with experimental measurements of TATA and NFY motif perturbation experiments.** (**A**). The predicted expression level per sequence for FANTOM CAGE by Puffin (x-axis) is highly correlated with the experimental measurement with STAP seq (y-axis). (**B**). Basepair-level correlation between prediction and experimental measurement (y-axis) is high for sequences predicted with high expression level (x-axis).

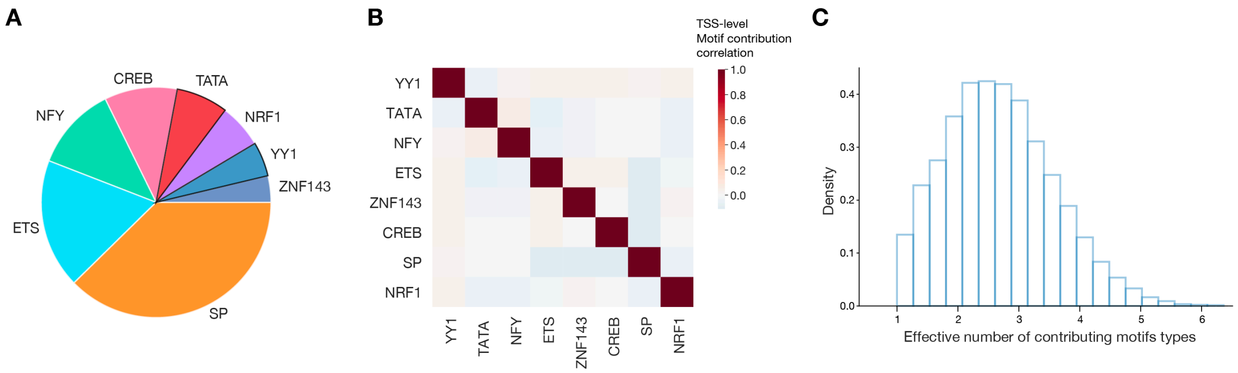


**Fig. S14. Summary statistics of motif contribution scores across human TSS.** (**A**) Proportions of motif contribution by motif types across the top 40,000 TSS ranked by FANTOM CAGE signal. Dark borders indicate directional motifs. (**B**). Pairwise correlations between motif contribution score for each motif type. For directional motifs TATA and YY1, only forward-direction motifs were considered. For bidirectional motifs the forward and reverse directional motif contributions were added. (**C**) The distribution of the effective number of contributing motif types (among YY1, TATA, SP, NFY, ETS, ZNF143, CREB, and NRF1) per TSS. The effective number is estimated by $2^{\text{entropy(bits)}}$. Entropy scores for contributing motif types were computed from motif contribution scores (for all motif types shown in **A** and **B**).

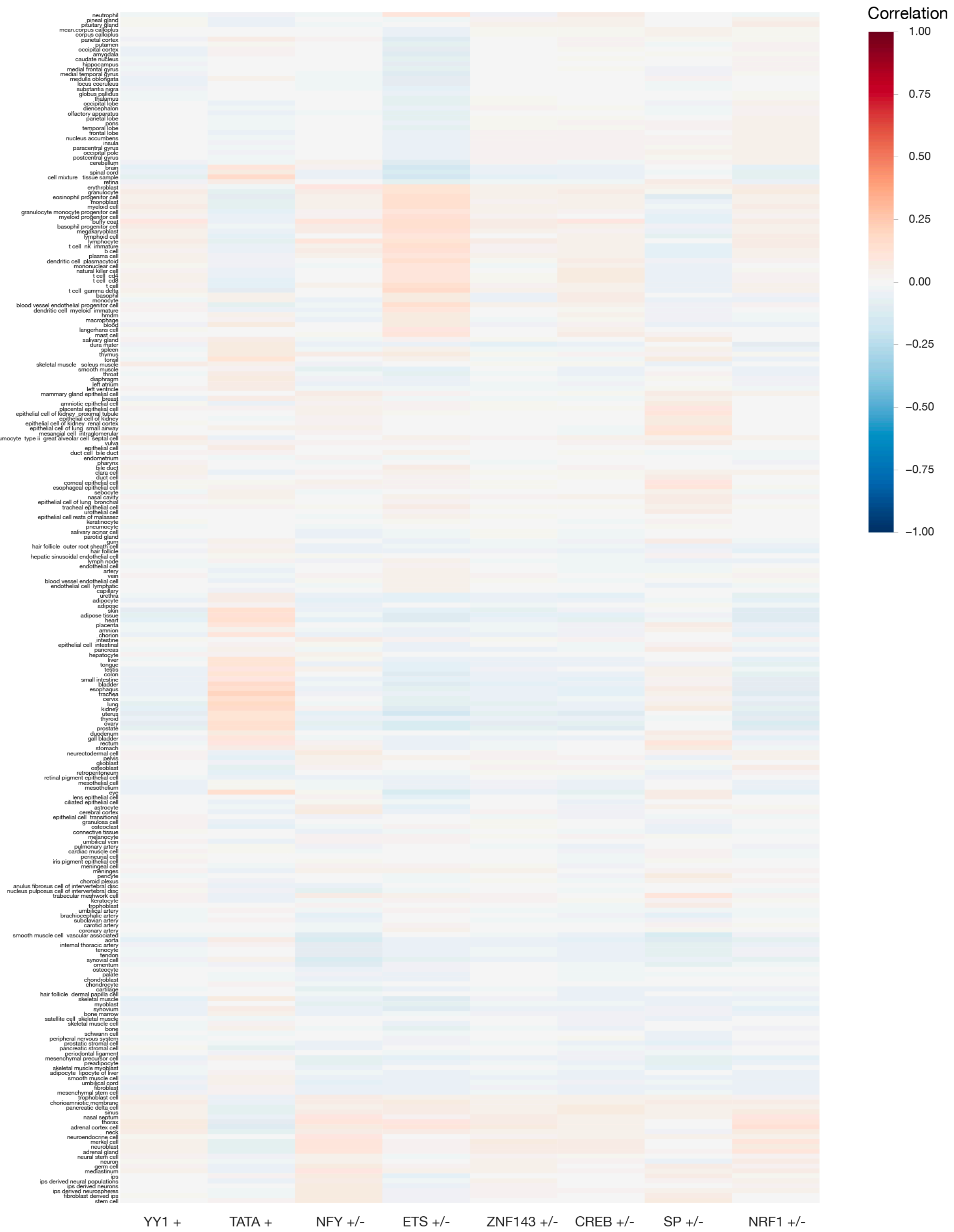


**Fig. S15. Correlation between motif contribution scores and cell type / tissue-specific gene expression.** Heatmap visualizes the correlations between motif contribution scores and cell type / tissue-specific expression values (log fold over average expression across > 200 cell type / tissues from the FANTOM project) across TSS.

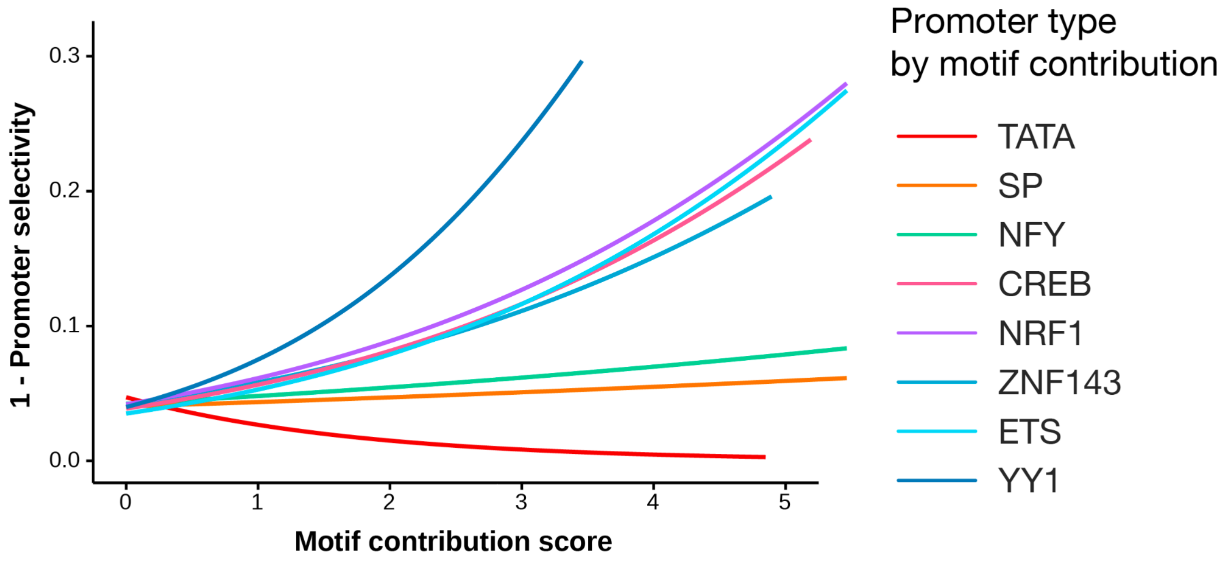


**Fig. S16. Motif contribution score predicts the selectivity of promoter from in silico insertion screen.** GAM-fitted curves of 1 - selectivity score (y-axis) versus motif contribution score (x-axis) of each motif type (color) across all TSS.

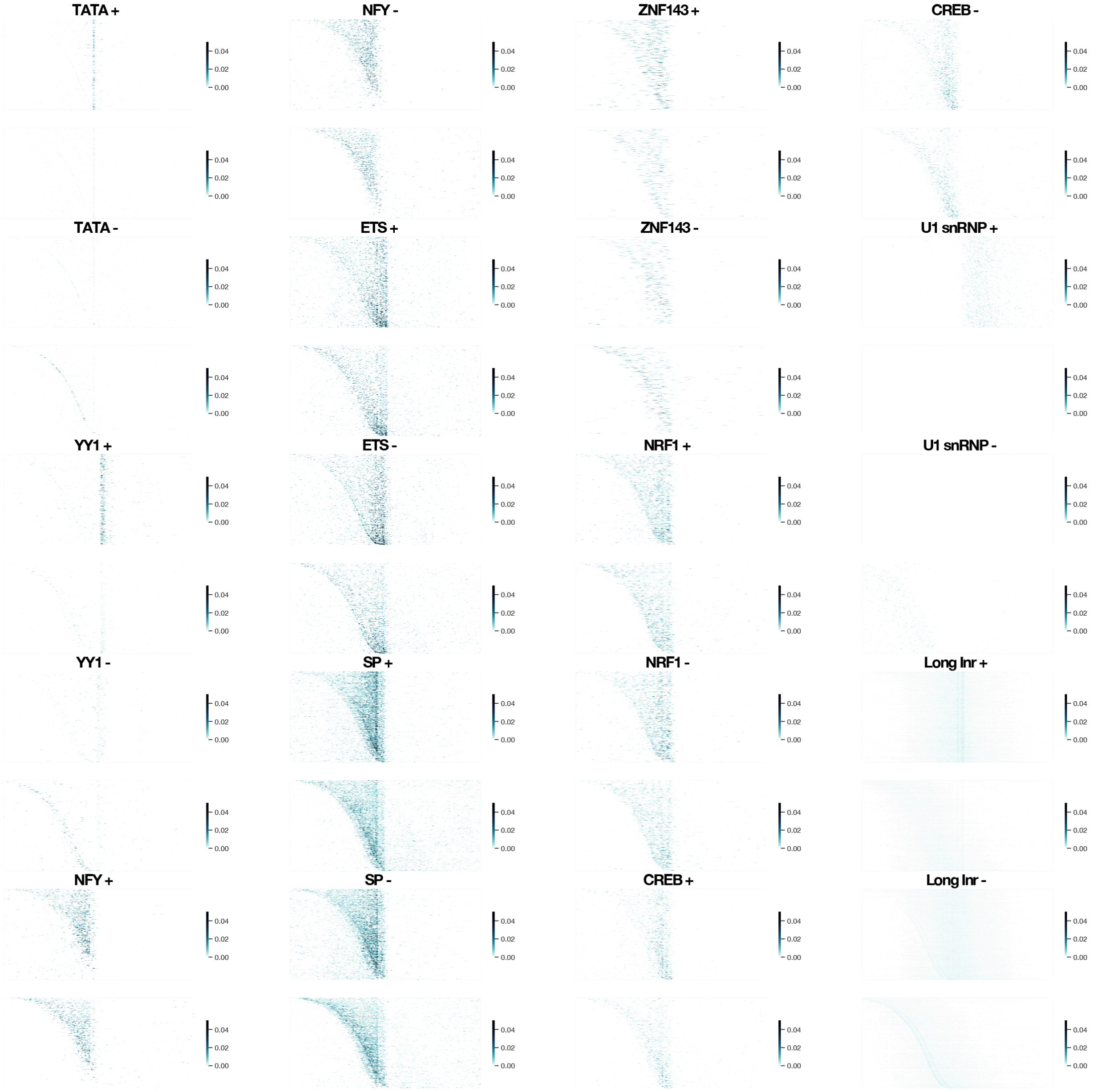


**Fig. S17.** Per-motif basepair contribution scores for forward (top) and reverse (bottom) strand transcription of 8,216 promoters, sorted by reverse TSS position, scaled by the sum of positive basepair contribution scores on both strands. The per-motif basepair contribution scores shown were computed for PRO-cap. The matrices shown in heatmaps were smoothed with a small rectangular filter of size 10x1.

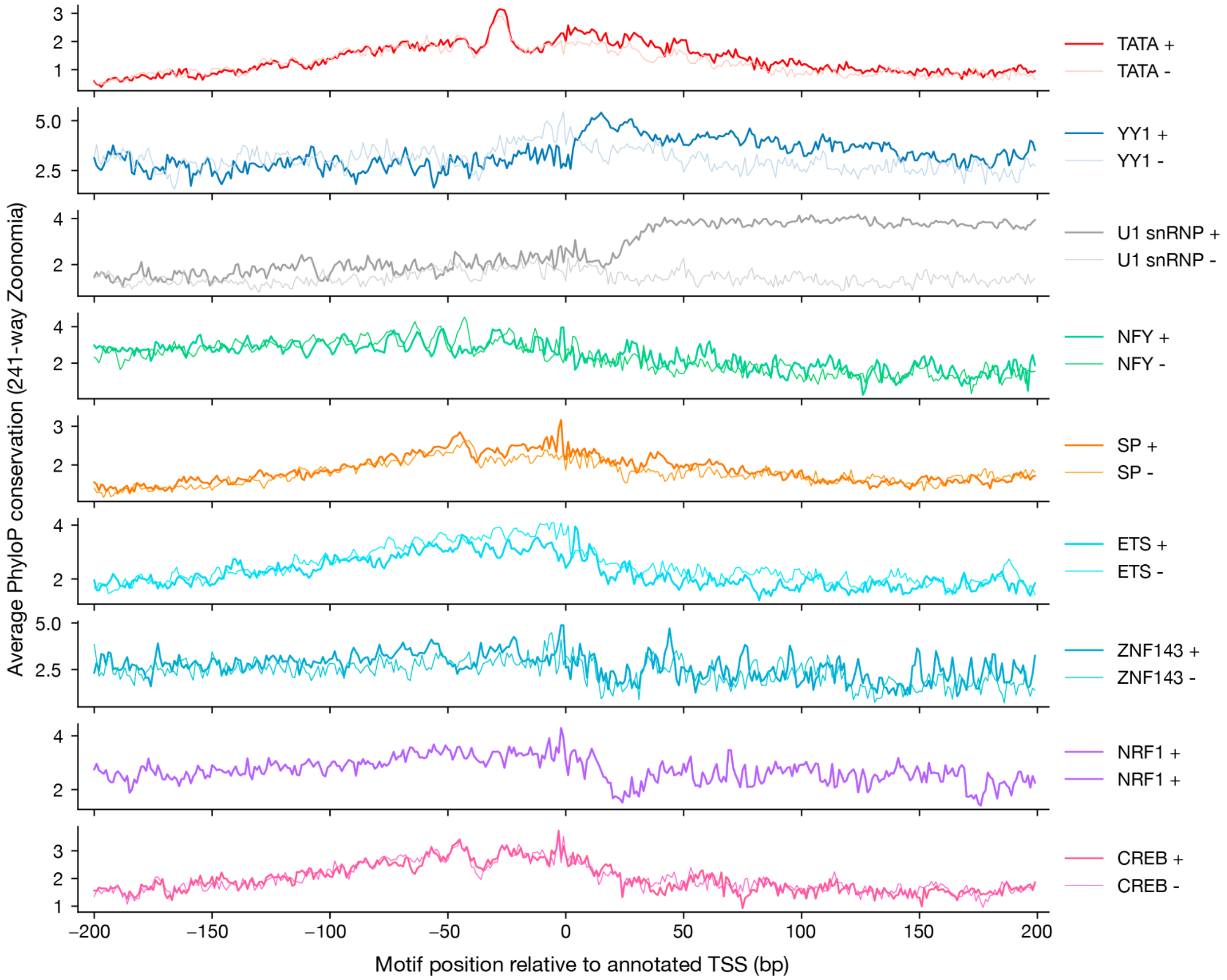


**Fig. S18. Position-specific sequence conservation scores for all motifs**. X-axis shows the position (bp) relative to annotated TSS (human genome viewpoint) and the y-axis shows average PhyloP scores for each motif, computed with the average weighting by motif activation scores. Note that the x-axis shows TSS-centric coordinates, which should be reversed when comparing with position-specific motif effects.

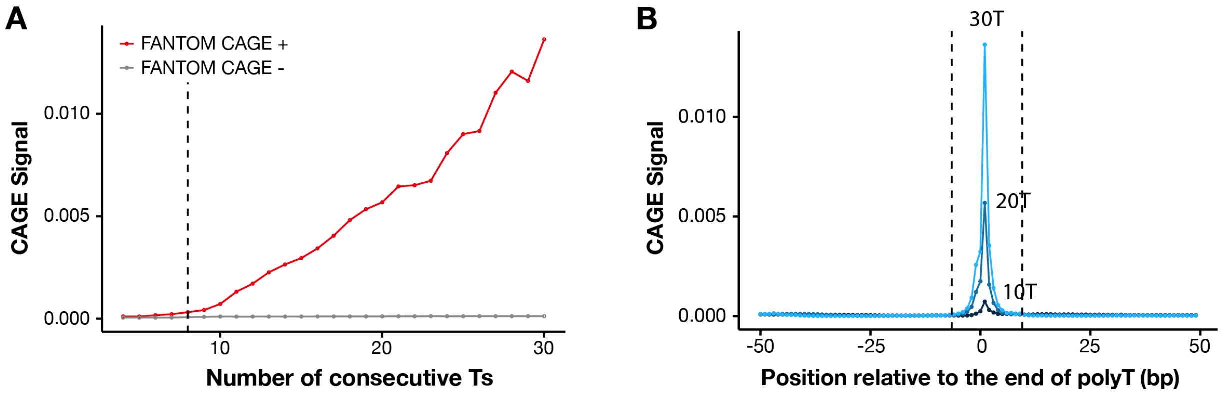


**Fig. S19. Analysis for correction of FANTOM CAGE-specific bias for consecutive Ts.** (**A**) Increase of uncorrected FANTOM CAGE signal (y-axis) for different numbers of consecutive Ts (x-axis). The uncorrected FANTOM CAGE signals were aggregated by the average signal, per position, relative to the last T of each poly-T, as shown in (**B**) which shows the position-specific signal for 10T, 20T, and 30T. The y-axis in panel (**A**) shows the maximum average signal across all positions for each number of consecutive Ts. The dashed line in (**A**) shows the threshold for correction (>=8T), and the dashed lines in (**B**) show the interval for correction, [-6, +10), relative to the last T of each poly-T.

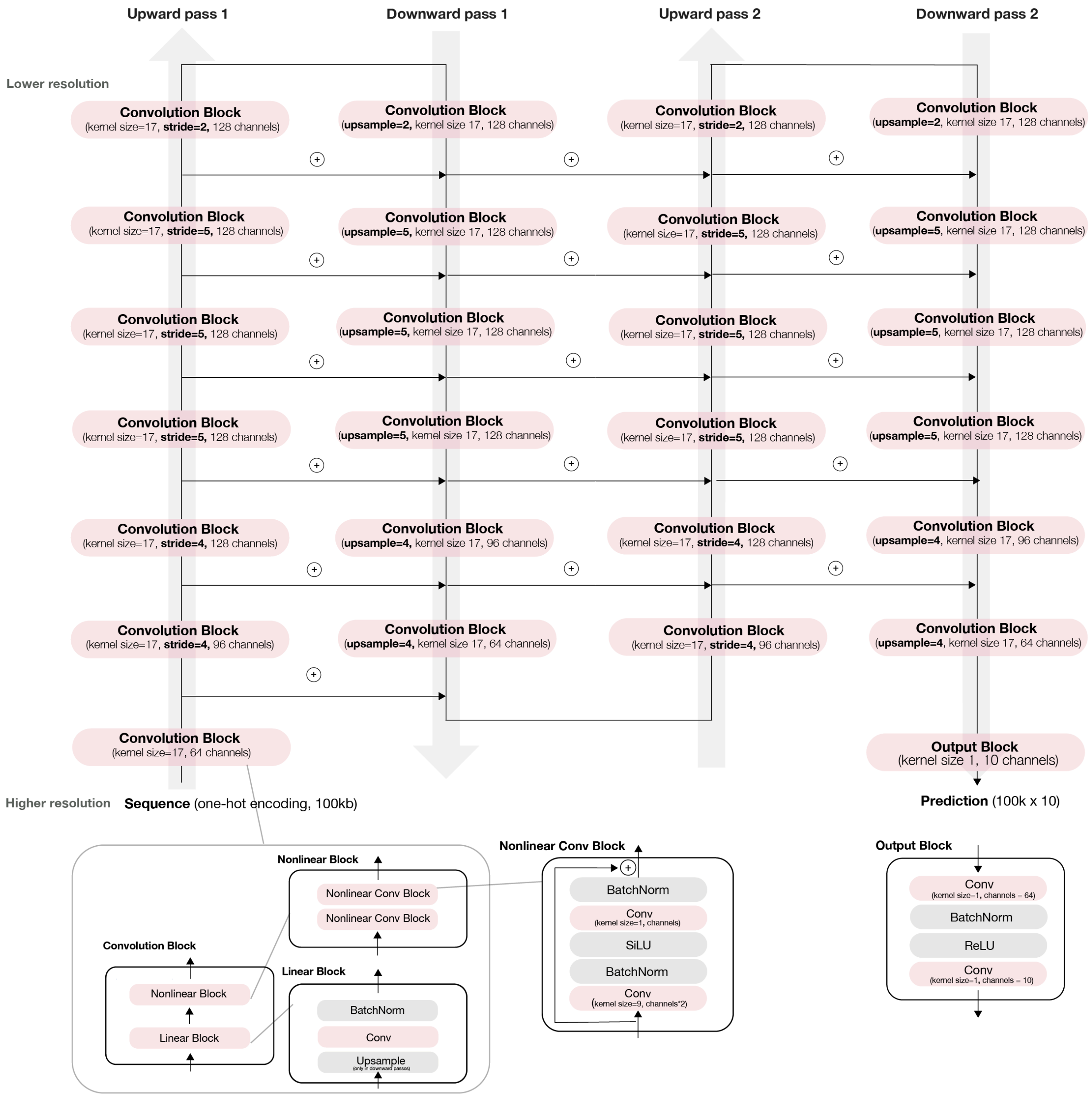


**Fig. S20. Schematic illustration of Puffin-D model architecture.** The architecture contains two upward and downward passes with residual connections. The spatial dimension downsampling and upsampling were implemented with strided convolution layers and upsampling layers. Each block is illustrated in more detail with a dedicated diagram. Layer parameters are the same as the block parameters unless otherwise indicated.

**Supplementary Text 1. Deep learning sequence model-inspired design of Puffin model**

The design of the Puffin model is inspired by our analyses of the deep learning sequence model Puffin-D, which was also developed for this manuscript. These two models are highly complementary and allowed us to have access to both the higher prediction performance and longer input context sequence size of the Puffin-D and the simple and interpretable Puffin model which has led to findings that would not have been possible without it.

The main approach for analyzing Puffin-D was virtual genetic screens, where we designed input sequences to Puffin-D, and analyzed the output to infer properties of the sequence dependencies captured by the model. Here we demonstrate a representative analysis, which showed two important aspects of the Puffin model design: strand- and position-specific effects of motifs and the lack of strong spatial interactions among motifs beyond the additive / multiplicative combination of their effects. In this analysis, we generated synthetic TSS sequences using random combinations of motifs identified from Puffin-D in silico mutagenesis results. Here we used Puffin motifs which produce similar results for consistency.

More specifically, we generated 10 million random synthetic TSS, each of them containing 3 motifs randomly selected from a list of motifs in both forward and reverse directions, inserted in random positions within an approximately 300bp window (the window size slightly varies due to motif length differences). The motif list also includes a placeholder 0bp "empty" motif as a negative control. The background sequence before motif insertion is randomly selected from neutral genomic sequences with no TSS activity from a 16Mb region on chr8, a holdout chromosome during training.

We then selected sequences with top TSS activities as predicted by Puffin-D on both forward and reverse strands separately and analyzed their motif positional distribution relative to the max predicted transcription initiation signal position. We observe that motifs have position-specific effects, most distinctively for TATA and YY1 (Supplementary Text Fig. 1). Moreover, some motifs have asymmetric or directional positional distribution between forward and reverse strand TSS, while other motifs are symmetrical (Supplementary Text Fig. 1). Such in silico experiments on Puffin-D also suggested the effect ranges of motifs. Therefore the strand-specific and position-specific effect is an important part of the simple model design.

Next, we assessed spatial positioning constraints between motifs captured by Puffin-D, in which we plotted pairwise positional distribution between every pair of TSS for synthetic sequences with top-predicted TSS activities. We can observe no strong constraints between pairs of motifs and motif positions appear largely independent from each other (Supplementary Text Fig. 2; the top-left and bottom-right corners of each scatter plot are forbidden regions due to the window size constraint rather than spatial interactions). This inspires a simple model design without explicitly modeling spatial interactions between motifs beyond the additive / multiplicative combination of motif effects, which also allows for simpler interpretation.

The initial Puffin model design contains only a single sequence pattern type, and we discovered that there exist two types of sequence patterns from the models learned, one that is long in size and has long effects range and the other is short in size and has local effects within the sequence pattern. Thus, we created a model design with only motif and initiator sequence pattern types, similar to the stage 1 model architecture. We then seek to simplify the model architecture, especially the motif sequence patterns to increase interpretability. After choosing 10 consensus motifs (>0.95 maximum cross-correlations across >7 replicates) and removing other motifs, we discovered that continued training always alters some motif sequence patterns that we initialize it with, which leads to the hypothesis that there exists another important sequence pattern type that is not included. Then we discovered that introducing the simple trinucleotide sequence pattern is sufficient to ensure that the motif sequence patterns are stable over continued training, which lead to the final Puffin model design.

More generally, we also propose these steps as a generally applicable approach to obtain a better understanding of the underlying biological problem through machine learning modeling: first, training a black-box deep learning model that aims to achieve the best performance; second, design and perform virtual genetic screens to understand the model; third, design simple interpretable models for the problem. All steps are iterative and can combine information learned from the later steps to improve the previous steps.

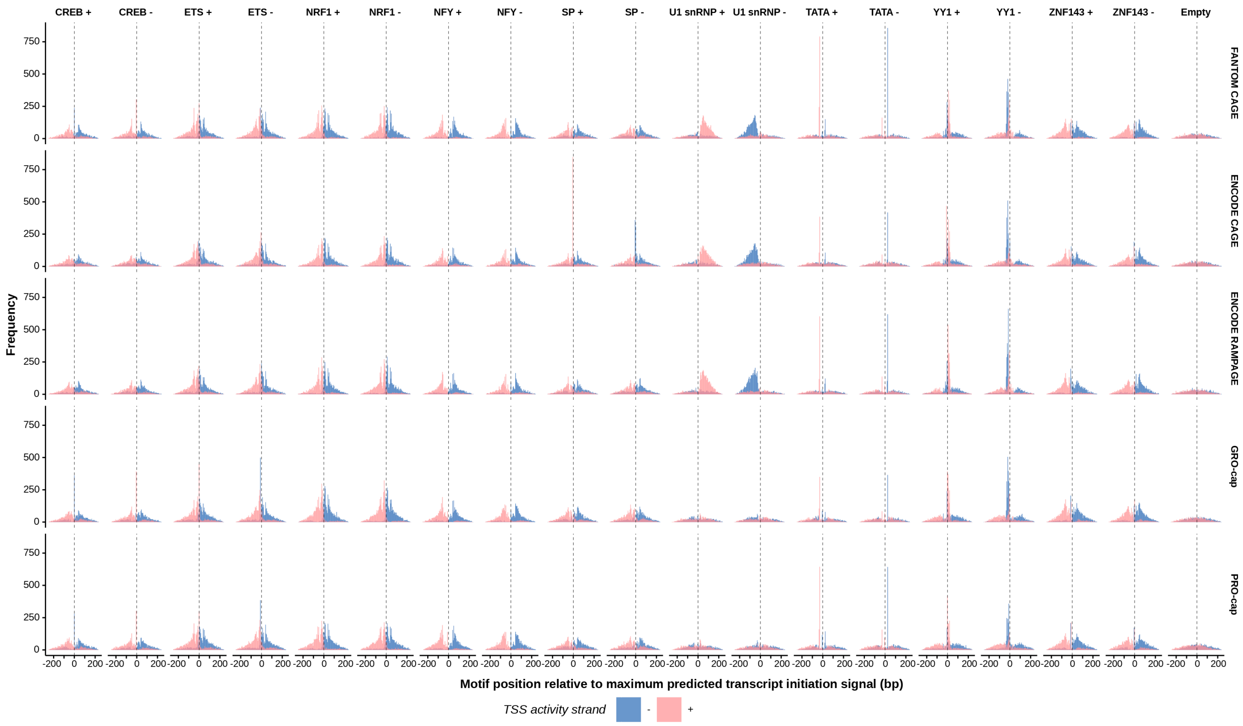


**Supplementary Text Fig. 1. Motif positional distribution for synthetic TSS with top 0.1% predicted activity by Puffin-D.** The positional distribution of each motif (bp relative to the position with the maximum predicted transcription initiation signal) is shown. Positional distribution relative to synthetic TSS on different strands (with maximum activity on forward versus reverse strand) was indicated by different colors.

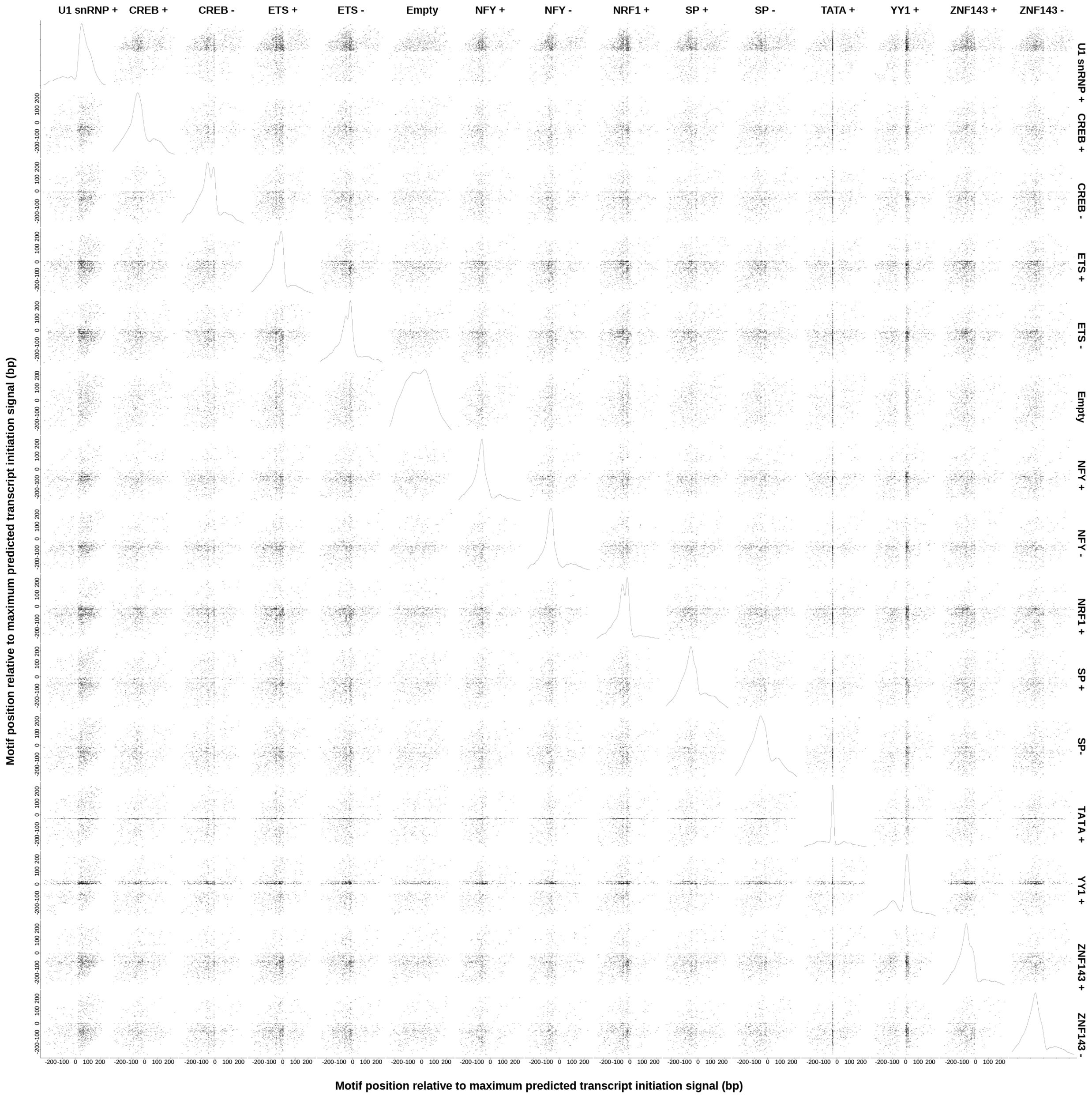


**Supplementary Text Fig. 2. Pairwise motif positional distribution for synthetic TSS with top 0.1% predicted activity by Puffin-D.** The positions of each motif pair were relative to the position with the maximum predicted transcription initiation signal. The top 0.1% synthetic TSS with activity on the forward strand was shown.

**Supplementary Text 2. Interpretation and normalization of trinucleotide effects learned in a machine learning model.**

Our first step of deriving and interpreting trinucleotide effects is converting the 32 trinucleotide sequence pattern weight scores and 64 sequence pattern position-specific effect curves to position-specific effect curves of each individual trinucleotide (AAA, AAC, AAG, AAT, ACA, …, TTT). This can be computed by first computing the 32 trinucleotide sequence pattern activations from an individual trinucleotide and then computing the sum of all trinucleotide sequence pattern effects from the activations. Looking up the trinucleotide position-specific effect scores for each trinucleotide is an equivalent model to the original formulation, and we consider this formulation here for simplicity.

It is important to note that the raw trinucleotide effect scores are still not yet ready for interpretation because these scores are derived from an under-constrained optimization problem. This can be readily shown from the fact that training replicates with different random initializations lead to very different trinucleotide sequence pattern weights, effects, and raw trinucleotide effects, while the total trinucleotide effects remain highly similar. In other words, for any fixed input, there are many drastically different sets of parameters that can lead to the same output. This is in contrast to learning motifs, for which the same set of motifs is reproducibly identified across training replicates. We also note that this issue cannot be resolved adequately by standard regularizations during training.

To intuitively describe the under-constrained nature of the learning problem, one can see that adding or subtracting the same position-specific effects to all trinucleotides does not affect the model output (to be more precise, it does not affect model output up a constant, which can be easily canceled out by bias terms in the model). Moreover, adding the same position-specific effects to all trinucleotide patterns matching A** where * matches any bases, and subtract the same effects to all trinucleotide patterns matching *A* (or **A) after shift by 1 (or 2 bp for **A) also do not change the model output. Similarly, operations can be done to patterns such as AA* and *AA. Obviously, such a lack of constraint limits the interpretability of the raw model weights or trinucleotide effect scores.

To address this issue and ensure the reproducibility and interpretability of trinucleotide position-specific effects, we impose additional constraints by applying normalization of raw trinucleotide effect scores, without changing the model output. The additional constraints we added are: 1. the sum of position-specific effect curves for all trinucleotides is 0 everywhere. 2. Sum of position-specific effect curves for trinucleotide matching the same mononucleotide patterns are the same regardless of their position in the trinucleotide. For example, the sum of effects for A** patterns is the same as *A* or **A. 3. Sum of position-specific effect curves for trinucleotide matching the same dinucleotide patterns are the same regardless of its position in the trinucleotide. For example, the sum of effects for AA* patterns is the same as *AA.

We designed a simple iteration correction algorithm to normalize the raw trinucleotide effect curves to satisfy such constraints without changing its output (up to a constant), detailed in our GitHub repository https://github.com/jzhoulab/puffin_manuscript. The normalized trinucleotide effect curves are highly reproducible across training replicates and can be intuitively interpreted. We expect this issue and solution to be also generally applicable to the interpretation of k-mer effects learned from machine learning models.

**Supplementary Data 1.**
List of transcription initiation signal profiles used in this manuscript. The URLs or accession numbers are provided.

**Supplementary Data 2.**
Consensus motifs and motif effects from Puffin training replicates. For each consensus motif, all instances of the motif identified from all training replicates were shown.

**Supplementary Data 3.**
Consensus motifs and motif effects from Puffin training replicates on mouse data. For each consensus motif, all instances of the motif identified from all training replicates were shown.

**Table S1.**
List of individually identifiable sequence patterns with name and IDs by Puffin

**Table S2.**
Motif contribution scores of human transcription start sites.

**Table S3.**
In silico insertion screen selectivity scores and motif selectivty scores.

**Table S4.**
Transcription start site annotations used in this manuscript.
